# Sphingolipid Homeostasis, Mitochondrial Activity, and PKA Signaling Drive an Azole-Tolerant State

**DOI:** 10.1101/2025.09.12.675915

**Authors:** Cécile Gautier, Eli I. Maciel, Corinne Maufrais, Naomi Lyons, Nora Kawar, Chen Bibi, Caleb Eickmann, Alex Rosenberg, Deveney Dasilva, Nathalia Fidelis Viera De Sa, Maurizio Del Poeta, Richard J. Bennett, Judith Berman, Iuliana V. Ene

## Abstract

Antifungal therapies frequently fail, resulting in persistent infections caused by the important opportunistic pathogen, *Candida albicans*, which is highly tolerant to azole drugs. Tolerance arises from a subpopulation of cells that survive prolonged drug exposure, yet the underlying mechanisms remain poorly defined. Here, transcription factor mutant library screens identified Mnl1 as a key repressor of tolerance. Loss of *MNL1* elevated cAMP levels and activated the PKA pathway, linking Mnl1 to growth control and stress responses. Additionally, *mnl1* cells exhibited altered sphingolipid composition, implicating Mnl1 in regulating membrane permeability, a key determinant of drug efflux and membrane integrity. Increased tolerance in *mnl1* cells was also linked to increased mitochondrial function. Thus, Mnl1 regulates tolerance through coordinated control of sphingolipid metabolism, mitochondrial activity, and PKA signaling, highlighting that tolerant cells adopt a distinct physiological state primed to withstand antifungal stress.

## INTRODUCTION

*C. albicans* is a prevalent human fungal pathogen and a significant cause of morbidity and mortality, particularly in immunocompromised individuals. This opportunistic pathogen can cause both superficial and life-threatening invasive infections, with an associated mortality of 27-60%, despite antifungal treatment^1^. The cornerstone of therapy for *C. albicans* infections is azole antifungals, particularly fluconazole (FLC), which target ergosterol biosynthesis by inhibiting the enzyme lanosterol 14α-demethylase (Erg11)^2,3^. Depleting ergosterol, an essential component of the fungal cell membrane, compromises membrane fluidity, leading to growth arrest and cell death^2^. *C. albicans* strains that are resistant or tolerant to azole antifungals are more likely to cause antifungal treatment failures^4,5^. Azole resistance is frequently associated with mutations causing increased expression of drug efflux pumps (e.g., Cdr1, Cdr2) or the target enzyme Erg11, or with altered Erg11-azole binding affinities^6–8^. Surprisingly, for *C. albicans*, the high rates of therapeutic failure do not align with the low prevalence of resistance detected in clinical susceptibility tests^9,10^. In addition, azole tolerance, defined as the ability of a subpopulation of cells to grow at high drug concentrations, has emerged as a potential concern^10,11^. Tolerance enables fungal cells to withstand supraMIC drug concentrations and is linked to infection persistence and higher mortality rates in candidemia patients^9–12^.

The molecular mechanisms underlying azole tolerance in *C. albicans* are poorly understood. Tolerance has been linked to stress responses that influence cell wall and plasma membrane integrity^10^. Thus, reduced tolerance was detected in strains affecting calcineurin pathway components and regulators (Cnb1, Crz1, Rcn1, and Rcn2), the cell wall maintenance mitogen-activated protein kinase Mkc1, the ergosterol biosynthesis regulator Upc2, the iron utilization regulator Iro1, and the vacuolar trafficking protein Vps21^11^. Hsp90, another key mediator of azole tolerance and a regulator of protein homeostasis, stabilizes several stress response regulators, including calcineurin and the Mkc1 pathway^11,13^. Tolerance also was increased by overexpression of transcription factors Crz1 and Gzf3; the deletion of *CRZ1*, but not *GZF3*, reduced tolerance; however, the precise roles of these factors in tolerance remain unclear^14^. The involvement of protein homeostasis via the ubiquitin-proteasome system is consistent with the observation that Rpn4 promotes tolerance by regulating protein homeostasis and by upregulating efflux pumps that reduce intracellular drug levels^15,16^. Tolerance varies with pH and with strain genetic backgrounds^11^, suggesting that multiple mechanisms are at play. Pharmacologically, tolerance can be abrogated by inhibitors of Hsp90 (geldanamycin, radicicol), calcineurin (cyclosporine A, FK506), Pkc1 (staurosporine), mTOR (rapamycin), sphingolipid biosynthesis (aureobasidin A), N-linked glycosylation (tunicamycin), and other agents like fluphenazine, which affect ABC transporters and calcineurin^11,17,18^. This supports the idea that multiple regulatory hubs mediate azole tolerance. Nonetheless, the molecular consequences of tolerance and how it integrates with broader stress response networks remain poorly defined.

The contribution of the cyclic AMP (cAMP)-dependent protein kinase A (PKA) pathway to tolerance has yet to be investigated. The PKA pathway regulates several processes, including stress responses, morphogenesis, metabolism, virulence, and cell death^19–21^. Initiated by ligand binding to a G-protein-coupled receptor, the PKA pathway activates adenylyl cyclase, thereby increasing cAMP levels. The binding of cAMP to the Bcy1 regulatory subunit releases and activates the Tpk1 and Tpk2 catalytic subunits, which phosphorylate multiple, partially overlapping, downstream targets^22,23^. In *C. albicans,* the PKA pathway affects azole susceptibility by regulating ergosterol biosynthesis, biofilm formation, and stress responses^24^. Cells lacking Ras1 or Cyr1 are hypersensitive to azoles, and those lacking the cAMP phosphodiesterase Pde2 have increased ergosterol and altered FLC responses^24–26^. PKA also controls biofilm-associated resistance^22,24,26^ and, in *C. auris*, regulates the expression of multidrug transporters^23^. By affecting intracellular drug accumulation and stress-induced cell death, PKA fine-tunes the balance between growth and survival under antifungal stress^19^.

In this study, we screened for novel transcriptional regulators of azole responses in *C. albicans*, identifying factors involved in resistance and/or tolerance, with a focus on those specifically affecting tolerance. We identified a network of 31 transcription factors that regulate azole tolerance, structured into two distinct regulatory modules: a stress response module and an unstressed log-phase growth module. Importantly, Mnl1, a regulator of the weak acid stress response^27^, was identified as a major repressor of FLC tolerance. *MNL1* deletion upregulates ATP biosynthesis genes and elevates cAMP levels, linking Mnl1 to mitochondrial function and the PKA pathway. Loss of *MNL1* also disrupted sphingolipid homeostasis, consistent with the observation that alterations in sphingolipid metabolism can compensate for reduced membrane ergosterol^28–30^. Taken together, these findings suggest that Mnl1 coordinates azole tolerance by regulating PKA signaling, mitochondrial activity, and lipid metabolism.

## METHODS

### Strains and growth conditions

*C. albicans* and *S. cerevisiae* strains used in this study are listed in **Supplementary Table 1**. Cells were cultured overnight in 3 mL of liquid YPD (2% bacto-peptone, 1% yeast extract, 2% dextrose (filter-sterilized)) at 30°C with shaking (200 rpm). Cell densities were measured using optical densities of culture dilutions (OD_600_) in sterile water using a Biotek Epoch 2 plate reader.

### Growth assays

For growth curves, *C. albicans* cells were seeded in 96-well plates at a concentration of 2 × 10^5^ cells/mL and grown in YPD at 30°C with orbital shaking for 24 h. Cell densities were measured every 15 min (OD_600_) using a Biotek Epoch 2 plate reader, and doubling times were calculated using the exponential growth phase of each curve. Measurements of lag phase duration were determined using the BioTek Gen5 software. Growth curves were performed with three biological replicates, each with technical duplicates.

### Construction of the Mnl1-Neon strain

To generate *C. albicans* strains with a tagged protein, we amplified *NAT1* from plasmid pJB-T155^31^ using primers 2313 and 2251, mNeonGreen from plasmid pJB-T409^32^ using primers 2315 and 2369. The plasmid backbone containing an *E. coli* origin of replication, and the Amp^R^ gene from plasmid pJB-T155 were amplified using primers 2325 and 2027. The promoter sequence of *TDH3* was amplified from the genomic DNA of *C. albicans* SC5314 with primers 2239 and 2314. The four template plasmids were digested with the Dpn1 restriction enzyme (at 37°C for 16 h), and the four linear amplified PCR products were purified using a GeneJET PCR Purification Kit (K0702, Thermo Scientific). For Gibson assembly, 0.3 pmols of each of the four fragments were adjusted to 10 μl volume in double distilled water. The reaction was initiated by adding 10 μl of HiFi DNA Assembly Master Mix (NEB-E5510S, Ornat) and incubating the reaction at 50°C for 2 h. The plasmid expression cassette (*NAT1-TDH3-mNeonGreen*) was amplified from pJB-T510 by PCR using PCRBIO VeriFi Mix (Rhenium Bio) with primers complementary to 40 bp 5’-to the start codon of the gene ORF, and 40 bp downstream of the start codon. The amplified PCR product was transformed into a fresh *C. albicans* SC5314 log phase liquid culture and transformants were selected on YPD plates supplemented with nourseothricin (400 µg/ml). Successful transformants were validated by diagnostic PCR using PhireGreen Hot Start II PCR Master Mix (F126L, Thermo Fisher Scientific) along with a primer specific for the expression cassette (primer 2039) and a primer carrying homology to the target gene. Plasmid and primer sequences are provided in **Supplementary Table 2**.

### Construction of the *pde2, rme1,* and *hcm1* deletion mutants

*C. albicans* SC5314 was transformed using the transient CRISPR/Cas9 system and the plasmid pV1093, as previously described^33^. This plasmid contains an sgRNA scaffold with the SNR52 promoter and terminator. PCR was used to generate the synthetic guide RNAs (sgRNAs) targeting *PDE2, RME1,* or *HCM1*. In these specific transformations, distinct sgRNAs specific to each gene were constructed and used. Additionally, pV1093 contains a NAT resistance cassette (encoded by the *SAT1* gene) and a CAS9 expression cassette, both of which can be amplified via PCR. All reactions used to construct the CRISPR components were cleaned and concentrated through ethanol precipitation, except for the second PCR for the synthetic guide RNA.

#### PCR-based Construction of CRISPR Components

##### Synthetic Guide RNA Construction

The synthetic guide RNA was constructed through three rounds of PCR. In the first PCR, the SNR52 promoter was amplified with a guide sequence using oligos #1 and #2. Simultaneously, the guide sequence was amplified with the scaffold and terminator using oligos #3 and #4, with pV1093 as the template. The products from the first PCR were combined and used in a second PCR to extend and complete the synthetic guide RNA. The second PCR product was used as the template for the third PCR, and nested oligos #5 and #6 were used to amplify the final sgRNA expression cassette.

##### Repair Template Construction

The repair template was generated through a single PCR. The repair template consists of a 96 bp forward oligo upstream of the gene of interest with an overhang to the NAT cassette and a 96 bp reverse oligo downstream of the gene of interest, also with an overhang to the NAT cassette. This PCR amplification of the repair template was successfully performed with DreamTaq polymerase.

##### CAS9 Cassette Construction

The CAS9 cassette was amplified through a single PCR using oligos #7 and #8 with pV1093 as the template.

*C. albicans* strain SC5314 was transformed via the lithium acetate method^34^. The following DNA amounts were used for the *PDE2* transformation: 3 μg of each sgRNA, 5 μg of the CAS9 cassette, and 5 μg of the repair template. The following DNA amounts were used for the *HCM1 and RME1* transformations: 5 μg of each sgRNA, 9 μg of the CAS9 cassette, and 8 μg of the repair template. Successful transformants were selected on YPD plates supplemented with 200 μg/mL nourseothricin after 2 days of incubation at 30°C. Individual colonies were resuspended in 100 μL of 20 mM NaOH and boiled at 99°C for 20 min, and 2 μL of the resulting suspension was used as the DNA template for PCR verification to confirm the transformation. Two PCR reactions were performed: one confirmed the absence of the gene using primers located within the gene of interest (for *PDE2* and *RME1)* or one primer located within the gene of interest and another primer outside it (for *HCM1)*, while the second PCR verified the integration of the NAT cassette using primers designed to anneal outside the gene of interest and within the NAT cassette. Information regarding plasmids and primers used is provided in **Supplementary Table 2**.

### Disk diffusion assays

*Candida* cultures were diluted to 1 × 10^6^ cells/mL in sterile water and 100 μL of each diluted culture was evenly spread on a YPD plate using glass beads. The plates were allowed to dry, and a disk containing 25 μg FLC (9166, Liofilchem) was placed in the center of each plate. For disk diffusion assays performed at different pH, the pH was adjusted using NaOH/HCl to reach pH 4, 6, 8, and 10 from that of the regular YPD (pH ∼6.5). For disk diffusion assays with different agents, the agar was supplemented with the following concentrations: FK506 0.5 µg/ml, geldanamycin 0.5 µg/ml, fluphenazine (FNZ) 10 µg/ml, oligomycin 10 µg/mL, omeprazole 32 and 128 µg/ml. For disk diffusion assays with acetic acid, 10 µl of glacial acetic acid (>99%) was added to blank disks (9999, Liofilchem). For disk diffusion assays under anaerobic conditions, the plates were grown using the AnaeroPouch System (Pouch-Anaero, Mitsubishi Gas Company) and AnaeroPack anaerobic gas generators^35^. The plates were incubated for 48 h at 30°C and photographed using a digital camera or a PhenoBooth+ (Singer Instruments) at 24 and 48 h. Images were analyzed using the R package diskImageR^36^. The radius of inhibition measures the distance (mm) from the center of the FLC disk to the point where a 20% reduction in growth occurs (RAD_20_). The fraction of growth (FoG_20_) calculates the area under the curve from the center of the disk to the 20% cutoff of inhibition. At least three independent disk assays were performed for each experiment.

### Ectopic expression assays

Ectopic expression of transcription factors was achieved using the doxycycline-regulated promoter in pNIM1 or pNIM6^37,38^, as previously described^39,40^. The CAY616 strain containing the pNIM1 vector and *Ca*GFP (*C. albicans*-adapted GFP) was used as the wild type control^40^. Disk diffusion assays were performed by growing *C. albicans* cells overnight in YPD, diluting the cultures to an OD_600_ of ∼0.1 in 3 mL of YPD supplemented with 50 µg/mL doxycycline (+dox). Control cultures were similarly grown in YPD without doxycycline (-dox). After 4 h of growth, the cells were plated on disk diffusion assays (as described above) on the appropriate agar plates (YPD or YPD +dox at 50 µg/mL). Disk diffusion assays were quantified using diskImageR, and RAD_20_ and FoG_20_ values were compared to those of untreated controls (-dox). At least three independent experiments were performed for each strain. Averaged ratios were defined for each strain and compared to the ratio (+dox/-dox) of the parent strain (CAY616). The averaged ratios and standard deviations (SD) of RAD_20_ (susceptibility) and FoG_20_ (tolerance) are provided relative to the WT strain and included in **Supplementary Table 3**.

### Minimum Inhibitory Concentration (MIC) testing

MIC testing was performed using broth microdilution assays in 96-well plates. FLC (PHR1160-1G, Sigma Aldrich) or myriocin (M1177-5M, Sigma Aldrich) were serially diluted (1 in 2) in liquid YPD to final concentrations of 0 to 128 μg/mL. Each well contained 125 μL of liquid YPD volume with *C. albicans* cells at a concentration of 2 × 10^5^ cells/mL. Plates were incubated at 30°C with shaking (200 rpm), and cell densities (OD_600_) were measured at 0, 24, and 48 h using an Epoch2 (Biotek) Spectrophotometer. The plates were examined for contamination after each incubation period. MIC_50_ values were determined after 24 h of growth by identifying the drug concentration leading to ≤ 50% growth relative to growth without FLC. SupraMIC values, or SMG, were determined after 48 h of growth by measuring the average growth levels above the MIC_50_ levels relative to growth without FLC. The same protocol was used for myriocin MICs, with myriocin concentrations ranging from 0 to 128 μg/mL. At least three independent replicates were performed for each experiment.

### Measurements of tolerance by CFU analysis

*C. albicans* overnight cultures were diluted to 5 × 10^3^ cells/mL in PBS, and ∼ 500 cells were spread onto YPD plates with or without 10 µg/mL FLC. The plates were incubated for 48 h at which point the number of colony-forming units (CFUs) was determined for each condition. Tolerance was determined as the fraction of the number of colonies grown on FLC plates relative to the number of colonies growing on plates without drug.

### Checkerboard assays

96-well plates were set up with drug combinations containing FLC (0–128 µg/mL, X axis) and 8-Br-cAMP (0–100 mM, Y axis) or H89 (0–100 µM, Y axis) and serially diluted (1 in 2) in liquid YPD. Each well was seeded with *C. albicans* cells at a concentration of 2 × 10^5^ cells/mL in 125 µL of liquid YPD. Plates were incubated at 30°C with shaking (200 rpm), and cell densities (OD_600_) were measured at 0, 24, and 48 h using a Biotek Epoch 2 plate reader. Plates were examined for contamination after each incubation period. FLC MIC_50_ values were determined after 24 h of growth for each row by identifying the drug concentration leading to less than or equal to 50% growth relative to the growth without FLC. Three to four independent assays were performed for each strain.

### ScanLag assay

The ScanLag assay was adapted from a previous study^11^ with minor changes. The strains were streaked from glycerol stocks onto YPD agar and incubated for 24 h at 30 °C. For experiments using PKA chemical modulators, the YPD plates were supplemented with 50 μM H89 or 100 mM 8-bromo-cAMP. Colonies were suspended in 1 mL PBS, diluted to 10^4^ cell/ml, and ∼500 cells were spread onto YPD plates with or without 10 µg/mL FLC. The plates were placed on scanners at 30°C and scanned every 30 min for 96 h. For experiments with light-sensitive compounds (e.g., 8-bromo-cAMP), scanning was initiated with a 12 h delay to reduce the exposure of reagents to light. Image analysis was done in MATLAB using the “ScanLag”^41,42^ adapted for yeast cells by changing the identification of the size of the colony to a minimum of 20 pixels. Experiments were performed with biological triplicates.

### Fluorescent imaging of lipid droplets

A total of 10^5^ cells/mL from an overnight *C. albicans* culture was resuspended in YPD in different conditions: no drug, FLC at 128 µg/ml, myriocin at 128 µg/ml. These suspensions were incubated for 3 h at 30°C with shaking (200 rpm). The cell suspension was washed twice with sterile PBS and incubated with 1 µg/mL BODIPY 493/503 (D3922, Invitrogen) for 10 min at 30°C and 200 rpm. The cells were washed twice with sterile PBS and imaged at a 60X magnification with a GFP filter and an exposure of 125 ms on a Zeiss AxioVision Rel. 4.8 microscope. The Speckle Counting pipeline of Cellprofiler4.2.6^43^ was used to identify smaller objects within larger ones and to establish a relationship between them. Objects with diameters of 30 to 100 pixels were considered yeast cells, while objects with a diameter of 1 to 6 pixels were classified as lipid bodies. The same pipeline was used to quantify several parameters: the number of lipid bodies within each cell, the highest pixel intensity within the lipid bodies, referred to as brightness, and the integrated intensity of lipid bodies, quantified by summing the pixel intensities within these structures. All assays were performed with three biological replicates, with at least 200 cells analyzed for each replicate.

### Fluorescent imaging of vacuoles

To stain the vacuoles, *C. albicans* cells from an overnight culture were diluted to 10^5^ cells/ml, resuspended in YPD or YPD+FLC (128 µg/ml), and incubated with 8 µM of FM4-64/SynaptoRed C2 (S6689, Sigma) for 3 h at 30°C with shaking. The cell suspension was washed with 1 mL of fresh YPD and transferred to 4 mL of YPD before incubating for 90 min at 30°C and 200 rpm. The cells were washed with sterile PBS before imaging at a 60X magnification with a mCherry filter and an exposure of 100 ms on a Zeiss AxioVision Rel. 4.8 microscope. The Speckle Counting pipeline of Cellprofiler4.2.6 was used to identify objects with diameters of 10 to 35 pixels, which were considered vacuoles. The sum of the pixel intensities within these vacuoles, represented by the integrated intensity, and the area of the vacuoles were quantified using the same pipeline. All assays were performed with three biological replicates, with at least 200 cells analyzed for each replicate.

### Fluorescent imaging of Mnl1

Overnight *C. albicans MNL1*-Neon cultures were diluted 1 in 50 in fresh YPD or YPD containing 10 µg/mL FLC, followed by incubation at 30°C with shaking for 4 h. Cells were then washed twice with sterile PBS and resuspended in PBS with 2.5 µg/mL DAPI (Thermo Scientific 62247). The cells were imaged on a Zeiss AxioVision Rel. 4.8 microscope at a 60X magnification with the DAPI filter with an exposure of 150 ms and with a GFP filter for 300 ms.

### Fluorescent imaging of mitochondria

A total of 10^5^ cells/mL from an overnight *C. albicans* culture was resuspended in YPD or YPD with FLC (128 µg/ml) and incubated for 3 h at 30°C and 200 rpm. The cells were washed twice with sterile PBS and incubated with 500 nM MitoTracker GreenFM (M7514, Invitrogen) and 500 nM MitoTracker Red CMXRos (M7512, Invitrogen) for 30 min at 30°C and 200 rpm. After this incubation, the cells were washed twice with sterile PBS and imaged using a Zeiss AxioVision Rel. 4.8 microscope at a 60X magnification with a GFP filter for 150 ms and an mCherry filter for 100 ms. All assays were performed with three biological replicates, with at least 200 cells analyzed for each replicate.

### Flow cytometry with MitoTracker dyes

The cells from an overnight culture were diluted 1/50 in 2 mL of YPD or YPD with FLC at a concentration of 128 µg/mL and incubated for 3 h at 30°C with shaking (200 rpm). The cell suspension was then washed twice with sterile PBS and resuspended in PBS with 500 nM of MitoTracker GreenFM and 500 nM MitoTracker Red CMXRos (M7512, Invitrogen) for 30 min at 30°C and 200 rpm. Next, the cells were washed twice with sterile PBS and diluted to a density of ∼10^6^ cells/mL in PBS. Fluorescence intensity was measured in 500 µl aliquots of cell suspension and data were collected from 10^5^ cells per sample using a Miltenyi MACSQuant Analyzer 10 Flow Cytometer. Cell populations were gated by SSC (SSC <10^3^ A.U) and FSC (FSC >10^4^ A.U.) to eliminate debris. Experiments were performed with three biological replicates. Analyses were performed using FlowJo 10.9.0.

### Flow cytometry with rhodamine 6G

Cells from an overnight *C. albicans* culture were diluted 1/50 in 2 mL of YPD with 1 µg/mL rhodamine 6G (10045820, Thermo Scientific Chemicals). The cell suspensions were incubated at 30°C with shaking (200 rpm) for 2 h and 24 h. Following these incubation periods, the cells were washed twice with sterile PBS and diluted to a density of ∼10^6^ cells/mL in PBS. Fluorescence was measured in 200 µl aliquots of cell suspension, and data were collected from 10^5^ cells per sample using a Miltenyi MACSQuant Analyzer 10 Flow Cytometer. Cell populations were gated by SSC/FSC to eliminate small debris particles. Experiments were performed with three biological replicates. Analyses were performed using FlowJo 10.9.0 software.

### Flow cytometry with azole-Cy5

Cells from an overnight *C. albicans* culture were diluted 1/50 in 2 mL of YPD with 1 µM azole-Cy5, followed by incubation at 30°C with shaking for 2 h and 24 h. For experiments using PKA chemical modulators, the cultures were supplemented with 50 μM H89 or 100 mM 8-bromo-cAMP. Following incubation, the cells were washed twice with sterile PBS and diluted to a density of ∼10^6^ cells/mL in PBS. To assess viability, the cells were stained with 0.5 mg/mL propidium iodide (PI). Fluorescence measurements were conducted on 200 µl aliquots of the cell suspension, with data collected from 10^5^ cells per sample using a Miltenyi MACSQuant Analyzer 10 Flow Cytometer. The cell populations were gated by SSC/FSC to exclude small debris particles. The experiments were conducted with three biological replicates, and analyses were performed using FlowJo 10.9.0 software.

### Sphingolipid quantification

*C. albicans* cells were cultured overnight in 3 mL of liquid SD medium (2% glucose, 0.67% Bacto yeast nitrogen base without amino acids, pH 5.4) at 30°C with shaking (200 rpm). Cells were harvested by centrifugation and washed twice with sterile PBS. Sphingolipid quantification was performed as previously described^28,44^. Lipid signals were normalized to the cellular phosphate (Pi) content. Lipid species that were not detected in any sample were excluded from further analysis. Only species present at levels exceeding or equal to 0.05 pmol of lipid per nmol of Pi were considered for analysis. All lipid quantifications were performed in biological triplicates, with mean values and standard errors of the mean (SEM) reported in **Supplementary Table 8**.

### Galleria mellonella infections

*G. mellonella* larvae were obtained from Vanderhorst Wholesale (Saint Marys, OH, USA), and the infection protocol was adapted from a previous study^11^. Larvae were kept at 15°C and used within 1 week of delivery. Larvae without signs of melanization and an average weight of 0.25 g were selected for each experiment in groups of 12. *C. albicans* inocula were prepared from cultures grown overnight at 30°C in liquid YPD and washed twice with sterile PBS. Cell densities (OD_600_) were measured using a SPECTRAmax 250 Microplate Spectrophotometer and adjusted to 3 × 10^7^ cells/mL in sterile PBS. Each larva was first injected with 3 × 10^5^ *C. albicans* cells via the last left pro-leg using a 10 µL glass syringe and a 26S gauge needle (Hamilton, 80300). A second injection with either FLC (1 mg/kg), FNZ (10 mg/kg), FLC+FNZ (1 mg/kg and 10 mg/kg), or sterile vehicle (PBS) was done via the last right pro-leg 2 h post-infection. The inoculum size was confirmed by plating of fungal cells on YPD. Infected and control groups of larvae were incubated at 37°C for 10 days, and survival was monitored daily. Larvae were recorded as dead if no movement was observed upon contact. Experiments were performed with 2-4 independent replicates (*n* = 24-48 total larvae per condition). Control groups of larvae included treatment with individual drugs (FLC or FNZ) without *C. albicans* infection. No significant killing of larvae was observed in either of these conditions relative to injections with PBS alone. Statistical differences between larval groups were tested using Wilcoxon matched pairs signed rank tests, * *P* <0.01.

### RNA Sequencing and analysis

*C*. *albicans* cells were cultured overnight in liquid YPD medium at 30°C with shaking (200 rpm) and diluted 1 in 50 in 5 mL YPD, YPD with 0.5 μg/mL FLC, or YPD with 0.25% glacial acetic acid. These concentrations were selected based on the MIC_50_ values of the WT strain for growth in FLC or acetic acid, using half of these values (**Supplementary Fig. 6a**). After 14 h at 30°C with shaking, 1 mL of the cell suspension was used to extract total RNA using the Ribopure-Yeast RNA kit (AM1926, Invitrogen), according to the manufacturer’s instructions. A Bioanalyzer (Agilent) was used for the qualitative sample analysis, and samples with RNA quality (RIN) scores of 7 or higher and a 260/280 ratio of 2.1 to 2.2 were used for sequencing. The Illumina Stranded Total RNA Prep, Ligation with Ribo-Zero Plus kit (20040525, Illumina) was used to convert total RNA into up to 384 dual-indexed libraries. The libraries were sequenced with the Biomics Platform at Institut Pasteur using pair-end Illumina stranded-mRNA sequencing. RNA sequences reported in this paper have been deposited in the NCBI Sequence Read Archive under BioProject ID PRJNA1165997.

The paired-end reads from the RNA-seq data were trimmed for low-quality reads, and Illumina TruSeq adapters were removed with Cutadapt v1.9.1^45^ with the following parameters: --trim-qualities 30 -e (maximum error rate) 0.1 --times 3 --overlap 6 --minimum-length 30. The cleaning of rRNA sequences was performed with Bowtie2 v2.3.3^46^ with default parameters. The cleaned reads from RNA-seq paired-end libraries were aligned to the *C*. *albicans* SC5314 reference genome (version_A22-s07-m01) with Tophat2 v2.0.14^47^ and the following parameters: minimum intron length 30, minimum intron coverage 30, minimum intron segment 30, maximum intron length 4000, maximum multihits 1, and microexon search. Genes were counted using featureCounts v2.0.0^48^ from the Subreads package (parameters: -t gene -g gene_id -s 2 -p). Analysis of differential expression data was performed in DESeq2^49^. The statistical analysis process includes data normalization, graphical exploration of raw and normalized data, tests for differential expression for each feature between conditions, raw P-value adjustment, and export of lists of features with significant differential expression between conditions. The genes with adjusted *P*-values less than 0.05 were considered differentially expressed compared to the non-treated condition (WT YPD). Differentially expressed genes between WT and *mnl1* cells are reported in **Supplementary Table 6**. The data variability was explored by performing hierarchical clustering and principal component analysis (PCA) of the whole sample set after the counts were transformed using a variance stabilizing transformation. Hierarchical clustering was calculated using Euclidean distance and the Ward criterion for agglomeration. PCA was performed using DESeq2. Log_2_-fold changes for *mnl1* relative to the parent strain or FLC relative to YPD were plotted as heatmaps using GraphPad Prism, and asterisks were used to denote significant *P-* adjust values as described above.

Gene annotation was performed by using the gene ontology (GO) database specific to C. *albicans*. We used the org.Calbicans.SC5314.eg.db annotation resource package for fungal species (https://github.com/lakhanp1/fungal_resources/tree/main/C_albicans/org.Calbicans.SC5314.eg.db) developed by the Wong Lab at the University of Macau. This package includes gene-based information from the CGD database and is open-source under the GNU GPL v2 license. We used the Bioconductor clusterProfiler package^50^ to generate a gene-to-Gene Ontology (GO) assignment table for all genes. Gene Set Enrichment analysis (GSEA) was performed using the “gseGO” function and GO Enrichment Analysis was performed using the “enrichGO” function. Our analysis categorized differentially expressed genes into three domains: Biological Process, Cellular Component, and Molecular Function. Only the Biological Process domain was reported in this study. This systematic approach facilitated a detailed exploration of the functional landscape associated with the differentially expressed genes, shedding light on their roles and interactions within the *C. albicans* genome.

### qRT-PCR

cDNA was made from the RNA isolated for sequencing using the Verso cDNA Synthesis Kit (Thermo Scientific, AB1453A). qRT-PCR reactions were performed using the SsoAdvanced Universal SYBR Green Supermix (1725271, Bio-Rad) in a CFX Opus 96 Real-Time PCR System (Bio-Rad). The PCR cycling conditions consisted of an initial denaturation at 95°C for 1 min, followed by 35 cycles of denaturation at 95°C for 10 s, and annealing/extension at 60°C for 30 s. qRT-PCRs were performed in triplicate, each with two technical replicates. The 2^−**CT**^ method was used for quantification, and the data were normalized to the reference gene *MAC1.* The validation of several genes differentially expressed in the RNA sequencing dataset was performed using qRT-PCR and is included in **Supplementary Fig. 15**. Gene expression levels are shown as log_2_-fold changes relative to WT cells grown in YPD.

### cAMP quantification

The cAMP-Glo Assay (V1501, Promega) was used to measure cAMP concentrations according to the manufacturer’s instructions. A cAMP standard curve was generated for each experiment. *C*. *albicans* cells were cultured to exponential phase in liquid YPD medium at 30°C with shaking (200 rpm). 1 mL of overnight culture was incubated with 25 μL of lyticase (1000 U/ml) for 1h at 30°C with shaking. The cells were washed twice with sterile PBS and diluted to 2 × 10^6^ cells/ml. A 10 μL aliquot containing 10^4^ cells was used for the cAMP-Glo assay, and the protocol for 96-well plates was followed. The experiments were conducted with three biological replicates, and analyses were performed using FlowJo 10.9.0 software.

### Whole genome sequencing, variant detection, and coverage analysis

WT and *mnl1* cells were cultured in 5 mL YPD medium overnight at 30°C with continuous shaking at 200 rpm. The DNA was extracted using the Qiagen Genomic DNA kit, utilizing the QIAGEN Genomic-tip20/G. Everything was performed according to the manufacturer’s instructions. Sequencing libraries were prepared by SeqCenter (Pittsburgh, PA) using the Illumina DNA Prep and sequenced on an Illumina NovaSeq 6000. The genome sequences and General Feature Files for the SC5314 reference genome (haplotype A, version A22-s07-m01-r130) were downloaded from the *Candida* Genome Database (http://www.candidagenome.org/). Reads were aligned to the SC5314 reference genome (haplotype A chromosomes) using Minimap2 version 2.17^51^. Sequence Alignment/Map tools (SAMtools) v1.10 (r783) and Picard tools version 2.23.3 (http://broadinstitute.github.io/picard) were used to filter, sort, and convert the SAM files. Single nucleotide polymorphisms (SNPs) were called using the Genome Analysis Toolkit (GATK) version 3.6^52,53^, according to the GATK Best Practices. SNPs were filtered using the following parameters: VariantFiltration, QualByDepth (QD) <2.0, LowQD, ReadPosRankSum <−8.0, LowRankSum, FisherStrand (FS) >60.0, HightFS, MQRankSum <12.5, MQRankSum, MQ <40.0, LowMQ, HaplotypeScore >13.0. Coverage levels were calculated using GATK. Besides passing GATK’s filters, we also classified heterozygous positions as those with an allelic ratio of the number of alternative allele reads/total number of reads comprised between 20% and 80% and homozygous positions as those with an allelic ratio of the number of alternative allele reads/total number of reads of >80% and <20%. The number of heterozygous positions was calculated per 1 kbp window using vcftools^54^ and the genotypes assigned with GATK. To estimate nuclear and mitochondrial chromosome copy numbers, the read alignment depth was calculated for 1 kbp windows across each genome and normalized to the average depth per genome. Sequence read data have been deposited in the NCBI Sequence Read Archive under BioProject ID PRJNA1165997.

### Network Analyses with PathoYeastract+

To identify regulatory interactions among the transcription factors (TFs) associated with azole susceptibility or azole tolerance, we used the PathoYeastract+ database (http://pathoyeastract.org/). The “Rank Genes by TF” module was employed to extract known and predicted regulatory relationships between the TFs based on experimental evidence from gene expression and DNA-binding studies. The resulting network was visualized and analyzed to identify highly interconnected TFs and key regulatory hubs. Enrichment of interactions within the tolerance-associated TF set compared to the rest of the *C. albicans* genome was assessed using a hypergeometric test (*P* <0.05). The network was categorized into functional modules (e.g., stress response, unstressed log phase growth) based on Gene Ontology (GO) annotations and published literature.

### Statistical analyses

All experiments represent the average of three or more biological replicates; error bars represent the standard error of the mean. Statistical analyses were performed using two-tailed *t* tests, one-way analysis of variance, and Wilcoxon matched-pairs signed rank tests using Microsoft Excel or Prism 10 (GraphPad). *R*^2^ tests were used for simple linear regressions, and two-tailed nonparametric Spearman correlation tests were used to test correlations. All error bars show means +/-SEM. Significance was assigned for *P* <0.05, asterisks denote *P* values as follows: **** *P* <0.0001; *** *P* <0.001; ** *P* <0.01; * *P* <0.05.

## RESULTS

### Identification of transcription factors that regulate *C. albicans* azole responses

To identify transcription factors (TFs) that affect FLC susceptibility (RAD_20_) and/or tolerance (FoG_20_), we screened two isogenic TF libraries (196 knock out (KO) mutants and 225 overexpression (OE) mutants that have 178 genes in common) using FLC disk diffusion assays analyzed with established methods^28^. We focused on the genes that consistently altered FLC drug responses (susceptibility and/or tolerance) by ±20% relative to the WT when both deleted and overexpressed. Nine of these TFs affected susceptibility (RAD_20_, a measure of resistance): Ace2, Hfl1, Hms1, Mrr1, Pho4, Stp2, Tac1, Tec1, and Upc2 (**Supplementary Table 3**). The identification of Mrr1, Tac1 and Upc2, which are important for the expression of MDR transporters, ABC efflux pumps, and ergosterol biosynthesis genes^55–57^, respectively, provided confidence in the screen. We focused on tolerance-specific TFs by excluding those that altered susceptibility, which yielded 31 TFs that altered tolerance (FoG_20_) when either deleted or overexpressed (**Supplementary Table 3**); among these, Crz1 and Iro1 were previously implicated in FLC tolerance^11,14^. The screen revealed known and novel TFs that modulate FLC responses, with more TFs affecting tolerance (31) than susceptibility (9), highlighting the prominent role of TFs in modulating tolerance.

### FLC tolerance correlates with intracellular FLC accumulation and time of colony appearance

We previously examined genetically diverse *C. albicans* isolates and found that tolerance levels inversely correlated with the intracellular accumulation of a fluorescent azole probe^11^. However, the contributions of diverse genetic factors could not be ruled out because these strains differed from one another at thousands of genetic loci. To test the association of azole accumulation and tolerance in isogenic strains, we compared 48 TF KO mutants with diverse tolerance levels for steady-state intracellular azole-Cy5 accumulation 24 h after drug exposure. As seen earlier^11^, tolerance inversely correlated with azole-Cy5 accumulation (r = -0.78, *P* <0.001, two-tailed nonparametric Spearman correlation, **Fig. 1a**). By contrast, susceptibility did not correlate well with azole-Cy5 accumulation at 24 h (r = 0.42, *P* = 0.002, two-tailed nonparametric Spearman correlation, **Supplementary Fig. 1a**). This strengthens the idea that reduced drug accumulation contributes to the growth of tolerant subpopulations at supraMIC azole concentrations. Previously, we also found that highly tolerant strains form colonies with a shorter relative time of appearance (ΔToA, defined as the delay in first detection of colonies on supraMIC FLC concentrations relative to on drug-free plates)^11^. ToA is measured using ScanLag assays, which track colony emergence over time using flatbed scanner imaging^11,42^. In bacteria, ToA is a proxy for lag phase duration^42^. Since *C. albicans* cells transiently arrest growth upon FLC exposure^58^, we propose that ToA serves as a proxy for exit from FLC-induced cell cycle arrest. For the 11 TF KO mutants examined, tolerance level was inversely correlated with ΔToA (r = -0.846, *P* <0.001, two-tailed nonparametric Spearman correlation, **Fig. 1b**). Thus, higher tolerance is associated with faster colony appearance on drug, suggesting faster exit from cell cycle arrest. The relationship between tolerance, intracellular azole accumulation, and shorter lag time in isogenic TF mutants reinforces results first seen with genetically diverse isolates^11^, and supports the idea that lower intracellular drug levels drive tolerance. Notably, resistant isolates have faster ΔToA than tolerant strains, and the ΔToA level does not correlate with susceptibility levels (r = 0.196, *P* = 0.543, two-tailed nonparametric Spearman correlation, **Supplementary Fig. 1b**). This highlights a distinction between the mechanisms underlying tolerance and resistance. Resistant strains do not arrest appreciably following exposure to supraMIC drug concentrations. We posit that this is because resistance mutations largely mitigate the detrimental physiological effects of the drug.

**Figure 1:**
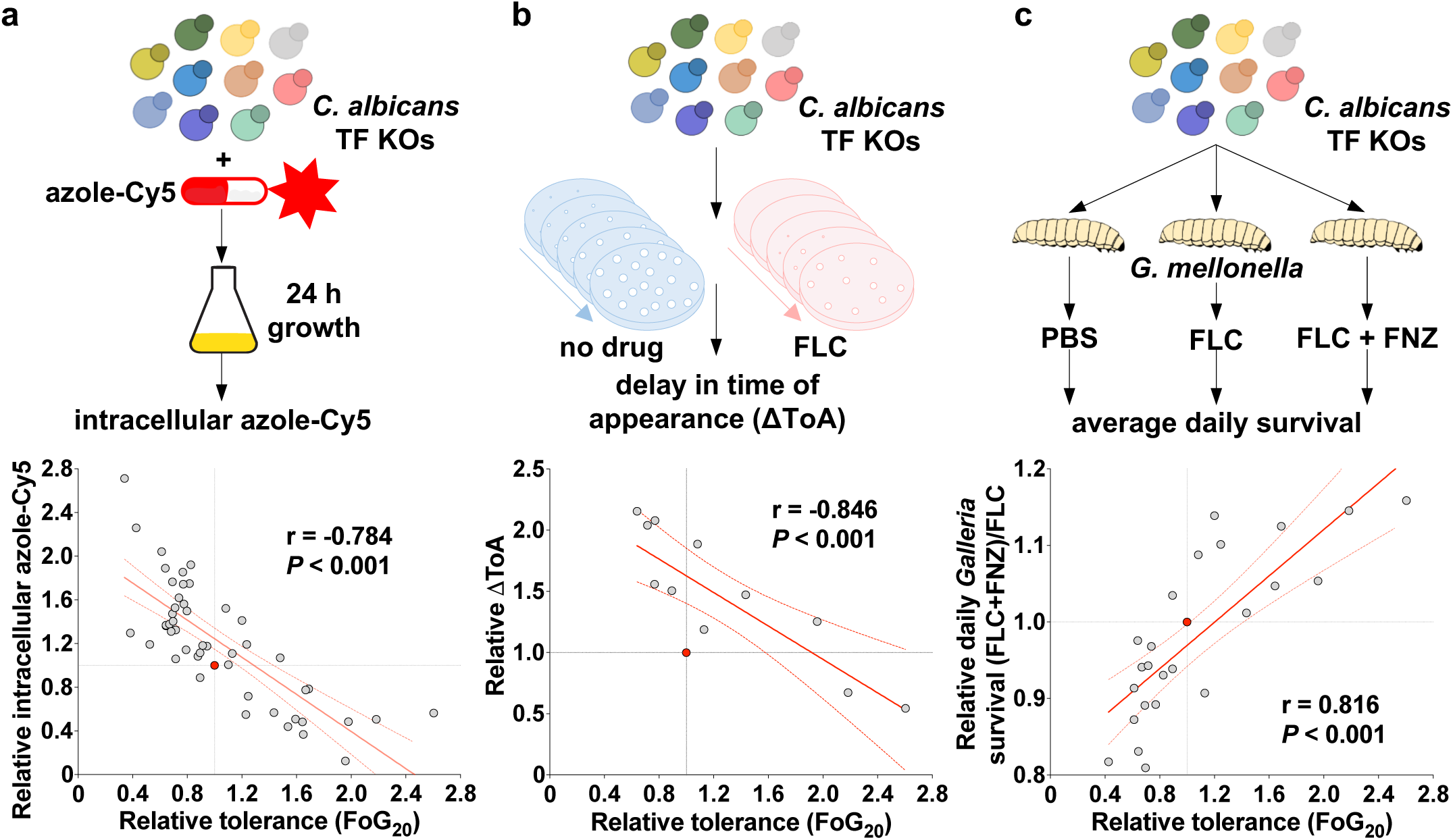
Azole tolerance correlates with intracellular azole-Cy5 accumulation, the time of colony appearance in the presence of drug, and the clearance of infection by an adjuvant *in vivo*. **a**, 48 *C. albicans* TF KOs strains were given Cy5 conjugated-fluconazole (azole-Cy5, 1 µg/mL) for 24 h, then steady state intracellular levels of azole-Cy5 were measured by flow cytometry. The plot shows the correlation between tolerance (FoG_20_) and intracellular azole-Cy5, both parameters shown relative to the corresponding WT strain. **b,** The time of colony appearance (ToA) on plates containing FLC (10 µg/mL) or no drug was measured using ScanLag assays for 11 TF KO mutants. The plot shows the correlation between FLC tolerance (FoG_20_) and ΔToA (the difference in the ToA on FLC relative to the ToA on plates without drug), relative to the corresponding WT strain. **c,** 24 *C. albicans* TF KO strains were used to infect *G. mellonella* larvae, which were subsequently given either PBS (control), FLC (1 mg/kg), or a combination of FLC (1 mg/kg) and fluphenazine (FNZ, 10 mg/kg). Survival of larvae was monitored daily for 14 days, and the percent survival was calculated for each treatment group daily (n = 12 larvae/group; 2-4 groups/condition). The plot shows the correlation between relative FLC tolerance (FoG_20_) and relative daily survival of *G. mellonella* treated with FLC+FNZ relative to FLC alone ((average daily survival with FLC+FNZ)/(average daily survival with FLC)), with both parameters shown relative to the corresponding WT strain. Correlations were assessed using two-tailed nonparametric Spearman correlation coefficients, and significance was noted for *P* <0.05. In all plots, the red dot indicates data for the parental strain.

### Fluphenazine enhances the efficacy of FLC therapy *in vivo,* especially with high-tolerance isolates

As previously observed in diverse isolates, adjuvant drugs like fluphenazine (FNZ) can enhance FLC efficacy both *in vitro* and *in vivo*^11^. *In vitro*, the FLC+FNZ combination eliminated tolerance in AHY135, the parental strain of the TF KO collection (**Supplementary Fig. 1c**), consistent with the prior study. Furthermore, FNZ increased the intracellular accumulation of azole-Cy5, with a higher proportion of cells showing high fluorescence (**Supplementary Fig. 1d-e**). Thus, FNZ boosts FLC activity by disrupting tolerance and increasing intracellular drug levels.

To ask if FNZ disrupts tolerance *in vivo*, we used the *Galleria mellonella* larval infection model, which recapitulates key features of systemic *C. albicans* infection, including growth at 37°C and engagement with host immunity. Each larva was inoculated with one of 24 different TF KO mutants and the parental strain; two hours later, larvae were treated with PBS (control), FLC (1 µg/mL), or FLC+FNZ (1 µg/mL and 10 µg/mL). The mutants exhibited distinct virulence profiles and varied responses to FLC treatment (**Supplementary Fig. 1f**): average daily survival ranged from 7.7% (*rob1*) to 35.5% (*rim101*), compared to 25.8% for the WT strain. FLC treatment increased WT-infected survival by 1.9-fold (to 49%), with a broad range of effects in the mutants, from >4-fold increases (e.g., *dal81*, *rob1*) to no benefit (e.g., *crz1*, *hap43*). Notably, neither susceptibility nor tolerance alone predicted the effect of FLC (*P* >0.5 for both RAD_20_ and FoG_20_, two-tailed nonparametric Spearman correlation). By contrast, FLC+FNZ had a modest impact on the strains (0.9- to 1.29-fold relative to FLC alone) and positively correlated with tolerance, but not with susceptibility (**Fig. 1c**, **Supplementary Fig. 1f**; FoG_20_, r = 0.816, *P* <0.001; RAD_20_, *P* >0.5; two-tailed nonparametric Spearman correlation). Thus, the FLC+FNZ combination is most effective against highly tolerant strains.

### A highly interconnected “tolerance network” of TFs controls FLC responses

Of the 31 TFs that altered tolerance by ±20% in both TF KO and TF OE datasets without affecting susceptibility (**Fig. 2a**, detailed analyses in **Supplementary Fig. 2**), four TFs decreased tolerance when deleted and when overexpressed (Grf10, orf19.3928, Ron1, and Stb5), and 13 TFs increased tolerance when deleted and when overexpressed (Arg81, Bcr1, Chp2, Efg1, Fcr3, Fgr15, Irf1, Mnl1, orf19.173, Rap1, Rim101, Zcf39, and Zfu3). In addition, 11 TFs (Csr1, Crz1, Gat1, Iro1, Isw2, Mig2, orf19.1150, Rob1, Ume7, Yng1, and Zfu2) were designated “tolerance enhancers” because they increased tolerance when the gene was overexpressed and decreased tolerance when the gene was deleted; 3 other TFs with the opposite effects (Ahr1, Rlm1, and Zcf2) were termed “tolerance repressors” (**Supplementary Fig. 3a**). Gene Ontology (GO) processes significantly enriched among these 31 TFs (>10-fold, *P* <0.05, FDR <0.01) included regulation of adhesion, response to metal ions, pH response, nutrient response, regulation of filamentous growth, growth control, asexual reproduction, biofilm formation, and inorganic ion homeostasis (**Supplementary Table 4**). These findings suggest that TFs involved in azole tolerance, but not susceptibility, are primarily linked to ion sensing and growth regulation pathways.

**Figure 2:**
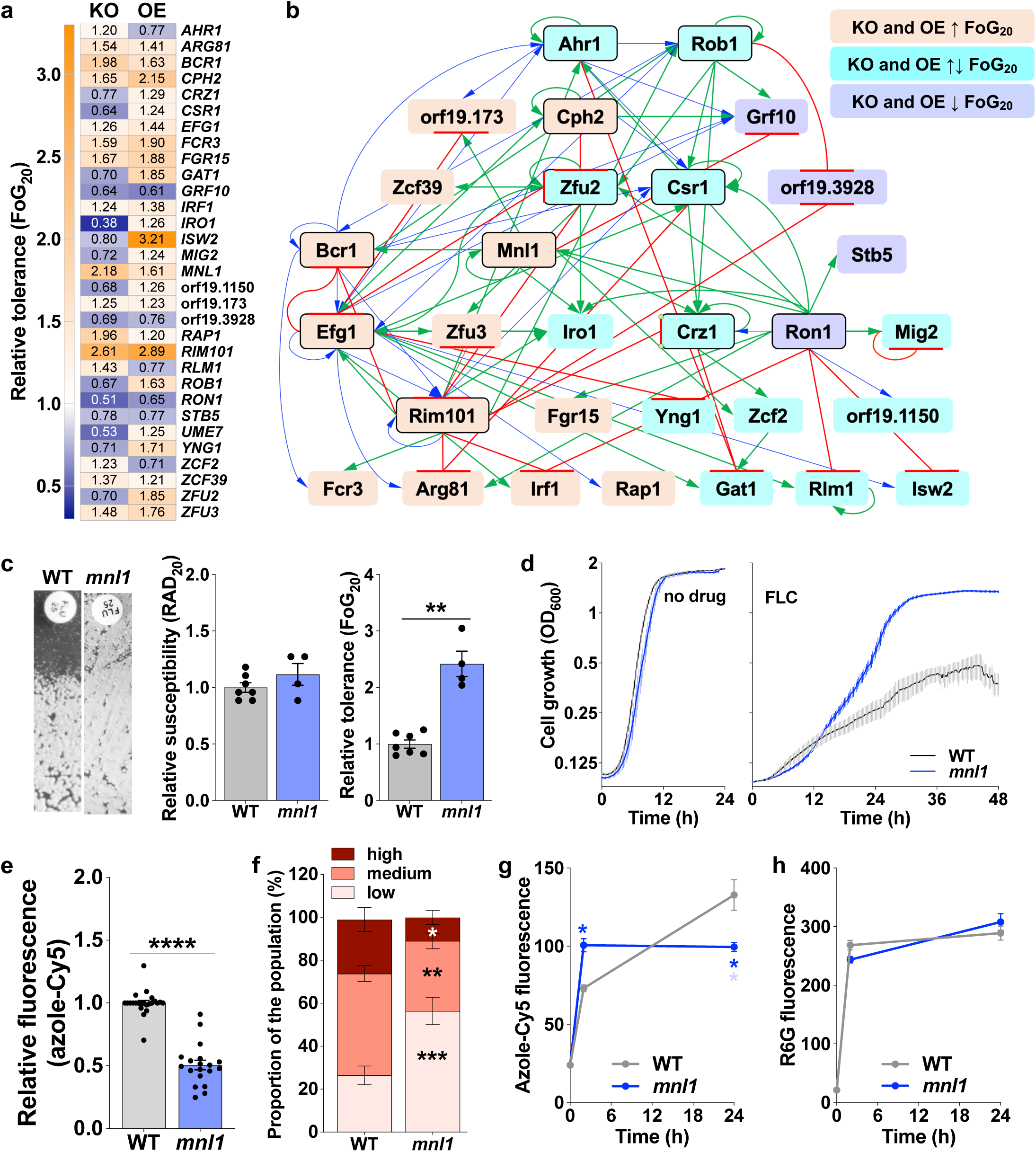
A highly interconnected network of 30 TFs modulates FLC tolerance. **a,** Changes in tolerance (FoG_20_) levels upon deletion (KO) or overexpression (OE) of the 31 regulators identified in the two TF screens. Tolerance is shown relative to the corresponding WT controls. **b,** Regulatory relationships among the 31 tolerance TFs based on gene expression or DNA binding evidence (PathoYeastract+), grouped according to the impact on FLC tolerance of overexpressing or deleting the respective TF: orange, both KO and OE mutants had increased tolerance; purple, both KO and OE mutants had decreased tolerance; turquoise, KO and OE mutants had opposite effects on tolerance. TFs Ahr1, Cph2, Crz1, Csr1, Efg1, Mnl1, Rim101, Rob1, Ron1, and Zfu2 (highlighted in black boxes) show significant enrichment of interactions for the targets in this set relative to the rest of the *C. albicans* genome *(P* <0.005, hypergeometric test). Interactions are marked by arrows according to the direction of the interaction: green, positive regulation; red, negative regulation; blue, unspecified. **c,** FLC disk diffusion assays of WT and *mnl1* cells, and RAD_20_ and FoG_20_ quantifications. **d,** Growth curves of WT and *mnl1* in the presence of FLC 10 µg/mL FLC. **e,** Intracellular azole-Cy5 fluorescence intensity levels for WT and *mnl1*, measured at 24 h relative to the WT strain. **f,** Population distribution of intracellular azole-Cy5 levels for WT and *mnl1*, based on azole-Cy5 accumulation levels. High, medium, and low levels were defined by flow cytometry analysis based on Cy5 fluorescence intensity, with gates set at <200 (low), between 200 and 2000 (medium), and >2000 (high). **g-h,** Time course of intracellular azole-Cy5 (g) and rhodamine 6G (h) fluorescence intensity levels for WT and *mnl1* measured at 2 and 24 h. Asterisks show significant differences relative to the corresponding WT condition (nonparametric t tests, * *P* <0.05, ** *P* <0.01, *** *P* <0.001, **** *P* <0.0001).

Regulatory interactions among the 31 tolerance-associated TFs were identified using PathoYeastract+^59^ revealing a highly interconnected “tolerance network” (**Fig. 2b**). These TFs were significantly enriched as targets of 11 regulators (Ahr1, Bcr1, Cph2, Crz1, Csr1, Efg1, Mnl1, Rim101, Rob1, Ron1, and Zfu2), relative to the genome-wide background (>10-fold enrichment, *P* <0.05, hypergeometric test, **Supplementary Table 5**). By contrast, a separate, smaller “resistance network” comprised 9 TFs (Ace2, Hfl1, Hms1, Mrr1, Pho4, Stp2, Tac1, Tec1, Upc2) that were associated with FLC susceptibility (**Supplementary Fig. 4**, **Supplementary Table 5**). The smaller size and lower complexity (average of 2.6 interactions between each of the TFs) of the susceptibility network highlight the broader, more interconnected (average of 3.1 interactions per TF) transcriptional program underlying azole tolerance, in contrast to the more compact regulatory architecture of *bona fide* resistance. Thus, a distinct regulatory architecture enables *C. albicans* to overcome FLC-induced stress.

Within the TF tolerance network, two modules were distinguishable - a stress response module and an unstressed log phase growth module (**Supplementary Fig. 3b-c**). The stress module included “alkaline”, “pH”, “weak acid”, and “drug stress”. Consistent with known functions, Rim101 was central to the alkaline and pH stress modules, and Mnl1 was central to the weak acid stress response^27,60,61^. Ahr1, Efg1, and Ron1 were central regulators in the unstressed growth module. Ahr1 regulates amino acid metabolism, cell cycle, phenotypic switching, and adhesion^62–64^. Ron1 regulates N-acetylglucosamine catabolism^65^, and Efg1 controls carbon metabolism, phenotypic switching, and morphogenesis^66–68^. The distinct stress response and growth modules emphasize the importance of both adaptive and baseline regulatory processes. Consistent with this, single-cell transcriptomic studies under prolonged FLC exposure found that *C. albicans* cells bifurcate into two distinct groups: ribo-dominant and stress-dominant subpopulations^69^. The ribo-dominant subpopulation prioritizes growth through ribosomal and mitochondrial gene expression, and the stress-dominant state promotes survival via stress-related genes (including TFs Cph2, Efg1, Mnl1, orf19.173, Rim101, Zcf2, and Zcf39). The two subpopulations reflect a strategic balance between growth and stress responses, underscoring the complex interplay of processes that contribute to FLC tolerance.

### Mnl1 represses FLC tolerance by regulating intracellular drug accumulation

Among the TFs identified in our screen, Mnl1 emerged as a key regulator: tolerance increased when *MNL1* was either deleted or overexpressed. Mnl1 ranked high in connectivity within the tolerance network (**Fig. 2a-b**, **Supplementary Table 5**). Deletion of *MNL1* increased tolerance without affecting susceptibility (**Fig. 2c**) and, under high FLC concentrations (10 µg/mL), *mnl1* cells (obtained by deleting both *MNL1* alleles) enhanced growth in liquid media, particularly after 24 h of FLC exposure (**Fig. 2d**). Among the TF deletion mutants, *mnl1* had the second-highest tolerance, following *rim101*. Given that the role of pH in tolerance and its association with Rim101 have been described previously^11,70^, we prioritized Mnl1 for further studies. Consistent with its high tolerance level, *mnl1* cells accumulated less azole-Cy5 at steady state, with a larger subpopulation having low intracellular drug levels (**Fig. 2e-f**). This phenotype was consistent for the TF KO library and for independently constructed *mnl1* deletion strains^27^, and was largely rescued by reintegration of a single *MNL1* allele (**Supplementary Fig. 5a-e**). These results suggest that the TF Mnl1 limits tolerance by promoting intracellular drug accumulation. To explore the dynamics of drug accumulation, we measured azole-Cy5 accumulation in *mnl1* cells at early (2 h) and late (24 h) times after FLC exposure (**Fig. 2g**). In WT cells, intracellular drug levels increased over time, whereas *mnl1* cells maintained significantly lower levels at 24 h. To determine whether differences in drug efflux could explain this phenotype, we measured rhodamine 6G fluorescence efflux, a proxy for ABC transporter activity (e.g., Cdr1, Cdr2)^71,72^. As in the diverse isolates^11^, rhodamine 6G fluorescence was similar in WT and *mnl1* cells (**Fig. 2h**), indicating that reduced azole accumulation in *mnl1* cells is not primarily due to increased efflux through ABC transporters.

### *mnl1* cells have altered sphingolipid homeostasis

To explore the mechanism(s) of high tolerance in *mnl1* cells, we compared the transcriptomes of WT and *mnl1* cells grown in YPD alone, or YPD supplemented with subMIC concentrations of FLC (0.5 µg/mL) or acetic acid (AcAc, 0.25%, **Supplementary Fig. 6a**). Principal component analysis (PCA) revealed that growth conditions drove ∼76% of the variance in gene expression, with acetic acid having the most pronounced effect compared to FLC or YPD alone. An additional ∼15% of the variance was attributable to the *mnl1* genotype (**Supplementary Fig. 6b**). In acetic acid, 3840 genes (62% of *C. albicans* ORFs) were differentially expressed in *mnl1* compared to WT cells; by contrast, only 119 (1.9%) and 84 (1.4%) genes were differentially expressed in YPD and in FLC conditions, respectively (**Supplementary Fig. 6c**, **Supplementary Table 6**). Thus, weak acid stress has a broader effect on the global transcriptional landscape, and Mnl1 is a key regulator of the weak acid stress response. In FLC, ergosterol biosynthetic genes and their transcriptional regulator, Upc2^57^, were induced to comparable levels in WT and *mnl1* strains. Similarly, expression levels of ABC and MDR transporter genes (including *CDR1*, *CDR2*, and *MDR1*) and their upstream regulators (Mrr1 and Tac1^55,56^) remained comparable between WT and *mnl1* cells with and without FLC (**Supplementary Fig. 6d-f**). Thus, consistent with the rhodamine 6G efflux experiments, increased tolerance observed in *mnl1* mutants is not driven by differential transcription of genes central to azole resistance, reinforcing the role of Mnl1 in tolerance rather than susceptibility.

GO enrichment analysis of genes differentially expressed in the *mnl1* and WT transcriptomes, with or without FLC, revealed lipid metabolism as the most enriched process (>6-fold enrichment, *P* <0.05, FDR <0.01, **Supplementary Table 7**). Indeed, several differentially expressed genes were linked to lipid homeostasis, with others involved in cell wall integrity, stress responses, cytoskeleton, mitochondria, and ion homeostasis (**Supplementary Fig. 7**). Targeted PCA showed distinct clustering of WT and *mnl1* cells, irrespective of FLC exposure, indicating that sphingolipid expression is intrinsically different between WT and *mnl1* cells (**Fig. 3a**). Multiple lipid biosynthesis enzymes were downregulated in *mnl1* cells under both YPD and FLC conditions (**Supplementary Fig. 8a-b**). Lag1 and Sur2 convert sphinganine to dihydroceramide to phytoceramide and phytosphingosine, while Aur1 converts phytoceramide to inositol phosphorylceramide. The sphingomyelin phosphodiesterase Asm3 and the mannosylinositol phosphorylceramide synthase Mit1 were also downregulated, while the flippase Rta2 was upregulated, suggesting compensatory changes.

**Figure 3.**
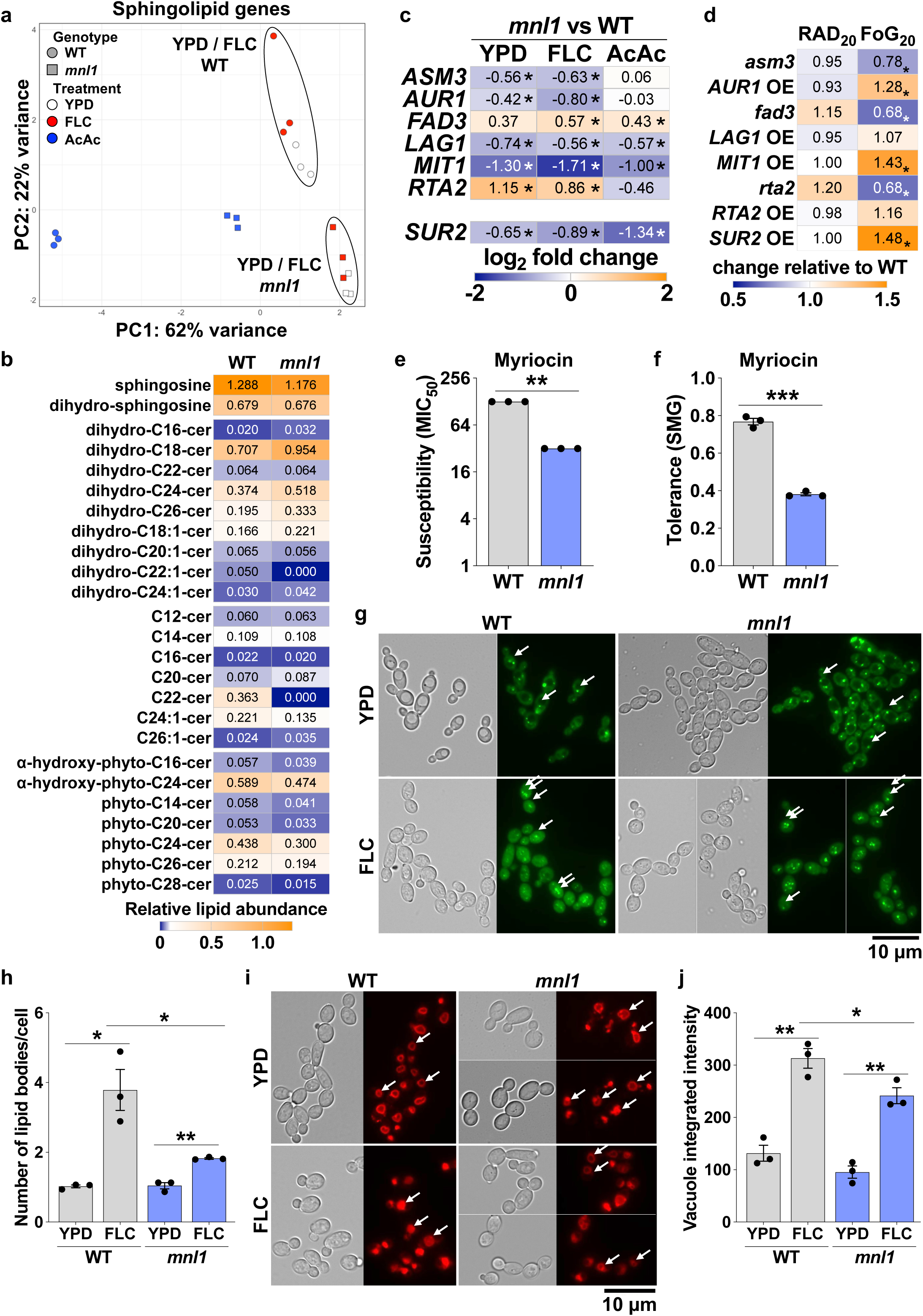
Mnl1 impacts lipid homeostasis. **a,** PCA analysis of sphingolipid genes for WT and *mnl1* cells grown in YPD, FLC, or acetic acid (AA). **b,** Abundance of sphingolipid species in WT and *mnl1* cells, shown relative to inorganic phosphate present in each strain. **c,** Heatmap of log_2_-fold changes in gene expression observed between WT and *mnl1* cells grown in YPD or FLC for sphingolipid genes. **d,** FLC susceptibility (RAD_20_) and tolerance (FoG_20_) of deletion or overexpression mutants of genes involved in sphingolipid metabolism, shown relative to the corresponding WT cells. All mutants were constructed in the SC5314 background. **e-f,** Myriocin susceptibility (MIC_20_, e) and tolerance (SMG, f) for WT and *mnl1 cells*. **g,** BODIPY staining of WT and *mnl1* cells grown in YPD or FLC (128 µg/mL). Scale bar, 10 µm. **h,** Number of lipid droplets per cell for WT and *mnl1* cells grown in either YPD or FLC and stained with BODIPY. **i,** FM4-64 staining of WT and *mnl1* cells grown in YPD or FLC (128 µg/mL). Scale bars, 10 µm. **j,** Integrated intensity of vacuoles stained with FM4-64 in WT and *mnl1* cells grown in either YPD or FLC (128 µg/ml). Asterisks show significant differences relative to the corresponding WT condition or between YPD and FLC conditions (nonparametric t tests, * *P* <0.05, ** *P* <0.01, *** *P* <0.001).

These transcriptional changes were reflected at the lipid level, as *mnl1* cells had altered sphingolipid composition compared to WT cells. Specifically, several long-chain dihydro-ceramide species (including dihydro-C18, C22, C24-cer) were elevated in *mnl1*, while complex ceramides such as C22-cer and phyto-C24-cer were reduced (**Fig. 3b**, **Supplementary Table 8**). These shifts in sphingolipid composition suggest that Mnl1 regulates both the synthesis and maturation of sphingolipids, leading to an accumulation of long-chain dihydroceramides and a depletion of complex ceramides. Given that longer sphingolipid chains increase membrane fluidity, enhance lipid raft formation, and promote permeability^73,74^, these compositional changes likely alter the biophysical properties of *mnl1* membranes and contribute to their altered azole responses.

To investigate whether sphingolipid homeostasis contributes to the high tolerance of *mnl1 cells*, we measured the tolerance of strains overexpressing or lacking genes that are differentially regulated in *mnl1* mutants, including strains overexpressing *AUR1*, *LAG1*, *MIT1*, *RTA2*, and *SUR2*, and strains lacking *ASM3*, *RTA2*, or *FAD3*, which encodes an omega-3 fatty acid desaturase^75^ that is upregulated in *mnl1* in FLC (**Fig. 3c**). Overexpression of *AUR1*, *MIT1*, and *SUR2* increased tolerance, and deletion of *ASM3*, *FAD3*, or *RTA2* reduced it (**Fig. 3d**), without affecting susceptibility. Consistent with these findings, *mnl1 cells* were more susceptible and less tolerant to myriocin, which inhibits an early step in sphingolipid biosynthesis (**Fig. 3e-f**, **Supplementary Fig. 8a**). These results suggest a role for fatty acid composition and sphingolipid architecture in modulating azole tolerance. Moreover, early steps in sphingolipid synthesis may modulate azole susceptibility as well as tolerance.

To further explore lipid-related differences between WT and *mnl1 cells*, we visualized lipid droplets and vacuolar membranes stained with BODIPY or FM4-64, respectively. Lipid droplets are essential for lipid metabolism and stress responses^76^, and their accumulation has been associated with increased membrane permeability and cell death^77^. Without FLC, *mnl1* and WT cells displayed similar numbers of lipid droplets. FLC exposure increased droplet formation in both strains, but to a greater extent in WT cells (3.72-fold vs 1.75-fold in *mnl1*), indicating that WT cells experience higher membrane stress upon azole treatment (**Fig. 3g-h**). Nonetheless, lipid droplet size and fluorescence intensity were similar between the strains (**Supplementary Fig. 8c-d**). Vacuolar morphology also was altered in *mnl1 cells*, which exhibited smaller, less fragmented vacuoles with reduced FM4-64 fluorescence than those of WT cells (**Fig. 3i-j**, **Supplementary Fig. 8e**). Since vacuolar fragmentation is an early response to FLC that precedes growth inhibition^78^, the attenuated phenotype in *mnl1* is consistent with its improved growth under azole stress. These findings support a role for Mnl1 in regulating sphingolipid biosynthesis, increasing lipid droplet formation in response to FLC, and promoting vacuolar fragmentation. We posit that these differences in lipid homeostasis in *mnl1* mutants are central to the cellular responses and stress adaptation to azoles.

### *mnl1 cells* exhibit enhanced FLC-induced mitochondrial activity

Drug efflux is ATP-dependent, and mitochondria support azole resistance by producing ATP, regulating efflux pumps, and mediating iron homeostasis and lipid biosynthesis^79,80^. Thus, we asked whether Mnl1 affects mitochondrial properties. *mnl1 cells* had a 1.26-fold higher mitochondrial genome copy number than WT and showed increased expression of mitochondrial-encoded genes (**Supplementary Fig. 9a**, **Fig. 4a**). By contrast, several nuclear-encoded genes with established or predicted roles in mitochondrial function were downregulated, including *AIF1* (mitochondrial apoptosis), *AUT7* (mitochondrial degradation), *JSN1* (mitochondrial localization), and *PTC6* (mitophagy, **Supplementary Fig. 9b**). These data indicate that Mnl1 loss is associated with increased mitochondrial genome content and transcription and with the repression of nuclear genes required for mitochondrial degradation, suggesting a dysregulation of mitochondrial quality control that could influence drug tolerance.

**Figure 4:**
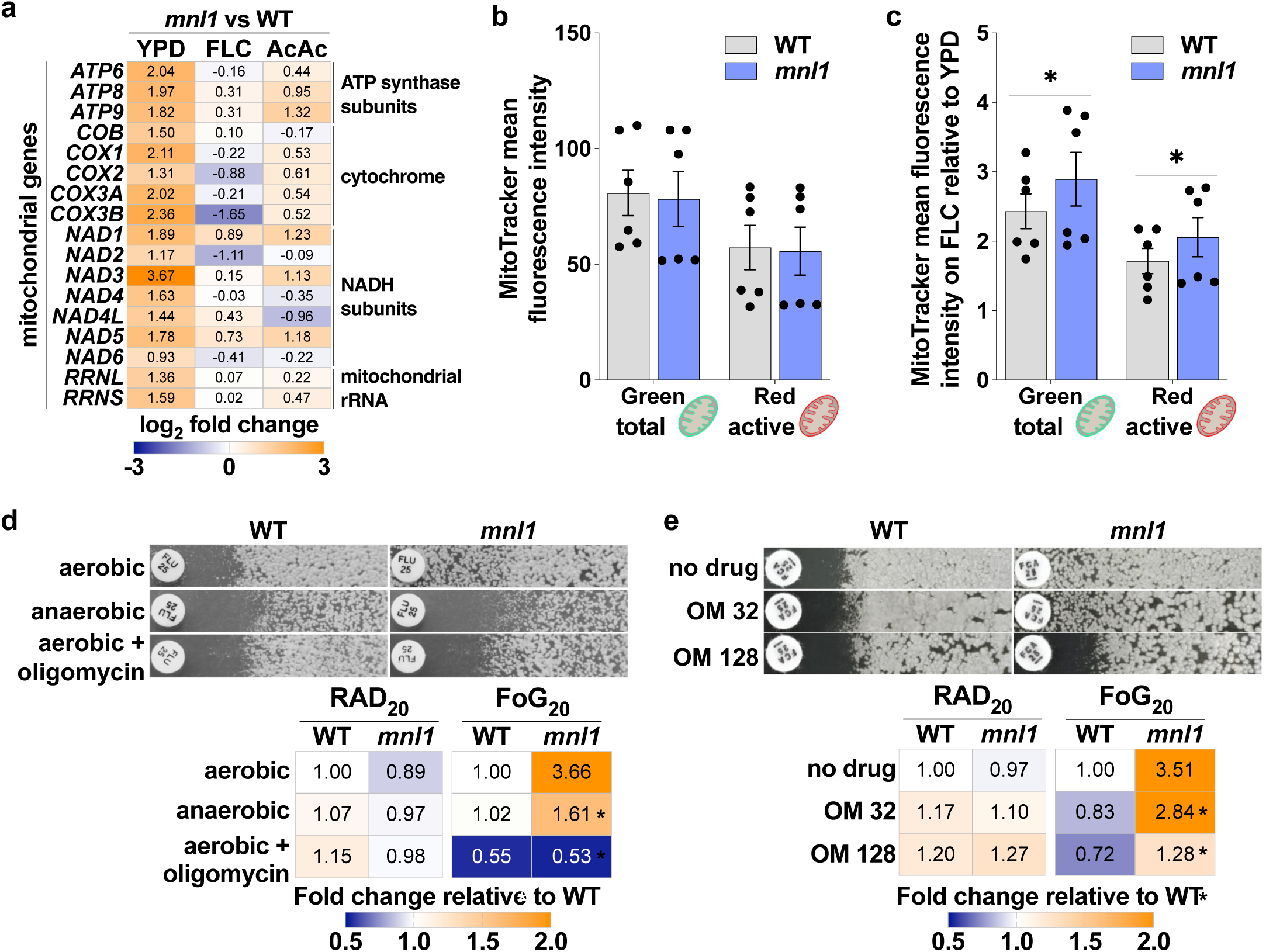
*mnl1 cells* exhibit enhanced mitochondrial activity. **a,** Heatmap of log_2_-fold changes in expression of genes encoded on the mitochondrial genome observed for *mnl1* relative to WT cells grown in YPD or FLC. **b-c,** Effect of FLC on abundance of total (MitoTracker Green) and active (MitoTracker Red) mitochondria for WT and *mnl1 cells.* Mean fluorescence intensity for each MitoTracker dye, in the presence of FLC (128 µg/mL) relative to YPD. **d,** FLC susceptibility (RAD_20_, h) and tolerance (FoG_20_, i) of WT and *mnl1 cells* from FLC disk diffusion assays on YPD plates supplemented with oligomycin (10 µg/mL) or grown under anaerobic conditions. Susceptibility and tolerance values are shown relative to WT cells grown under aerobic conditions. **e,** FLC susceptibility (RAD_20_, h) and tolerance (FoG_20_, i) of WT and *mnl1 cells* from FLC disk diffusion assays on YPD plates supplemented with omeprazole (OM, 32-128 µg/mL). Susceptibility and tolerance values are shown relative to WT cells grown without drug. Asterisks show significant differences relative to the corresponding WT condition or between YPD and drug conditions (nonparametric t tests, * *P* <0.05).

Under basal conditions, the number of total and active mitochondria, assessed by MitoTracker Green (membrane potential–independent) and MitoTracker Red CMXRos (membrane potential– dependent), respectively, was similar between *mnl1* and WT cells (**Fig. 4b**, **Supplementary Fig. 9c**). However, upon exposure to high FLC concentrations (128 µg/mL), *mnl1 cells* exhibited a greater increase in both total and active mitochondria (**Fig. 4c**). These findings suggest that *mnl1 cells* upregulate mitochondrial gene expression, and enhance mitochondrial activity in response to azole stress, features that may collectively support their improved growth under antifungal pressure.

To test whether the high tolerance of *mnl1 cells* depends on mitochondrial function, we measured FLC responses under anaerobic conditions and upon treatment with oligomycin, an inhibitor of mitochondrial ATP synthase. Neither anaerobic growth nor oligomycin treatment significantly altered susceptibility in WT or *mnl1* strains, but both conditions substantially reduced tolerance (**Fig. 4d**). This suggests that mitochondrial respiration is critical for tolerance, but dispensable for susceptibility. To further explore the role of energy metabolism in tolerance, we inhibited Pma1, the plasma membrane proton pump, using omeprazole. Pma1 localizes to lipid rafts and is highly expressed in *mnl1 cells* (1.4-fold increase vs. WT, **Supplementary Fig. 9d**), where it contributes to ATPase activity and membrane potential. Omeprazole reduced *mnl1* tolerance by 2.6-fold at 128 µg/ml, while having minimal impact on WT susceptibility or tolerance (**Fig. 4e**). These results suggest that ATP hydrolysis and proton pump activity support *mnl1*-mediated tolerance, possibly by sustaining membrane energetics under drug stress. Together, these findings indicate that *mnl1* cells rely on enhanced mitochondrial function and active proton pumping to promote azole tolerance.

### Increased cAMP levels and PKA signaling drive FLC tolerance

In the model yeast *S. cerevisiae*, Msn2/4-like TFs like ScCom2, which is an ortholog of Mnl1^81^, interact with the PKA pathway^82^, so we asked about the relationship between Mnl1 and PKA in *C. albicans*. Increased cAMP activates the PKA pathway, which modulates stress responses^20^ and is hypothesized to suppress the Mnl1-dependent weak acid response in dying cells^27^. Under acid stress, Mn1 upregulates Pde2, the phosphodiesterase that converts cAMP to AMP, thereby reducing intracellular cAMP^27^ (**Fig. 5a**). We compared PKA gene expression in the WT and *mnl1* transcriptomes under FLC exposure using targeted PCA. No differential expression of PKA genes or other components of this pathway was detected, suggesting that Mnl1 does not directly regulate the PKA pathway (**Fig. 5b**, **Supplementary Fig. 9b**). We then asked if Mnl1 affected PKA-regulated genes, perhaps indirectly. Indeed, several PKA-regulated genes and ATP-related genes (e.g., ATPases) were differentially expressed in *mnl1*, irrespective of FLC exposure (log_2_-fold change <-0.5 or >0.5, or p-adjust <0.05, **Fig. 5c**, **Supplementary Fig. 9d**). The plasma membrane ATPases Pma1 and Pmc1, and orf19.3219, whose *S. cerevisiae* ortholog is involved in Pma1 activation^83^, were upregulated in *mnl1*. Conversely, Mnn42, a negative regulator of intracellular ATP, was downregulated. Thus, *mnl1 cells* may maintain elevated intracellular ATP levels, potentially enhancing energy-dependent stress responses. Supporting this hypothesis, *mnl1 cells* exhibited 1.44-fold higher basal cAMP levels compared to WT cells (**Fig. 4d**), consistent with increased PKA pathway activity. Overall, these findings suggest that while Mnl1 does not directly control the PKA pathway, its loss elevates basal cAMP and PKA activity, leading to transcriptional changes that favor higher intracellular ATP levels.

**Figure 5:**
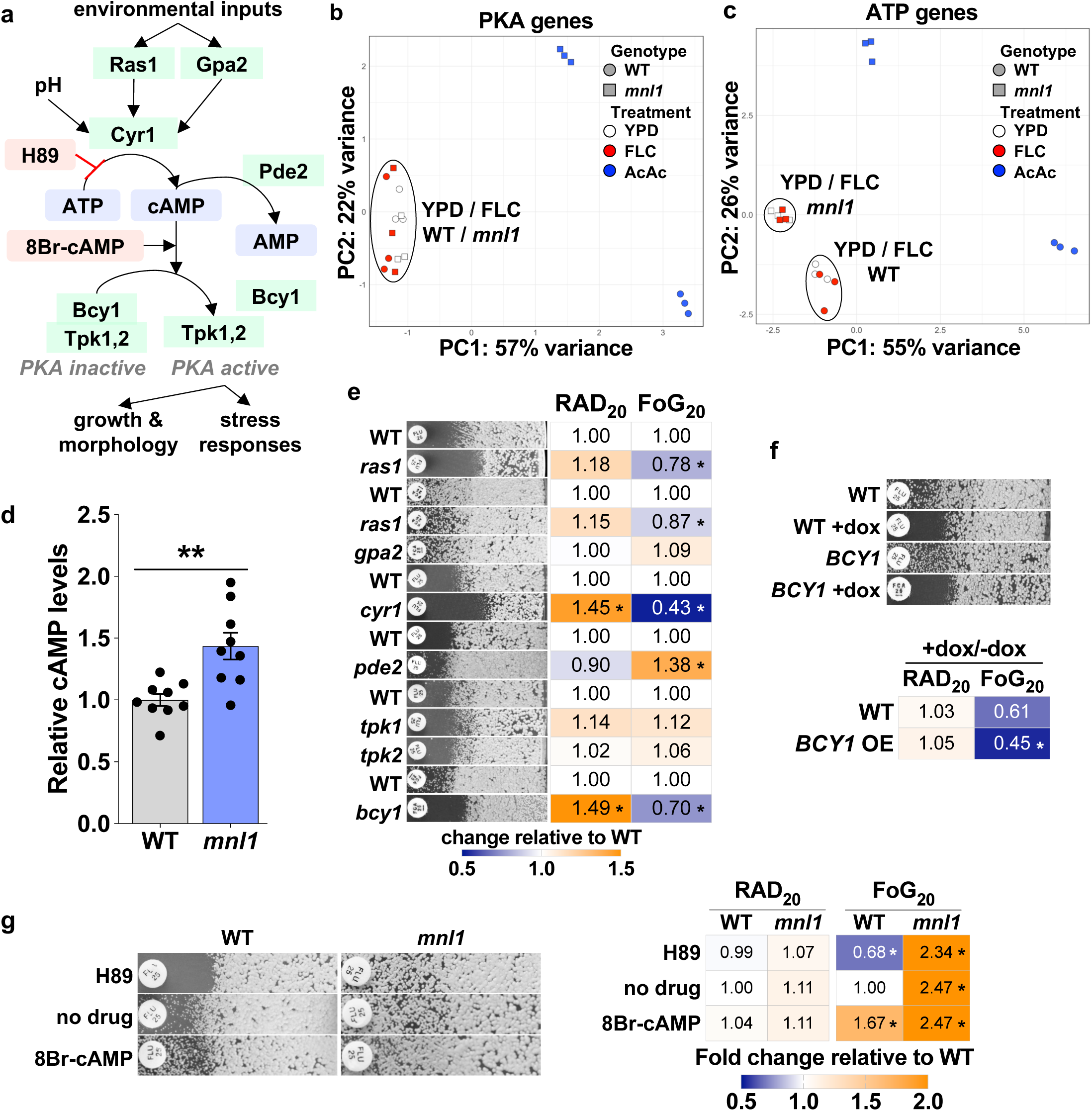
The cAMP-PKA pathway regulates FLC tolerance. **a,** Schematic of the PKA pathway in *C. albicans*, adapted from Hogan et al., 2009^20^. **b-c,** PCA analyses of genes belonging to the PKA (a) and ATP (b) GO terms for WT and *mnl1 cells* grown in YPD, FLC, or acetic acid (AA). Genes included in these GO terms are listed in **Supplementary Table 9. d,** cAMP levels measured in WT and *mnl1 cells*. **e,** Components of the PKA pathway affect FLC responses. Representative images of FLC disk diffusion assays for *ras1* (two independent mutants), *gpa2*, *cyr1*, *pde2*, *tpk1*, *tpk2*, and *bcy1* deletion mutants. All mutants were constructed in the SC5314 background. The corresponding WT strains are shown for each set of mutants. The heatmap shows the susceptibility (RAD_20_) and tolerance (FoG_20_) of the different PKA mutants relative to the corresponding WT strains. **f,** Susceptibility (RAD_20_) and tolerance (FoG_20_) levels for cells overexpressing *BCY1*, with values showing the ratio between cells treated with doxycycline or not (+dox/dox). **g,** Representative images of FLC disk diffusion assays supplemented with the PKA inhibitor H89 (100 µM), or the cAMP analog 8-Bromo-cAMP (8Br-cAMP, 100 mM) for WT and *mnl1 cells*. The DMSO solvent was used as the no-drug control. Asterisks show significant differences relative to the corresponding WT condition or between no drug and H89/8Br-cAMP conditions (nonparametric t tests, * *P* <0.05, ** *P* <0.01).

Given the elevated cAMP levels and mitochondrial activity in *mnl1 cells* during FLC stress, we asked how cAMP-PKA signaling contributes to tolerance by examining components of the PKA pathway (**Fig. 5e**). Deletion of *GPA2*, *TPK1*, or *TPK2*, which encode the G-protein alpha subunit and two catalytic PKA isoforms, respectively, did not affect tolerance. Loss of *RAS1* slightly increased susceptibility and modestly reduced tolerance (∼20% decrease), suggesting a minor role in this context. Strikingly, loss of Cyr1 (adenylate cyclase) or Bcy1 (the PKA regulatory subunit) significantly increased susceptibility (1.45- and 1.49-fold, respectively) and reduced tolerance (0.43- and 0.70-fold, respectively). Moreover, overexpression of *BCY1* modestly decreased tolerance (0.74-fold, **Fig. 5f**), while deletion of *PDE2*, which prevents cAMP degradation, increased tolerance by 1.38-fold (**Fig. 5e**). These data support the idea that elevated cAMP enhances PKA signaling, thereby promoting FLC tolerance. Together, these results emphasize that precise regulation of the PKA pathway, through balanced cAMP levels and stoichiometric control of both catalytic and regulatory subunits, is essential for maintaining the optimal growth–stress response required for azole tolerance.

Next, we modulated intracellular cAMP levels pharmacologically. In WT cells, H89, an inhibitor of ATP-to-cAMP conversion, reduced tolerance by 1.46-fold; conversely, 8-bromo-cAMP, a phosphodiesterase-resistant cAMP analog^84^, enhanced tolerance by 1.67-fold, without affecting susceptibility (**Fig. 5g**). Importantly, in *mnl1 cells*, neither compound significantly altered tolerance, suggesting that tolerance in the absence of Mnl1 is either cAMP-independent or already saturated due to elevated basal cAMP levels. Indeed, multiple assays confirmed that further elevating cAMP in *mnl1* has limited or even detrimental effects (**Supplementary Fig. 10**). Colony formation assays at supraMIC FLC concentrations mirrored these results: H89 reduced and 8-bromo-cAMP increased colony number for the WT, while H89 had no effect and 8-bromo-cAMP slightly reduced colony formation for *mnl1* (**Supplementary Fig. 10a**). These data suggest that excessive cAMP may impair *mnl1* growth under drug stress. H89 also delayed colony emergence in both strains, whereas 8-bromo-cAMP had no impact on *mnl1* (**Supplementary Fig. 10b**). In WT cells, accumulation assays revealed that intracellular azole-Cy5 levels were increased by H89, and decreased by 8-bromo-cAMP, whereas in *mnl1 cells*, both compounds increased azole-Cy5 (**Supplementary Fig. 10c-d**). Furthermore, *mnl1* cell death increased at high 8-bromo-cAMP doses, and excessive cAMP was toxic to both strains in checkerboard assays (**Supplementary Fig. 10e-f**). Indeed, high concentrations of cAMP alone were sufficient to inhibit growth (**Supplementary Fig. 10g-h**). Together, these results imply that cAMP homeostasis critically shapes azole tolerance: moderate increases in cAMP promote PKA signaling and drug tolerance, while excessive cAMP compromises viability, particularly in *mnl1 cells*. Thus, *mnl1* modulates azole tolerance (and not susceptibility) at least in part via alterations in basal levels of cAMP, which consequently affect downstream functions including mitochondrial properties and, we posit, lipid homeostasis.

## DISCUSSION

This study uncovered key transcriptional networks that regulate azole tolerance in *C. albicans*, revealing potential avenues for improving antifungal treatment. By screening TF deletion and overexpression libraries, we identified a highly interconnected network of tolerance regulators that govern growth, stress responses, ion homeostasis, and nutrient signaling. This network encompassed two functional modules: one coordinating stress adaptation and another regulating unstressed growth. Thus, tolerance may arise from the coexistence of two cellular programs: a stress adaptation state and a permissive growth state. The involvement of established regulators such as Ahr1, Crz1, Mnl1, and Rim101 supports the idea that tolerance relies on multiple stress pathways and extensive cellular reprogramming. A consistent feature of TF mutants with higher tolerance levels was that they accumulated less azole-Cy5 and exited FLC-mediated arrest more rapidly. Importantly, in the *G. mellonella* infection model, a tolerance-targeting adjuvant effectively cleared infections by highly tolerant strains. Together, these findings provide a mechanistic framework linking stress homeostasis to azole tolerance, intracellular drug concentration, cell cycle features of drug responses, and highlight promising molecular targets for therapeutic intervention.

Among the TFs identified, Mnl1 emerged as a central regulator of azole tolerance, revealing a broader role in fungal stress responses than previously appreciated. Mnl1 was originally linked to weak acid stress^27^; here, we found that Mnl1 also reduces azole tolerance. A prior study using TF knockout strains had noted altered colony morphology in the *mnl1* mutant on FLC media^85^, which may have reflected tolerance rather than true resistance, although the experimental design did not allow for this distinction. Here, we show that Mnl1 is a key repressor of azole tolerance in *C. albicans*, highlighting mechanisms distinct from classical resistance pathways such as ergosterol biosynthesis and increased drug efflux pump activity. Rather, Mnl1 fine-tunes cellular responses by modulating intracellular azole accumulation, PKA signaling, mitochondrial function, and sphingolipid metabolism, contributing to a distinct stress-responsive state that enables growth under antifungal pressure (**Fig. 6**). Furthermore, *MNL1* was expressed at higher levels in cells with a stress-responsive transcriptional signature (relative to cells with a growth/ribosomal signature)^69^. We posit that these diverse processes are likely to be interdependent, an issue that warrants further investigation.

**Figure 6.**
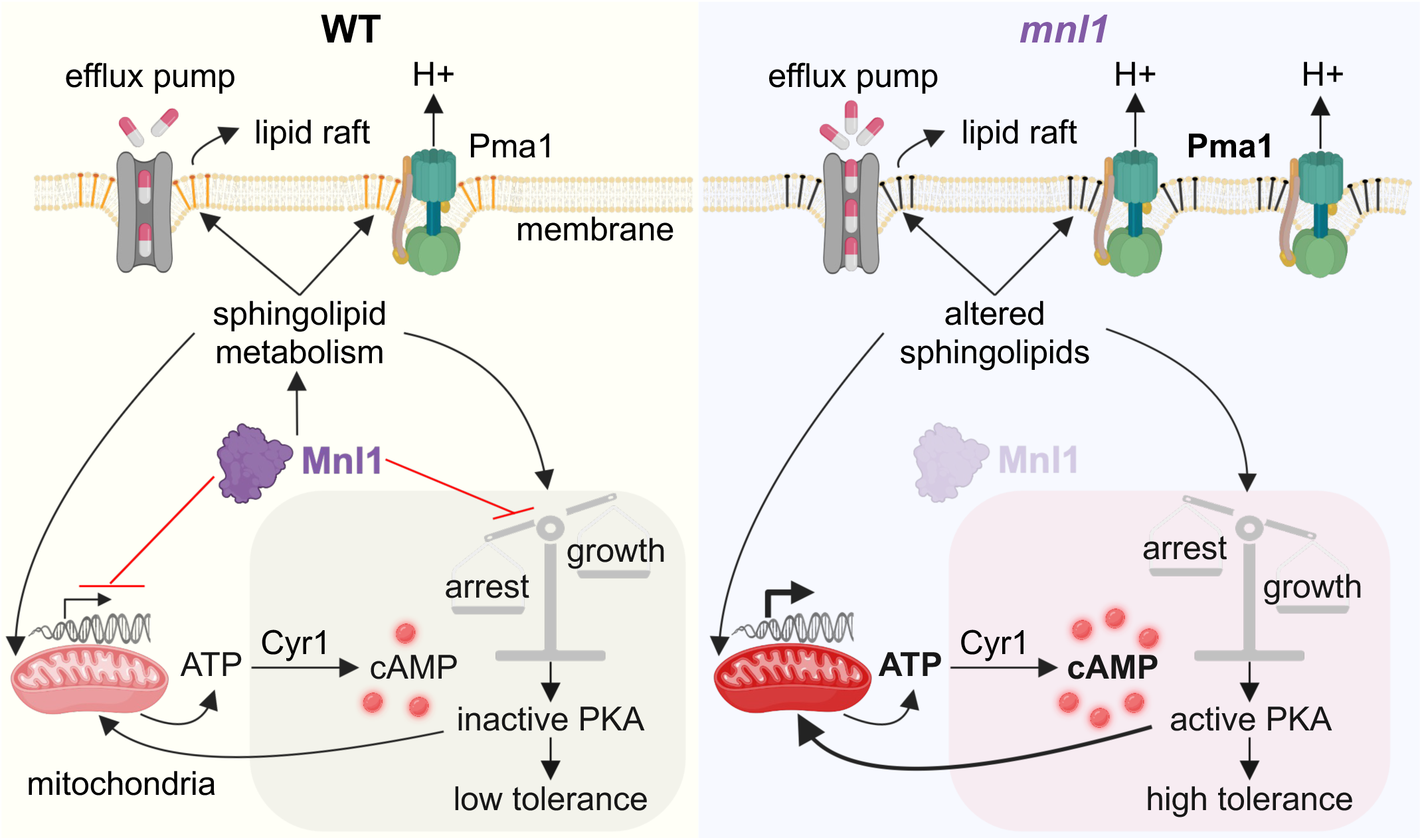
Proposed model for how Mnl1 modulates sphingolipid homeostasis, mitochondrial function, and PKA signaling to coordinately regulate FLC tolerance. WT Mnl1 (left panel) regulates sphingolipid composition, which modulates both the plasma and mitochondrial membranes. In *mnl1* mutant cells (right panel), increased ATP levels elevate cAMP production, driving heightened activation of the PKA pathway, which has a critical role in modulating the balance between cell growth and cell death. *mnl1ΔΔ* cells also display increased mitochondrial gene expression, which can increase ATP production. Elevated ATP levels, along with increased expression of Pma1 (a plasma membrane ATPase located in lipid rafts), enhance energy production, which in turn supports growth under stress. Additionally, changes in sphingolipid composition may alter membrane structure, potentially impacting membrane permeability and the spatial organization of efflux pumps and plasma membrane ATPases like Pma1. This model highlights how metabolic and membrane dynamics collectively influence azole tolerance.

Transcriptomic and lipidomic analyses revealed a distinct sphingolipid profile in *mnl1* mutants, with transcriptional regulation of sphingolipid biosynthetic enzymes matched by differences in specific sphingolipid species. This indicates that Mnl1 regulates membrane composition, particularly lipid raft architecture, domains essential for the positioning and activity of the Pma1 proton pump and drug efflux transporters such as Cdr1^86–88^. The upregulation of a sphingolipid flippase and a fatty acid desaturase in *mnl1 cells* suggests broader changes in lipid architecture, which likely alter membrane permeability and microdomain organization. These changes are functionally linked to tolerance as deletion or overexpression of lipid-modifying genes affected tolerance, but not susceptibility. We posit that these membrane changes compensate, to some degree, for the ergosterol depletion that occurs with FLC exposure.

Interestingly, despite reduced intracellular azole-Cy5 accumulation in *mnl1 cells*, neither transcriptomic data nor rhodamine 6G efflux assays point to increased ABC transporter expression or activity. This suggests that reduced intracellular drug levels are driven by alternative mechanisms, and we suggest that altered membrane lipid composition could partially compensate for canonical drug efflux. While we cannot fully exclude contributions from other transporters or post-transcriptional regulation of efflux activity, the data support a model in which sphingolipid remodeling alters membrane architecture or facilitates transporter compartmentalization, indirectly affecting drug uptake and/or efflux. This mechanism is distinct from classical resistance mechanisms involving direct overexpression of efflux pumps or alterations in Erg11, the drug target.

In parallel, *mnl1 cells* have elevated basal cAMP levels, implicating PKA pathway activation in their enhanced tolerance. This mechanism is likely indirect, as Mnl1 does not transcriptionally regulate PKA pathway genes; however, *mnl1 cells* exhibit metabolic shifts, especially in mitochondrial activity, that likely increase ATP availability, thereby supporting adenylate cyclase activity and boosting cAMP production. Notably, modulation of Bcy1, the regulatory subunit of PKA, reduced tolerance, while removal of Pde2 (a cAMP-degrading phosphodiesterase) increased it, underscoring the importance of maintaining a fine balance of cAMP signaling. While PKA activation promoted growth under FLC stress, excessive cAMP levels in *mnl1 cells* were detrimental, impairing viability and tolerance. Thus, optimal, not maximal, PKA signaling is essential for azole tolerance.

Mitochondrial function was also important for Mnl1-mediated tolerance. *mnl1 cells* upregulated mitochondrial genes and ATPases, consistent with an energy-active state that likely supports drug efflux, lipid remodeling, and stress response programs. This state was not due to increased mitochondrial abundance, but rather enhanced mitochondrial activity. Functional assays confirmed that mitochondrial function is essential for tolerance: both oligomycin treatment and anaerobic growth reduced tolerance without affecting susceptibility, indicating that mitochondrial respiration fuels tolerance mechanisms.

Sphingolipids, the cAMP/PKA pathway, and mitochondria form a tightly interconnected regulatory network controlling metabolism, stress responses, and apoptosis. Literature from model yeast shows that sphingolipids directly regulate mitochondrial function, influencing membrane potential, apoptosis, and mitochondrial outer membrane permeability^89,90^. Another sphingolipid, dihydrosphingosine, induces mitochondrial reactive oxygen species (ROS) and cytotoxicity^91^. The reduced complex ceramides and increased dihydroceramides in *mnl1 cells* suggest that sphingolipid homeostasis could directly modulate mitochondrial function. In *S. cerevisiae*, inhibiting sphingolipid synthesis with myriocin downregulates PKA activity, allowing Msn2/4-mediated stress response activation^92^. Conversely, PKA activation enhances mitochondrial biogenesis and cytochrome levels, thereby promoting mitochondrial function^93^. Mutations that upregulate PKA cause hypersensitivity to sphingolipid biosynthesis defects, while PKA deletion confers resistance^94^. Our study positions Mnl1 at the core of a feedback network in which sphingolipid metabolism, mitochondrial function, and PKA signaling engage in coordinated crosstalk to regulate stress adaptation and establish a growth-permissive, drug-tolerant physiological state (**Fig. 6**).

Historically, azole resistance has been linked to alterations in ergosterol biosynthesis and efflux pump activity. Stress response pathways were often mentioned as contributing to susceptibility/resistance as well^8,29,80,95^. However, these reports reflect a time when little distinction was made between susceptibility and tolerance. The work here emphasizes that the ability to tolerate drug stress involves a physiological state that depends upon several stress response pathways. Unlike resistance mechanisms that rely on drug-induced gene expression changes, we found that FLC does not drive a major transcriptional response in tolerant cells. Instead, azole tolerance in *C. albicans* can arise from a coordinated rewiring of metabolic and signaling pathways. The transcription factor Mnl1 acts as a master regulator of this alternative state, linking membrane composition, intracellular signaling, and bioenergetic capacity. Thus, tolerance and susceptibility are distinct, and tolerance cannot be a mere failure of drug action. Rather, tolerance is an active, energy-dependent adaptation strategy that relies on an alternative physiological state.

## ACKNOWLEDGEMENTS

We thank Prof Al Brown (University of Exeter, Exeter) for the gift of *MNL1*/*MSN4* mutant strains, Dr Benjamin Heineike (Charité Universitätsmedizin, Berlin) for fruitful discussions on the PKA pathway, Dr Kevin Carlson (Brown University, Providence RI) for support with flow cytometry experiments during the TF screen, Dr Pedro Monteiro (Universidade de Lisboa) and Prof Miguel Teixeira (Institute for Bioengineering and Biosciences) for assistance with the PathYeastract+ analyses. This work was funded through the following grants: FRM Espoirs de la Recherche Postdoctoral Fellowship (to EIM), Brown University UTRA (to CE), NIH NIAID R21AI139592 (to IVE), ANR GENOMEHET (to IVE), PTR Carnot Pasteur (to IVE), CIFAR Azrieli Global Scholarship (to IVE), NIH AI166869/AI168222/AI141893 (to RJB), NIH AI125770 (to MDP), European Research Council (ERC) under the European Union’s Horizon 2020 research and innovation program (grant agreement No 951475, to JB), and the Israel Science Foundation (grant No. 2644/24) within the Biomedical Research Grants Track of MAVRI program (to JB). MDP is a recipient of the Research Career Scientist (RCS) Award (IK6 BX005386) from the Department of Veterans Affairs. MDP is a Co-Founder and Chief Scientific Officer (CSO) of MicroRid Technologies Inc. The goal of MicroRid Technologies Inc. is to develop new antifungal agents for therapeutic use.

