## Supplementary Information for "Sphingolipid Homeostasis, Mitochondrial Activity, and PKA Signaling Drive an Azole-Tolerant State"

This document includes:

- **Supplementary Results**
- **Supplementary Figure Legends**
- **Supplementary Table Legends**
- **Supplementary References**
- **Supplementary Figures**

### Supplementary Results

**Tolerance is more sensitive to perturbations in TF gene expression relative to susceptibility.** The two TF libraries were screened using FLC disk diffusion assays and analyzed using established methods[1]. A cutoff of ±20% was used for selecting TFs with altered susceptibility and/or tolerance levels. Normalization of the drug responses to the corresponding wild type (WT) strain revealed that susceptibility was less variable (SD = 0.16X and 0.09X of WT values for TF KO and TF OE collections, respectively) than tolerance (SD = 0.35X and 0.45X of WT values for TF KO and TF OE collections, respectively) (**Supplementary Fig. 2a**, **Supplementary Table 3**). Of the 196 TF KO mutants, 73 and 121 mutants affected $RAD_{20}$ and $FoG_{20}$ responses, respectively. Among the 225 TF OE mutants, 14 and 87 mutants affected $RAD_{20}$ and $FoG_{20}$, respectively (**Supplementary Fig. 2b**). Notably, a larger fraction of mutants affected tolerance rather than susceptibility (61.7% of KO and 38.7% of OE mutants affected tolerance compared to 37.2% of KO and

6.2% of OE mutants which affected susceptibility); this implies that tolerance is more sensitive to changes in TF gene expression relative to susceptibility. Furthermore, very few mutants caused higher resistance (decreased $RAD_{20}$, 2/75 (2.7%) and 1/14 (7.1%) of KO and OE mutants, respectively, **Supplementary Fig. 2b**). Intriguingly, over half of the KO mutants with altered susceptibility also had altered tolerance (40/73 (54.8%), as did 10/14 (71.4%) of OE mutants (**Supplementary Fig. 2c**). This suggests that many of these TFs contribute to both FLC resistance and tolerance. To increase the confidence of the identified hits, we selected TFs that altered susceptibility/tolerance in both TF KO and OE screens. This resulted in 9 TFs affecting susceptibility (**Supplementary Fig. 2d**). To identify tolerance-specific TFs, we excluded those impacting susceptibility, yielding 81 KO and 77 OE mutants. Among these, 31 TFs overlapped between the KO and OE libraries and were selected for further analysis (**Supplementary Fig. 2d**).

**Tolerance does not correlate with changes in growth dynamics.** We previously examined the connection between tolerance, susceptibility, and the ability of cells to grow, using 7-20 clinical isolates that differed in their genetic backgrounds[2,3] and identified general trends among *C. albicans* isolates. Here, we screened mutants constructed in a single genetic background (SC5314) to focus on the effects of specific genes. As in our prior study, there was no correlation between tolerance and susceptibility for either the TF KO or the OE collections (**Supplementary Fig. 2e-f**; two-tailed nonparametric Spearman correlation, $P$ >0.05 for both collections). Consistent with the previous study, neither the doubling time nor the lag phase length in liquid growth measured for mutants from the TF KO collection (**Supplementary Table 11**) correlated with tolerance (**Supplementary Fig. 2g-h**; two-tailed nonparametric Spearman correlation, $P$ >0.05). This supports the idea that tolerance is not a direct consequence of alterations in growth dynamics.

**A compact transcriptional network underlies FLC susceptibility.** A subset of nine transcription factors (TFs) emerged as regulators of FLC susceptibility, based on the observation that both their deletion and overexpression consistently altered $RAD_{20}$ without affecting tolerance (**Supplementary Fig. 4a**). These included Ace2, Hfl1, Hms1, Mrr1, Pho4, Stp2, Tac1, Tec1, and Upc2. These TFs are implicated in various biological processes, including growth, filamentation, metabolism, and drug responses (see GO term analysis in **Supplementary Table 4**). Together, these TFs form a distinct susceptibility network that is markedly less interconnected than the broader tolerance network (**Supplementary Fig. 4b**), suggesting that azole resistance relies on a simpler, more modular regulatory architecture. Within this network, Ace2, Hms1, Mrr1, Pho4, Stp2, Tec1, and Upc2 (highlighted with black outlines) appear as central nodes, exhibiting significantly enriched regulatory interactions with other network members relative to genome-wide expectations (**Supplementary Table 5**). Notably, for most TFs in this module, both loss and overexpression led to increased susceptibility (higher $RAD_{20}$, **Supplementary Fig. 4a**), pointing to the importance of maintaining optimal expression levels for resistance. These findings support the notion that azole resistance depends on a finely balanced transcriptional program, distinct from the stress-adaptive and highly interconnected tolerance network.

**Transcriptome profiling of *mnl1 cells* reveals broad rewiring of stress, lipid, and mitochondrial pathways**. To define the transcriptional program controlled by Mnl1, we performed RNA-seq on *mnl1* and WT cells grown in YPD, YPD+FLC (0.5 μg/ml), or YPD+AcAc (acetic acid, 0.25%). Across all three conditions, the *mnl1 strain* displayed significant changes in gene expression relative to WT, affecting diverse cellular processes (**Supplementary Fig. 7a**, **Supplementary Table 6**). A core set of differentially expressed genes was shared among YPD alone, FLC, and AcAc conditions, including genes predicted or documented to be involved in lipid homeostasis (*MIT1*, *LAG1*, *RSD1*, *SUR2*), ion homeostasis (*IRO1*, *orf19.5754*), stress-response (*CTA26, IML2*, *SIP5*), mitochondria (*AIF1*, *JSN1*), ATP (*orf19.3219*), and PKA signaling (*PGA23*) (**Supplementary Fig. 7b**). A second group of overlapping changes was shared between YPD and FLC conditions, marked by an altered expression of lipid homeostasis (*AUR1*, *ASM3, RTA4*), stress response (*FGR29*, *SOD5*), and metabolic (*URO99*, *PMU2*) genes. A third and larger set of lipid homeostasis (*ATG15,*

*DRS2, YJU3*), transport (*CTA3*, *SVL3*), ion homeostasis (*HAP43*), ATP metabolism (*PMA1*, *PMC1*), cell wall integrity (*DFI1*, *PNG2*, *PHR1*, *PIR1*, *BMT3-5*, *EXG2, SKN1*), cell cycle (*CDC20*, *CCN1*), and stress response (*CEK1*, *CRZ1, TPS1*) genes were commonly regulated in the YPD and AcAc conditions. Finally, several cell cycle (*CDC34*), cytoskeleton organization (*BBC1*, *BOI2*, *BUD14*), cell wall integrity (*CAS5*, *CHS8*, *MNN15*, *SUN41*), ion homeostasis (*ZRT2*), mitochondria (*AUT7*), lipid homeostasis (*FAD3*), and stress response (*ASR1-2*, *STP4*) genes were differentially expressed in the FLC and AcAc conditions.

In addition to these shared responses, each condition elicited distinct *mnl1*-specific expression signatures (**Supplementary Table 6**). These results indicate that Mnl1 activity connects lipid and ion homeostasis, PKA signaling, cell wall integrity, and a broader stress-response network involving major regulators such as Crz1, Cas5, Cek1, Stp4, and Tps1. Even without drug exposure, Mnl1 shapes a transcriptional program that primes cells for stress adaptation. Its loss reprograms this network toward a state that changes FLC tolerance by modulating lipid metabolism, stress signaling, mitochondrial function, and cell wall integrity.

**Calcineurin and Hsp90 inhibitors reduce FLC tolerance in WT and *mnl1* cells.** To assess the role of stress response pathways in mediating the FLC tolerance of *mnl1* cells, we performed disk diffusion assays in the presence or absence of geldanamycin (Hsp90 inhibitor), FK506 (calcineurin inhibitor), or fluphenazine (FNZ). Geldanamycin and FK506 modestly increased FLC susceptibility in both strains (**Supplementary Fig. 11**). Importantly, similar to the tolerance of diverse strains[2], the higher tolerance of *mnl1* was sensitive to the addition of adjuvant drugs, including FNZ, FK506, and geldanamycin (**Supplementary Fig. 11**). These results demonstrate that stress response regulators such as Hsp90 and calcineurin contribute to FLC tolerance even in *mnl1*, and that their pharmacological inhibition can overcome their elevated tolerance phenotype.

**Mnl1 regulates the TF orf19.173, which affects intracellular drug accumulation**. *Orf19.173* encodes a predicted transcriptional target of Mnl1 under acetic acid stress conditions[4]. It is positioned downstream of Mnl1 in the stress module and downstream of Ahr1 in the growth module of the tolerance network (**Supplementary Fig. 3b-c**). Deletion of *orf19.173* increased FLC tolerance without affecting susceptibility (**Supplementary Fig. 12a**), suggesting a role in modulating azole responses. Under high FLC conditions (10 µg/mL), the *orf19.173* strain exhibited a modest growth enhancement in liquid culture compared to the WT (**Supplementary Fig. 12b**). Similar to *mnl1*, *orf19.173* cells had reduced azole-Cy5 accumulation and a larger subpopulation with low intracellular fluorescence levels, although the effect was less pronounced (**Supplementary Fig. 12c-d**). These phenotypes support the idea that *orf19.173* functions downstream of Mnl1 and may partially account for the effect of Mnl1 on tolerance. Tracking azole-Cy5 uptake dynamics revealed a modest increase in azole-Cy5 levels in *orf19.173* cells from 2 to 24 h (**Supplementary Fig. 12e**). Furthermore, rhodamine 6G efflux was unchanged relative to the WT strain (**Supplementary Fig. 12f**), indicating that the reduced azole accumulation in *orf19.173* is not due to increased efflux via ABC transporters. Collectively, these data suggest that Mnl1 represses tolerance in part through *orf19.173*, while also acting via additional, unidentified targets.

**Mnl1, and its *S. cerevisiae* ortholog Com2, regulate FLC tolerance.** In the model yeast *S. cerevisiae*, Msn2/4-like TFs co-regulate general stress responses. ScMsn2 and ScMsn4 are paralogs that activate transcription by binding to STRE elements in promoter regions[5,6], a response down-regulated by the PKA pathway[7]. ScCom2 (Yer130c), a third member of the ScMsn2/4 TF family, is phosphorylated by the PKA pathway and regulates sulfur dioxide stress responses at low pH[8,9]. However, *C. albicans* stress responses have diverged significantly from *S. cerevisiae*[10,11]. *C. albicans* has only two Msn2/4-like family members: Msn4 is closest to ScMsn2/4, and Mnl1 is an ScCom2 ortholog[4]. Unlike in *S. cerevisiae*, the *C. albicans* Msn2/4-like TFs Msn4 and Mnl1 do not coordinate a general stress response; by contrast, Mnl1 regulates the weak acid stress response[4,10,12], while Msn4 has unknown functions. In the tolerance network, Mnl1 is central to the stress response module, positively regulating TFs Bcr1, Crz1, Iro1, orf19.173, Zcf2 and negatively regulating Grf10 (**Supplementary Fig. 3b**). In FLC disk diffusion assays, neither Mnl1 nor Msn4 affects FLC susceptibility, while the deletion of *MNL1* increased FLC tolerance and was complemented by the reintegration of an intact copy of *MNL1* (**Supplementary Fig. 5a-c**). Furthermore, deleting both *MNL1* and *MSN4* increased tolerance more than deleting *MNL1* alone (**Supplementary Fig. 5a,c**). Thus, Mnl1 inhibits tolerance and does not affect susceptibility. Interestingly, we found a similar role for ScCom2: deletion of *ScCOM2* increased FLC tolerance without affecting susceptibility (**Supplementary Fig. 12g**). By contrast, deletion of *ScMSN2* or *ScMSN4* did not alter FLC responses (**Supplementary Fig. 12g**). Deletion of ScAzf1, the *S. cerevisiae* ortholog of *C. albicans* orf19.173, which regulates carbon metabolism and cell wall organization[13], also increased FLC tolerance but did not affect susceptibility (**Supplementary Fig. 12g**). Thus, *C. albicans* Mnl1 and orf19.173, and their respective orthologs ScCom2 and ScAzf1, likely have conserved roles in FLC tolerance, whereas CaMsn4, ScMsn2, and ScMsn4 appear dispensable. In *S. cerevisiae*, the Msn2/4 TFs drive the general stress response, but in *C. albicans*, Msn4 does not, highlighting functional divergence that may be linked to its pathogenic lifestyle. Instead, both ScCom2 and Mnl1 repress FLC tolerance, revealing conserved roles in these responses.

**The FLC tolerance of the *mnl1* mutant is pH sensitive.** Mnl1 regulates the weak acid stress response in *C. albicans*, making *mnl1* cells more sensitive to acetic acid[11]. Mnl1-dependent genes contain STRE-like elements (SLEs) in their promoters, which drive transcription under weak acid stress, while Nrg1 antagonizes this response by competing for SLE binding (**Supplementary Fig. 13a**)[4]. We tested the effect of simultaneous exposure of cells to acetic acid and FLC; as expected, the *mnl1* deletion mutant from the TF KO library was more sensitive to acetic acid, consistent with previous findings[4]. By contrast, the effect of acetic acid was similar in WT and *orf19.173* cells (**Supplementary Fig. 13b**). Interestingly, when FLC and acetic acid were added to the same disk, the $RAD_{20}$ increased for *orf19.173*, indicating that this mutant is more susceptible than the WT to FLC in the presence of acetic acid (**Supplementary Fig. 13b**). For all three strains, tolerance was reduced when FLC was provided together with acetic acid, consistent with the prior observation that low pH reduces tolerance[2].

Next, we examined the effect of pH on FLC responses using disk diffusion assays on YPD plates at pH 4, 6, 8, and 10, relative to a pH of 6.5 in regular YPD plates. In WT cells, $RAD_{20}$ increased at pH 10, while tolerance remained low across pH conditions (**Supplementary Fig. 13c**). In *mnl1 cells*, $RAD_{20}$ increased gradually with pH, while tolerance decreased slightly at pH 4, and was cleared at pH 10 (**Supplementary Fig. 13c**). This suggests that Mnl1 maintains growth homeostasis in the face of changing pH.

We examined Mnl1 localization under different conditions, as its nuclear presence is essential for transcriptional activity. Using a strain that constitutively expresses mNeonGreen-Mnl1 from the *TDH3* promoter[14], we found that *Mnl1* remained cytoplasmic after 4 h in FLC (10 μg/ml) or acetic acid (0.25%) (**Supplementary Fig. 13d**). Transcriptomic analysis revealed that *MNL1* expression was unchanged by FLC ($log_2$-fold change = -0.009) but significantly upregulated by acetic acid ($log_2$-fold change = 0.491, p-adjust <0.05, **Supplementary Table 6**), reinforcing its role in the acetic acid response and consistent with the pH sensitivity of azole responses in *mnl1 cells*. However, since Mnl1 is neither transcriptionally induced nor nuclear-localized during FLC exposure, we posit that its high FLC tolerance likely stems from intrinsic cellular differences rather than a transcriptional response.

Our results show that the azole tolerance of *mnl1* is sensitive to changes in pH. The reduction in tolerance with pH suggests that Mnl1 coordinates a pH-dependent stress response. This observation aligns with the role of Mnl1 in the weak acid stress responses[4], and the pH sensitivity of the tolerance phenotype suggests that Mnl1 integrates multiple environmental signals to modulate FLC responses.

**Fine-tuning of the PKA pathway via cAMP modulates cell growth during FLC exposure.** To further understand how cAMP regulates FLC tolerance, we examined its role in colony growth, time of colony appearance, and intracellular azole-Cy5 accumulation. As an independent measure of FLC tolerance, we quantified colony formation at supraMIC FLC (10 μg/ml). Consistent with disk diffusion results observed with WT cells, H89 decreased tolerance 2.72-fold, and 8-bromo-cAMP increased it 3.73-fold (**Supplementary Fig. 10a**). In *mnl1*, H89 had no effect, while 8-bromo-cAMP slightly reduced tolerance (1.33-fold, **Supplementary Fig. 10a**). This suggests that further elevating cAMP in *mnl1*, which already has high basal cAMP levels, may impair its ability to tolerate FLC.

Next, we monitored the time of colony appearance with and without supraMIC FLC (10 µg/ml). While 8-bromo-cAMP did not significantly affect the ΔToA of *mnl1* cells, H89 delayed colony emergence in both WT and *mnl1 strain*s (**Supplementary Fig. 10b**), suggesting that reduced cAMP negatively impacts FLC tolerance. We also measured the effect of H89 and 8-bromo-cAMP on intracellular azole-Cy5 accumulation. In WT cells, H89 increased while 8-bromo-cAMP decreased azole-Cy5 levels (**Supplementary Fig. 10c**). By contrast, both compounds increased azole-Cy5 levels in *mnl1* cells by reducing the subpopulation with low accumulation and increasing the subpopulation with high azole-Cy5 (**Supplementary Fig. 10c-d**). This supports the idea that excessive cAMP can impair tolerance. Finally, propidium iodide (PI) staining revealed that H89 increased PI uptake in WT cells, indicating elevated cell permeability, a marker of cell viability (**Supplementary Fig. 10e**). In *mnl1* cells, both H89 and 8-bromo-cAMP increased PI staining, with 8-bromo-cAMP causing a stronger effect (**Supplementary Fig. 10e**). These results support the idea that the two strains differ in membrane architecture and that unbalanced cAMP levels can compromise cell viability and reduce FLC tolerance.

To further assess how cAMP impacts FLC responses, we performed checkerboard assays with FLC, H89, and 8-bromo-cAMP, measuring susceptibility ($MIC_{50}$, at 24 h) and tolerance (SMG, at 48 h) for WT and *mnl1* cells. Increasing H89 (0–100 µM) did not significantly alter susceptibility, and reduced tolerance in both strains, particularly at higher doses (**Supplementary Fig. 10f**). By contrast, low 8-bromo-cAMP concentrations enhanced resistance in *mnl1*, but at high doses (100 mM), susceptibility and tolerance were reduced (**Supplementary Fig. 10f**). To determine the direct impact of H89 and 8-bromo-cAMP on cell growth, we measured susceptibility and tolerance in the absence of FLC. H89 did not reach an inhibitory level at the concentrations tested, while both WT and *mnl1* had an $MIC_{50}$ (minimum inhibitory concentration of drug that leads to less than 50% growth relative to wells without drug) of 25 mM for 8-bromo-cAMP, indicating toxicity at high concentrations (**Supplementary Fig. 10g**). Consistent with the results obtained using FLC in combination with 8-bromo-cAMP, the *mnl1* mutant displayed reduced tolerance to 8-bromo-cAMP compared to the WT strain (**Supplementary Fig. 10h**). These results emphasize the importance of balanced cAMP concentrations for optimal azole responses (**Supplementary Fig. 10i**).

**Susceptibility TFs preferentially target transport and ergosterol genes, whereas tolerance TFs largely induce the expression of sphingolipid, ATP, and mitochondrial function genes.** We next compared the regulatory connections between transcription factor (TF) networks controlling susceptibility (9 TFs) or tolerance (31 TFs) and genes assigned to selected GO terms relevant to FLC adaptation and Mnl1-regulated processes. These included *C. albicans* GO terms such as ergosterol, transmembrane transport, sphingolipid, PKA, ATP, and mitochondrial organization (the list of genes belonging to each term is provided in **Supplementary Table 9**). Regulatory interactions between TFs and genes of interest were identified using the "Rank Genes by TF" module on PathoYeastract+[15]. Relative to the tolerance network of TFs, the susceptibility network displayed markedly more connections per TF to several GO terms, including transmembrane transport (45.00 connections/TF for susceptibility TFs vs. 23.68 for tolerance TFs), ergosterol (10.67 vs. 4.10), ATP (16.44 vs. 7.58), and mitochondrial organization (23.78 vs. 9.52) (**Supplementary Fig. 14a-b**, **Supplementary Table 10**). Sphingolipid genes showed relatively few connections with either network (1.44 vs 1.06). Analysis of the ratio of induced to repressed connections revealed further differences. In the susceptibility network, most GO terms had a balanced ratio near 1, except for sphingolipid (0.50), ATP (0.89), and mitochondrial organization (0.47) genes, which were predominantly repressed (**Supplementary Fig. 14a**, **Supplementary Table 10**). Ergosterol genes were similarly induced by TFs from both networks (1.38 vs 1.24). By contrast, the tolerance network showed an enrichment for the induction of sphingolipid genes (2.63) and a shift toward induction for mitochondrial organization (1.39) and ATP genes (1.37), suggesting that these processes might be more actively promoted in the tolerance state (**Supplementary Fig. 14b**, **Supplementary Table 10**). These findings indicate that susceptibility TFs are more broadly connected to membrane transport, mitochondrial, ATP, and ergosterol genes, but repress multiple lipid and mitochondrial genes. In contrast, tolerance TFs, while connected to fewer targets in these GO categories, are more likely to activate genes involved in sphingolipid metabolism, mitochondrial organization, and ATP production, potentially contributing to the establishment of a drug-tolerant physiological state.

**Validation of RNA-sequencing by RT-qPCR.** To validate the RNA-sequencing results, we performed RT-qPCR analysis of a subset of genes that showed differential expression in the *mnl1* mutant compared to the WT strain. The selected genes included *MIT1*, *RBR1*, *RTA2*, *SOD5*, and *SUR2*, which represent functions in stress responses and sphingolipid metabolism. RT-qPCR confirmed the general trends observed in the RNA-seq dataset. For example, *MIT1* and *SUR2* were consistently downregulated in *mnl1* cells relative to WT cells, whereas *RTA2* and *SOD5* were upregulated during growth on YPD and YPD+FLC (**Supplementary Figure 15**). These results support the robustness of our RNA-seq analysis and confirm that loss of *MNL1* leads to reproducible transcriptional changes in pathways linked to stress responses and lipid metabolism.

**Whole-genome sequencing of *mnl1* and WT strains.** To confirm that the phenotypes observed in the *mnl1* strain were not attributable to secondary mutations, we performed whole-genome sequencing of both the *mnl1* mutant and the isogenic WT parental strain from the TF KO collection. Comparative analysis revealed no additional single-nucleotide polymorphisms (SNPs), insertions, deletions, or structural variations between the WT and *mnl1* mutant genomes, aside from the expected targeted deletion at the *MNL1*. Copy number and heterozygosity profiles (**Supplementary Fig. 15a-b**) were nearly identical, indicating that the observed phenotypic differences described in this study are attributable solely to the loss of *MNL1* and not to secondary genomic alterations.

### Supplementary Figure Legends

**Supplementary Figure 1**: **Relationships between FLC susceptibility, tolerance, intracellular drug accumulation, time of colony appearance, and *in vivo* FLC tolerance. a,** Relative intracellular azole-Cy5 for TF KO mutants incubated with azole-Cy5 (1 µg/ml) for 24 h. Data is shown relative to the corresponding WT strain (red box). Experiments were performed with 2 to 5 biological replicates. **b,** Relative differences in the time of colony appearance (ΔToA) on FLC (10 µg/mL) plates relative to plates without FLC for 11 TF KO strains, then normalized to the ΔToA of the WT isolate. A total of 78-787 colonies across at least 3 biological replicates were examined for each strain. **c,** FNZ clears tolerance in the TF KO WT strain. Images show representative images of FLC disk diffusion assays (25 µg/disk) and quantification of FLC susceptibility ($RAD_{20}$) and tolerance ($FoG_{20}$) with and without FNZ (10 µg/ml). **d,** Fluphenazine (FNZ) increases intracellular azole-Cy5 accumulation in *C. albicans*. Mean fluorescence intensity levels for azole-Cy5 (1 µg/ml) intracellular accumulation after 24 h of exposure with and without FNZ (10 µg/ml). Asterisks show statistical comparisons between FLC-Cy5 and FLC-Cy5+FNZ conditions, nonparametric t-test, * $P$ <0.05, ** $P$ <0.01. **e,** Distribution of cell populations with different azole-Cy5 accumulation levels (low <200, medium 200-2000, high >2000). **f,** Survival of *G. mellonella* larvae infected with one of 24 different TF KO strains during 14-day systemic infections, followed by administration of PBS (control), FLC (1 mg/kg), or a combination of FLC (1 mg/kg) and FNZ (10 mg/kg). Survival was monitored daily, and values show the average daily survival for each condition relative to the number of infected larvae in each group. (FLC+FNZ)/FLC shows the impact of combination therapy relative to treatment with FLC alone. TFs in heatmaps are sorted according to the relative FLC tolerance of each TF KO strain ($FoG_{20}$). FLC susceptibility ($RAD_{20}$) levels are included for reference, also shown relative to the susceptibility of the corresponding WT.

All correlations were assessed using two-tailed nonparametric Spearman correlation coefficients, and significance was noted for $P<0.05$.

**Supplementary Figure 2: Transcription factor (TF) screening for altered fluconazole (FLC) drug responses using disk diffusion assays. a,** Changes in susceptibility ($RAD_{20}$) and tolerance ($FoG_{20}$) across the two TF libraries, shown relative to the corresponding WT controls. **b,** The number of TF mutants affecting susceptibility and tolerance across each library, using a ±20% cutoff difference relative to the WT strain. **c,** Overlap in the TFs that impact both susceptibility and tolerance for each library. **d,** Overlap between libraries in TFs that affect susceptibility versus those that specifically influence tolerance. **e-f,** Correlations between relative FLC susceptibility ($RAD_{20}$) and relative tolerance ($FoG_{20}$) for the TF KO (e) and TF OE (f) collections. **g-h,** Correlations between relative FLC tolerance ($FoG_{20}$) and relative doubling time (g) or relative lag time (h) for the TF KO mutants. All parameters are shown relative to the corresponding WT strains, marked in red.

**Supplementary Figure 3: 31 TFs cluster into two major tolerance regulatory modules. a,** Tolerance ($FoG_{20}$) levels for the 31 regulators identified in TF KO and OE screens relative to their WT parent. The panel shows $FoG_{20}$ levels when these regulators are either deleted (KO, X-axis) or overexpressed (OE, Y-axis). Tolerance enhancers and repressors are marked in the upper left quadrant and the lower right quadrant, respectively. **b-c,** The tolerance network includes two regulatory modules, a stress response module (b) and an unstressed log-phase growth module (c). The stress module includes alkaline, pH, weak acid stress, and drug responses. Interactions are marked according to the direction of the interaction: green, positive regulation; red, negative regulation; blue, unspecified.

**Supplementary Figure 4: A network of 9 TFs regulates FLC susceptibility.** Changes in susceptibility ($RAD_{20}$) levels upon deletion (KO) or overexpression (OE) of the 9 regulators identified in the two TF screens.

Susceptibility is shown relative to the corresponding WT controls. **b,** Regulatory relationships among the 9 susceptibility TFs based on gene expression or DNA binding evidence (PathoYeastract+), grouped according to the impact of overexpressing (OE) or deleting (KO) the respective TF on FLC susceptibility: orange, both KO and OE lead to increased susceptibility; turquoise, KO and OE lead to opposite effects on susceptibility. TFs Ace2, Hms1, Mrr1, Pho4, Stp4, Tec1, and Upc2 (highlighted in black boxes) show significant enrichment of interactions for the targets in this set of TFs relative to the rest of the *C. albicans* genome *(P* <0.005, hypergeometric test). Interactions are marked by arrows according to the direction of the interaction: green, positive regulation; red, negative regulation; blue, unspecified.

**Supplementary Figure 5**: **An independent set of mutant strains[62] reinforces the role of Mnl1 in FLC tolerance. a,** Representative disk diffusion assays images with FLC (25 μg/disk) for WT, deletion, and gene add-back strains. **b-c,** Susceptibility ($RAD_{20}$, b) and tolerance ($FoG_{20}$, c) for FLC disk diffusion assay of the strains in (a). **d,** Relative average intracellular azole-Cy5 levels for WT, deletion, and gene add-back strains. **e,** Population distribution profiles for intracellular azole-Cy5 levels for the strains in (d). For b-e, the vertical lines and the different colored bars indicate distinct groups of isogenic strains. In each group, comparisons were made against the first strain in the group (the WT and *msn4mnl1 URA3-/+* strain, respectively). Asterisks show statistical comparisons using nonparametric t-tests, * *P* <0.05, ** *P* <0.01.

**Supplementary Figure 6**: **Gene expression differences in ergosterol or drug transporters are not associated with the increased tolerance of *mnl1 cells*. a,** Susceptibility assays of WT and *mn1* cells with FLC and acetic acid (AcAc). The table shows susceptibility ($MIC_{50}$, measured at 24 h) and tolerance (SMG, measured at 48 h). The $MIC_{50}$ values were used in selecting treatment concentrations in RNA sequencing. **b,** PCA analysis of the transcriptomes of WT and *mnl1* cells grown in YPD (white), YPD+FLC (0.5 μg/ml, red), or YPD+AcAc (0.25%, blue). The scatter plot shows the relationships between 18 samples and PC1 and PC2 variables. **c,** Number of differentially expressed genes between WT and *mnl1* cells grown in YPD, YPD+FLC, or YPD+AcAc. **d,** Changes in gene expression levels for ergosterol and drug transporter genes when comparing *mnl1* and WT cells or FLC and YPD conditions. Values represent $log_2$-fold changes, asterisks denote p-adjust values <0.05. **e-f,** PCA analyses for ergosterol (e) and transmembrane transport (f) genes from the RNA-sequencing dataset.

**Supplementary Figure 7:** Summary of differentially expressed genes between WT and *mnl1* cells exposed to FLC (1 μg/ml) or acetic acid (AcAc, 0.25%) for 14 h. **a,** Volcano plots showing differentially expressed genes between WT and *mnl1* cells in YPD alone or YPD+FLC (noted as FLC) conditions, with the genes colored according to the p-adjust value and $log_2$-fold change. **b,** Heatmaps show differentially expressed genes between WT and *mnl1* cells that are common in: YPD, FLC, and AcAc; YPD and FLC; YPD and AcAc; or FLC and AcAc (marked in red boxes). The annotations on the right denote either experimentally established or predicted gene functions, inferred from homology with *S. cerevisiae*.

**Supplementary Figure 8**: **Sphingolipid biosynthesis genes are differentially expressed between WT and *mnl1* cells. a,** Schematic of the sphingolipid biosynthetic pathway in *C. albicans*. Enzymes colored in red are differentially regulated between WT and *mnl1 cells*, with the direction of the arrows indicating the direction of the regulation in the *mnl1* strain. **b,** Differentially expressed sphingolipid genes between WT and *mnl1 cells* and between YPD and FLC-grown cells. Heatmap values represent $log_2$-fold changes, asterisks denote p-adjust values <0.05. **c-d,** Lipid body integrated intensity (c) and maximum lipid body brightness (d) for WT and *mnl1 cells* grown in either YPD or FLC (128 μg/mL) and stained with BODIPY. **e,** Vacuole area for WT and *mnl1 cells* grown in either YPD or FLC (128 μg/mL) and stained with FM4-64. Asterisks show significant differences relative to the corresponding WT strain or between YPD and FLC conditions (nonparametric t tests, * *P* <0.05, ** *P* <0.01, *** *P* <0.001).

**Supplementary Figure 9**: **Mitochondrial activity analysis of WT and *mnl1 cells*. a,** Whole genome sequencing of WT and *mnl1* strains from the TF KO collection reveals differences in the copy numbers of the mitochondrial genome. The heatmap shows the relative chromosome copy numbers across the 8 nuclear chromosomes and the mitochondrial genome. **b,** Differential gene expression levels between WT and *mnl1 cells* for mitochondrial genes encoded in the nucleus and for genes encoding components of the PKA pathway. Heatmap values represent $log_2$-fold changes; asterisks indicate significant differences (p-adjust <0.05). **c,** Representative images of WT and *mnl1 cells* stained with MitoTracker Green and MitoTracker Red dyes with or without FLC exposure (128 µg/mL). Scale bar, 10 µm. **d,** Differential gene expression levels between WT and *mnl1 cells* for genes regulated by the PKA pathway, and ATP genes. Heatmap values represent $log_2$-fold changes; asterisks indicate significant differences (p-adjust <0.05).

**Supplementary Figure 10**: **The PKA pathway regulates the balance between growth and cell death during FLC exposure. a,** Effects of chemical modulation of the PKA/cAMP pathway with H89 (100 µM) and 8-Bromo-cAMP (8Br-cAMP, 100 mM) on WT and *mnl1* CFU growth on YPD plates with or without FLC (10 µg/mL). **b,** Differences in the time of appearance with and without FLC (10 µg/ml) for WT and *mnl1* colonies on YPD plates exposed to DMSO, the PKA inhibitor H89 (100 µM), or the cAMP analog 8-Bromo-cAMP (100 mM). **c-d,** H89 (100 µM) and 8-Bromo-cAMP (100 mM) alter azole-Cy5 intracellular accumulation (c) and azole-Cy5 population distribution (low <200, medium 200-2000, high >2000) (d). **e,** Relative PI fluorescence intensity levels for WT and *mnl1* strains treated with H89 (100 µM) or 8-Bromo-cAMP (100 mM) for 24 h. **f,** Checkerboard assays showing the effect of increasing concentrations of H89 (0 to 100 µM) and 8-Bromo-cAMP (0 to 100 mM) on FLC susceptibility and tolerance (SMG) for WT *and mnl1* strains. **g-h,** Broth microdilution assays with FLC, 8-Bromo-cAMP, and H89. $MIC_{50}$ (g) and SMG (h) levels are indicated for each drug, measured at 24 h and 48 h, respectively. **i,** Schematic illustrating the effect of cAMP concentrations on *C. albicans* cell growth in the presence of FLC. Growth is maximized at optimal cAMP levels, but cells do not grow when cAMP is depleted or found in excessive amounts. Asterisks show significant differences relative to DMSO-treated cells of the same genotype (* $P$ <0.05; ** $P$ <0.01, *** $P$ <0.001).

**Supplementary Figure 11: The impact of tolerance adjuvants on the high tolerance of *mnl1 cells*.** Representative images and diskImageR quantification of susceptibility ($RAD_{20}$) and tolerance ($FoG_{20}$) for WT and *mnl1 cells* on disk diffusion assays with FLC alone or FLC supplemented with FNZ (10 µg/ml), geldanamycin (0.5 µg/ml), or FK506 (0.5 µg/ml). The heatmap shows $RAD_{20}$ and $FoG_{20}$ levels relative to those of the WT strain with FLC alone. Asterisks show significant differences relative to the corresponding no drug condition (nonparametric t tests, * $P$ <0.05).

**Supplementary Figure 12: Additional *C. albicans* and *S. cerevisiae* regulators of FLC tolerance. a,** Representative images of disk diffusion assays of WT, *msn4*, and *orf19.173* TF KO strains using FLC (25 µg/disk). FLC susceptibility ($RAD_{20}$) and tolerance ($FoG_{20}$) based on disk diffusion assays for WT, *mnl1*, and *orf19.173*, shown relative to the WT strain. **b,** Growth curves of WT, *msn4*, and *orf19.173* TF KO strains with and without FLC (10 µg/mL). **c,** Azole-Cy5 fluorescence intensity levels for WT, *msn4,* and *orf19.173* measured at 24 h and shown relative to the WT strain. **d,** Population distribution for WT, *msn4*, and *orf19.173* based on azole-Cy5 accumulation levels (low <200, medium 200-2000, high >2000). **e-f,** Time course of azole-Cy5 (e) and rhodamine 6G (f) fluorescence intensity levels for WT and *orf19.173* measured at 2 h and 24 h. **g,** FLC disk diffusion assays with representative images and quantification of susceptibility ($RAD_{20}$) and tolerance ($FoG_{20}$) for *S. cerevisiae* WT, *com2*, *msn2*, *mns4,* and *azf1* deletion strains. Asterisks show significant differences relative to the corresponding WT strain (nonparametric t tests, * $P$ <0.05, ** $P$ <0.01).

**Supplementary Figure 13: The impact of weak acid stress and pH on FLC drug responses. a,** Weak acid stress regulators in *C. albicans* include Mnl1 and Nrg1 (adapted from[4]). **b,** Disk diffusion assays of WT, *mnl1,* and *orf19.173* deletion strains using FLC (25 µg/disk), acetic acid (10 µl/disk), or a combination of FLC + acetic acid (AcAc, 25 µg FLC + 10 µl glacial acetic acid). Susceptibility ($RAD_{20}$) and tolerance ($FoG_{20}$) levels are shown relative to the WT strain. Asterisks show significant differences relative to the WT strain (nonparametric t tests, * *P* <0.05). **c,** Representative images and quantification of susceptibility ($RAD_{20}$) and tolerance ($FoG_{20}$) from disk diffusion assays of WT and *mnl1* on YPD plates at different pH levels. Asterisks show significant differences relative to standard YPD plates at pH 6.5 (nonparametric t tests, * *P* <0.05, ** *P* <0.01, *** *P* <0.001, **** *P* <0.0001). **d,** Localization of a Neon-tagged version of Mnl1 during growth in YPD or after 3 h exposure to FLC (10 µg/ml), or acetic acid (AcAc, 0.25%). Scale bar, 10 µm.

**Supplemental Figure 14**: **Relationships between tolerance and susceptibility TF networks and genes within selected GO terms. a-b,** Connections between the 9 susceptibility TFs (a) or the 31 tolerance TFs (b) (shown in blue nodes) and genes from selected GO categories - sphingolipid metabolism, PKA pathway, ATP metabolism, and mitochondrial organization (shown in yellow nodes). Gene lists for each GO term are provided in **Supplementary Table 9**. The analysis includes only connections to genes within these GO terms and excludes self-regulatory links and TF–TF connections within each network. The regulatory networks were generated using PathoYeastract+. Connection types: red, repression; green, induction; black, unspecified.

**Supplemental Figure 15**: **Validation of RNA sequencing data using RT-qPCR.** Expression levels of representative genes (*MIT1*, *RBR1*, *RTA2*, *SOD5*, and *SUR2*) were measured by both RNA-seq and qRT-PCR, demonstrating consistency between the two methods. The *ACT1* gene was used for normalization of qRT-PCR results. For RNA sequencing data, values represent $log_2$-fold changes, asterisks denote p-adjust<0.05.

**Supplementary Figure 16: Whole-genome sequencing of *mnl1* and WT strains. a,** Copy number variation across the 8 *C. albicans* chromosomes for the WT and *mnl1* mutant strains. The drop in read depth corresponds to the targeted deletion at the *MNL1* locus on chromosome R (highlighted in red). **b,** Allele frequency plots for the WT and *mnl1* strains are shown across the 8 chromosomes as a proxy for genome heterozygosity.

### Supplementary Table Legends

**Supplementary Table 1: Strains used in this study.**

**Supplementary Table 2: Primers and plasmids used in this study.**

**Supplementary Table 3: Screening of the TF KO and OE collections.** Susceptibility ($RAD_{20}$) and tolerance ($FoG_{20}$) levels for each mutant are shown relative to the WT strain; SD = standard deviation. TFs that affect the two drug responses when both KO and OE are also listed.

**Supplementary Table 4: Significantly enriched GO biological process terms associated with susceptibility and tolerance TFs.** The analysis was performed on the 9 susceptibility regulators and the 31 tolerance regulators. Terms were identified with the Gene Ontology Term Finder tool (*Candida* Genome Database) and selected based on significance (*P* <0.05), fold enrichment (>10X), and false discovery rate (FDR ≤0.01).

**Supplementary Table 5: Enriched interactions with other TFs in the susceptibility and tolerance networks.** Documented gene regulations between the TFs within each network, filtered by DNA binding and expression evidence, and based on significance (*P* <0.005, hypergeometric test, PathoYeastract+).

**Supplementary Table 6: Differentially expressed genes between WT and *mnl1 cells*.** Significant gene expression changes between WT and *mnl1 cells* grown in YPD, YPD+FLC (0.5 µg/ml), or YPD+acetic acid

(AcAc, 0.25%). Columns E-G indicate statistical significance for comparisons between WT and *mnl1* cells (1, p-adjust <=0.05; 0, p-adjust >0.05).

**Supplementary Table 7: Significant GO biological process terms enriched in the transcriptome of *mnl1* relative to WT cells.** GO enrichment was assessed using differentially expressed genes between WT and *mnl1 cells* grown in YPD or YPD+FLC (0.5 µg/ml). Terms were identified with the Gene Ontology Term Finder tool (*Candida* Genome Database) and selected based on significance (*P* <0.05), fold enrichment (>5X), and false discovery rate (FDR ≤0.01).

**Supplementary Table 8: Quantification of sphingolipid species by mass spectrometry analysis.** Values show lipid amounts relative to Pi (pmol of lipid/nmol of Pi).

**Supplementary Table 9: *C. albicans* genes within the GO terms used for PCA analyses.** Terms include ergosterol, transmembrane transport, sphingolipid, PKA, ATP, and mitochondrial organization; the genes belonging to each term were extracted from the *Candida* Genome Database.

**Supplementary Table 10: Summary of regulatory interactions (induced, repressed, and unspecified) between tolerance or susceptibility TF networks and genes within selected GO terms.** Interactions between TFs and gene sets were analyzed using PathoYeastract+. Only interactions with genes belonging to the respective GO terms were included (see **Supplementary Table 8**), while self-regulatory interactions and TF-TF connections within each network were excluded.

**Supplementary Table 11.** Growth parameters for mutants in the TF KO collection. Lag time and doubling time levels for each mutant are shown relative to the WT strain.

### Supplementary Figure 1.

**a**

| | $RAD_{20}$ | $FoG_{20}$ | Cy5 |
|---|---|---|---|
| *SWI4* | 1.22 | 0.34 | 2.71 |
| *IRO1* | 1.17 | 0.38 | 1.30 |
| *CZF1* | 1.28 | 0.43 | 2.26 |
| *UME7* | 1.18 | 0.53 | 1.19 |
| orf19.2476 | 1.29 | 0.61 | 2.04 |
| *CSR1* | 1.11 | 0.64 | 1.89 |
| *CAS5* | 1.08 | 0.64 | 1.37 |
| *GRF10* | 1.16 | 0.64 | 1.36 |
| *ROB1* | 1.11 | 0.67 | 1.38 |
| orf19.1150 | 0.96 | 0.68 | 1.31 |
| orf19.3928 | 1.20 | 0.69 | 1.77 |
| *UPC2* | 1.78 | 0.69 | 1.47 |
| *ZFU2* | 1.14 | 0.70 | 1.40 |
| *YNG1* | 1.11 | 0.71 | 1.53 |
| *MIG2* | 1.13 | 0.72 | 1.32 |
| *ZCF19* | 1.28 | 0.72 | 1.06 |
| *CTF1* | 1.34 | 0.74 | 1.62 |
| *MRR1* | 1.26 | 0.77 | 1.85 |
| *CRZ1* | 1.13 | 0.77 | 1.74 |
| *STB5* | 1.20 | 0.78 | 1.56 |
| *ZCF17* | 1.23 | 0.79 | 1.14 |
| *ISW2* | 0.92 | 0.80 | 1.50 |
| *TRY5* | 1.23 | 0.82 | 1.75 |
| *RPN4* | 1.42 | 0.83 | 1.92 |
| *SEF1* | 1.19 | 0.88 | 1.08 |
| *FCR1* | 1.11 | 0.89 | 0.89 |
| *HAP43* | 1.22 | 0.89 | 1.12 |
| *GLN3* | 1.32 | 0.91 | 1.18 |
| *BAS1* | 1.20 | 0.94 | 1.18 |
| WT | 1.00 | 1.00 | 1.00 |
| *WAR1* | 1.20 | 1.08 | 1.52 |
| *MRR2* | 1.26 | 1.10 | 1.01 |
| *MSN4* | 1.17 | 1.13 | 1.11 |
| *AHR1* | 1.18 | 1.20 | 1.41 |
| *ZCF2* | 0.98 | 1.23 | 0.55 |
| *IRF1* | 1.17 | 1.24 | 1.19 |
| orf19.173 | 1.16 | 1.25 | 0.72 |
| *RLM1* | 1.06 | 1.43 | 0.57 |
| *ZFU3* | 1.00 | 1.48 | 1.07 |
| *ARG81* | 1.07 | 1.54 | 0.44 |
| *FCR3* | 1.07 | 1.59 | 0.51 |
| *DAL81* | 1.09 | 1.64 | 0.48 |
| *CPH2* | 1.00 | 1.65 | 0.37 |
| *FGR15* | 1.13 | 1.67 | 0.78 |
| *STP4* | 1.26 | 1.69 | 0.79 |
| *RAP1* | 1.19 | 1.96 | 0.13 |
| *BCR1* | 1.19 | 1.98 | 0.49 |
| *MNL1* | 1.16 | 2.18 | 0.51 |
| *RIM101* | 1.07 | 2.61 | 0.56 |
| r | 0.425 | -0.784 | |
| *P* | 0.002 | <0.001 | |

**b**

| | $RAD_{20}$ | $FoG_{20}$ | ΔToA |
|---|---|---|---|
| *CSR1* | 1.11 | 0.64 | 2.16 |
| *MIG2* | 1.13 | 0.72 | 2.04 |
| *MRR1* | 1.26 | 0.77 | 1.56 |
| *CRZ1* | 1.13 | 0.77 | 2.08 |
| *FCR1* | 1.11 | 0.89 | 1.51 |
| WT | 1.00 | 1.00 | 1.00 |
| *WAR1* | 1.20 | 1.08 | 1.89 |
| *MSN4* | 1.17 | 1.13 | 1.19 |
| *RLM1* | 1.06 | 1.43 | 1.47 |
| *RAP1* | 1.19 | 1.96 | 1.25 |
| *MNL1* | 1.16 | 2.18 | 0.67 |
| *RIM101* | 1.07 | 2.61 | 0.54 |
| r | 0.196 | -0.846 | |
| *P* | 0.543 | 0.001 | |

**c**

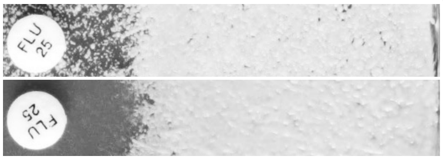


| | $RAD_{20}$ | $FoG_{20}$ |
|---|---|---|
| FLC | 14 | 0.39 |
| FLC + FNZ | 13 | 0.2 |

**d**

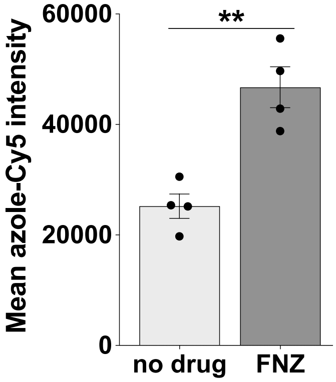


**e**

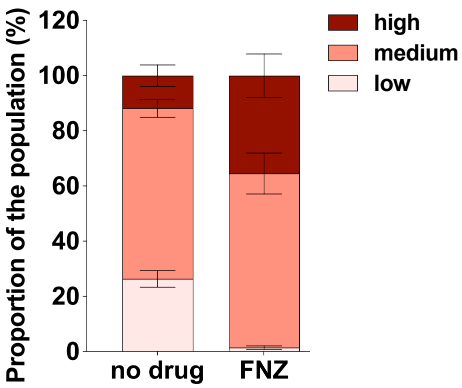


| | |
|---|---|
| high | * |
| medium | ns |
| low | ** |

**f**

| | $RAD_{20}$ | $FoG_{20}$ | Average daily survival (%) PBS | FLC | FLC+FNZ | FLC+FNZ / FLC |
|---|---|---|---|---|---|---|
| no infection | | | 82.02 | 79.05 | 76.49 | 0.97 |
| *CZF1* | 1.28 | 0.43 | 29.76 | 36.01 | 32.74 | 0.91 |
| *ZCF35* | 1.00 | 0.61 | 20.63 | 49.21 | 50.00 | 1.02 |
| orf19.2476 | 1.29 | 0.61 | 18.15 | 30.06 | 29.17 | 0.97 |
| *CSR1* | 1.11 | 0.64 | 31.94 | 51.19 | 55.56 | 1.09 |
| *GRF10* | 1.16 | 0.64 | 28.87 | 39.29 | 36.31 | 0.92 |
| *ROB1* | 1.11 | 0.67 | 7.74 | 31.85 | 33.33 | 1.05 |
| *UPC2* | 1.78 | 0.69 | 18.45 | 33.93 | 33.63 | 0.99 |
| *ZFU2* | 1.14 | 0.70 | 13.49 | 39.88 | 35.91 | 0.90 |
| *MIG2* | 1.13 | 0.72 | 23.61 | 40.48 | 42.46 | 1.05 |
| *CTF1* | 1.34 | 0.74 | 25.20 | 36.11 | 38.89 | 1.08 |
| *CRZ1* | 1.13 | 0.77 | 35.12 | 38.39 | 38.10 | 0.99 |
| *RPN4* | 1.42 | 0.83 | 33.13 | 56.75 | 58.73 | 1.04 |
| *FCR1* | 1.11 | 0.89 | 26.19 | 45.04 | 47.02 | 1.04 |
| *HAP43* | 1.22 | 0.89 | 28.69 | 32.50 | 37.40 | 1.15 |
| WT | 1.00 | 1.00 | 25.82 | 49.03 | 54.54 | 1.11 |
| *WAR1* | 1.20 | 1.08 | 13.69 | 48.21 | 58.33 | 1.21 |
| *MSN4* | 1.17 | 1.13 | 15.48 | 44.64 | 45.04 | 1.01 |
| *AHR1* | 1.18 | 1.20 | 12.50 | 35.71 | 45.24 | 1.27 |
| orf19.173 | 1.16 | 1.25 | 10.42 | 23.81 | 29.17 | 1.23 |
| *RLM1* | 1.06 | 1.43 | 25.00 | 54.37 | 61.19 | 1.13 |
| *DAL81* | 1.09 | 1.64 | 11.90 | 52.38 | 61.01 | 1.16 |
| *STP4* | 1.26 | 1.69 | 18.65 | 39.48 | 49.40 | 1.25 |
| *RAP1* | 1.19 | 1.96 | 12.30 | 31.15 | 36.51 | 1.17 |
| *MNL1* | 1.16 | 2.18 | 19.79 | 45.09 | 57.44 | 1.27 |
| *RIM101* | 1.07 | 2.61 | 35.52 | 44.64 | 57.54 | 1.29 |
| r | -0.084 | 0.816 | | | | |
| *P* | 0.691 | <0.001 | | | | |

### Supplementary Figure 2.

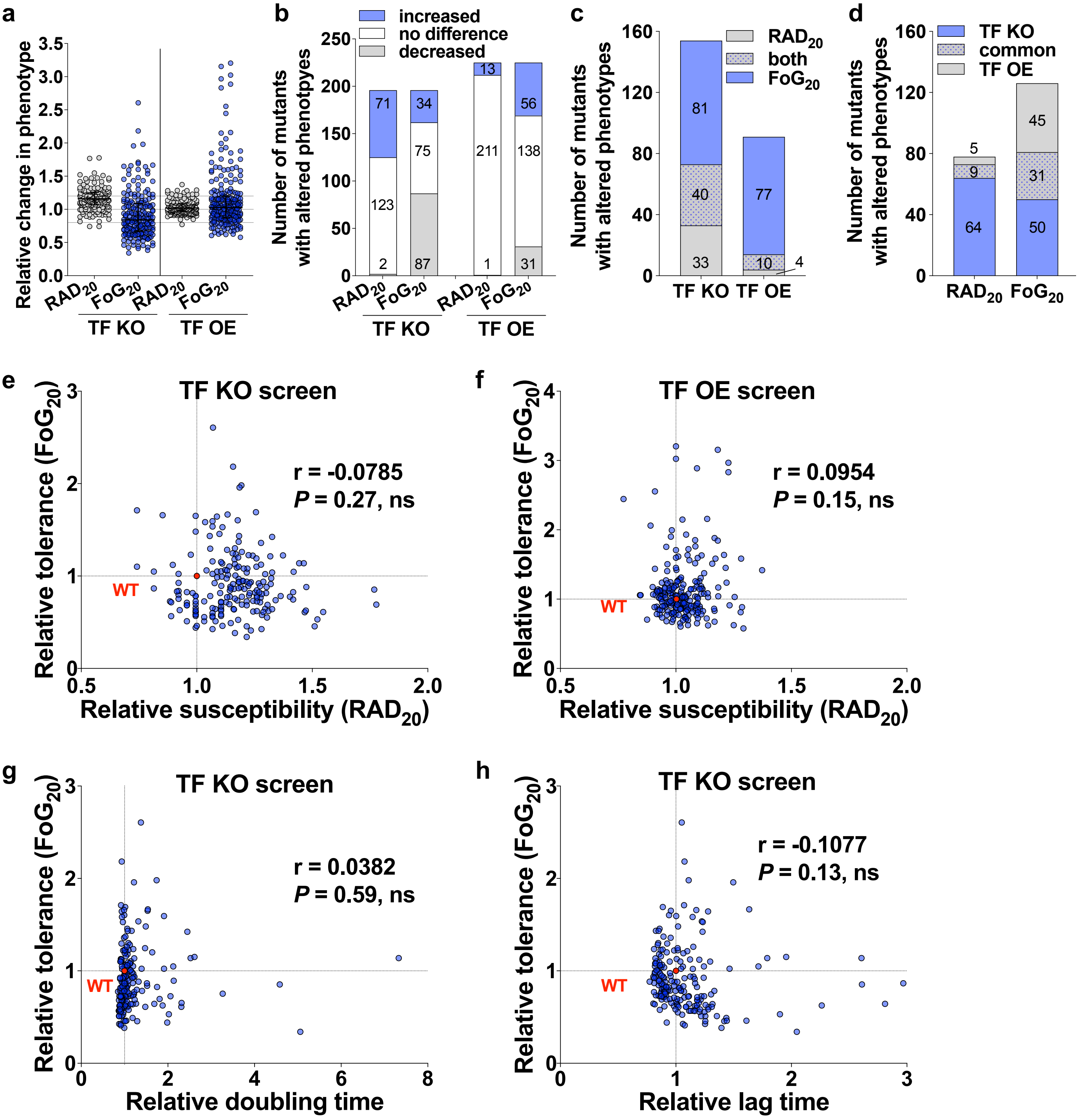

**Supplementary Figure 3.**

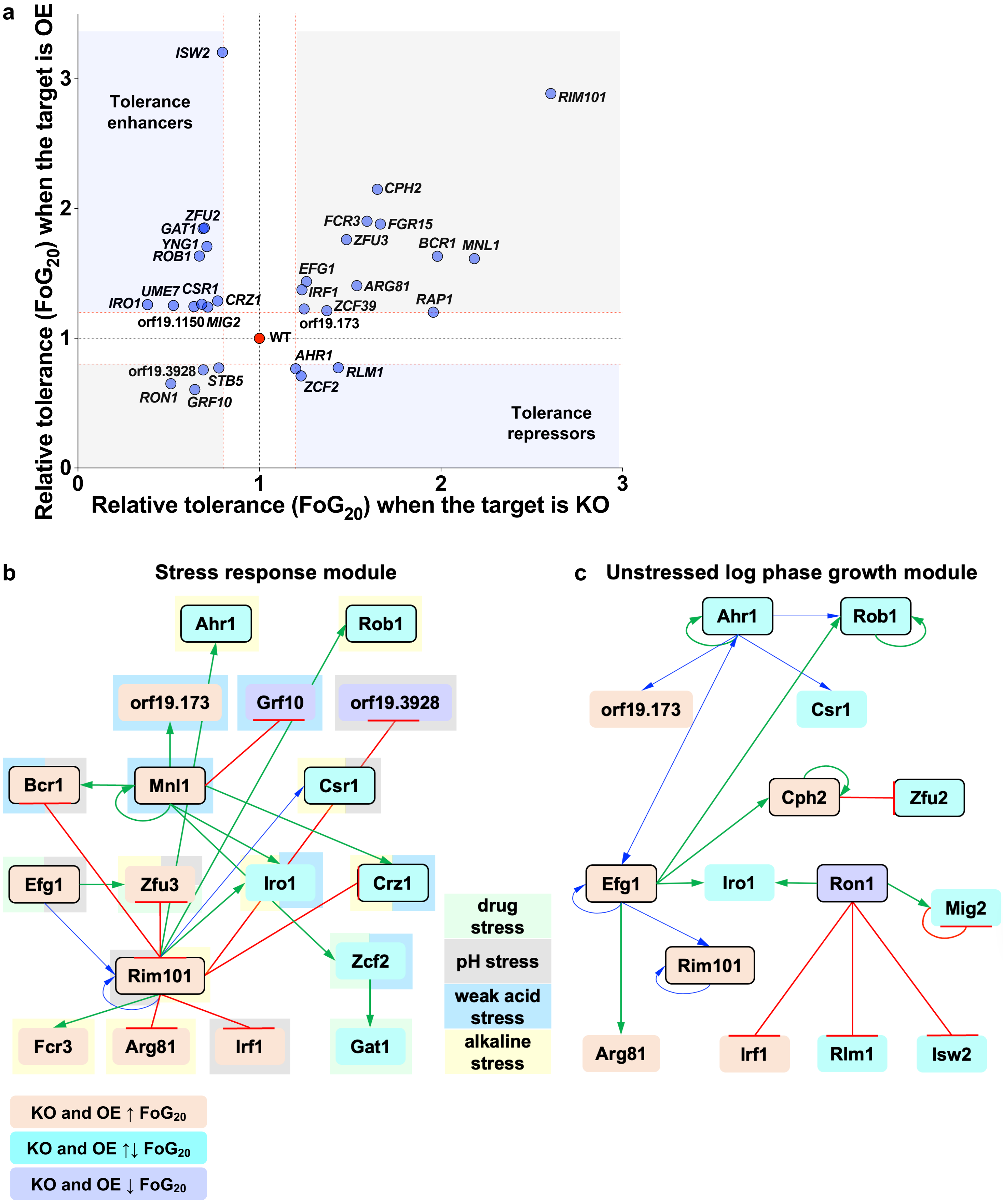

**Supplementary Figure 4.**

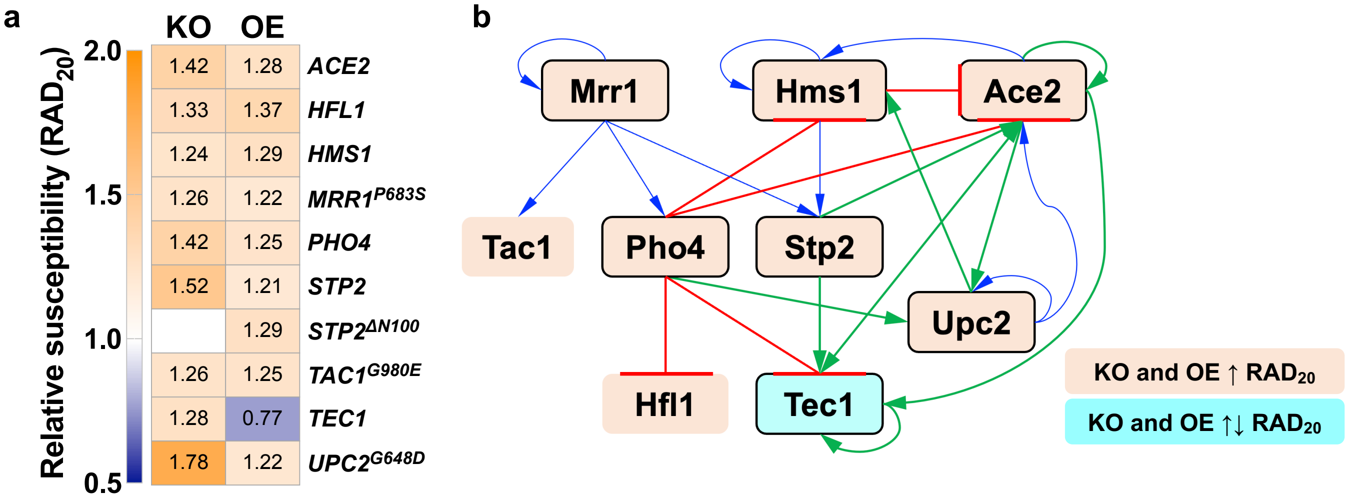

**Supplementary Figure 5.**

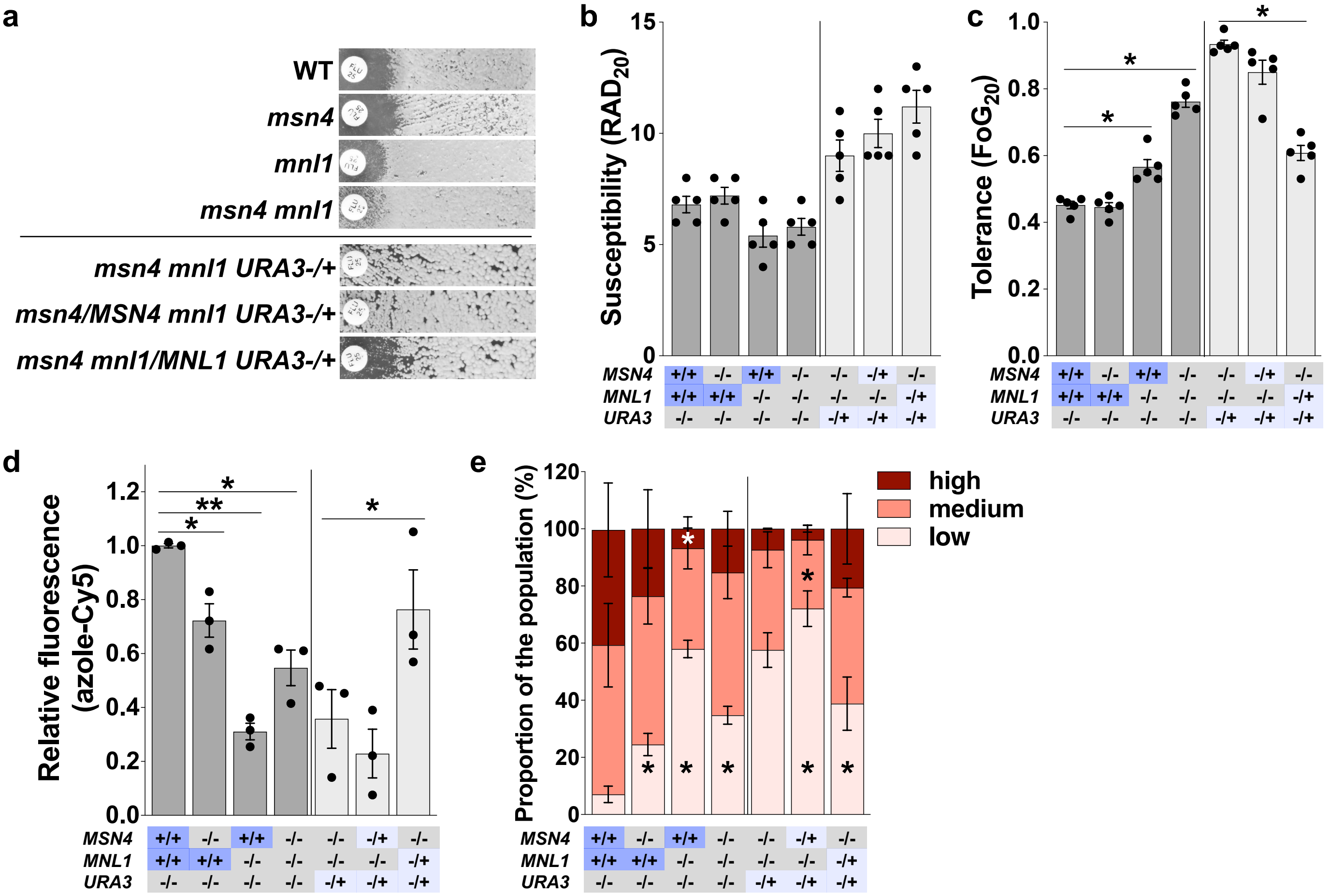

### Supplementary Figure 6.

**a**

| Condition | WT | | *mnl1* | |
|---|---|---|---|---|
| | $MIC_{50}$ | SMG | $MIC_{50}$ | SMG |
| FLC | 1 | 0.58 | 1 | 0.88 |
| AcAc | 0.5 | 0.09 | 0.25 | 0.20 |

**b**

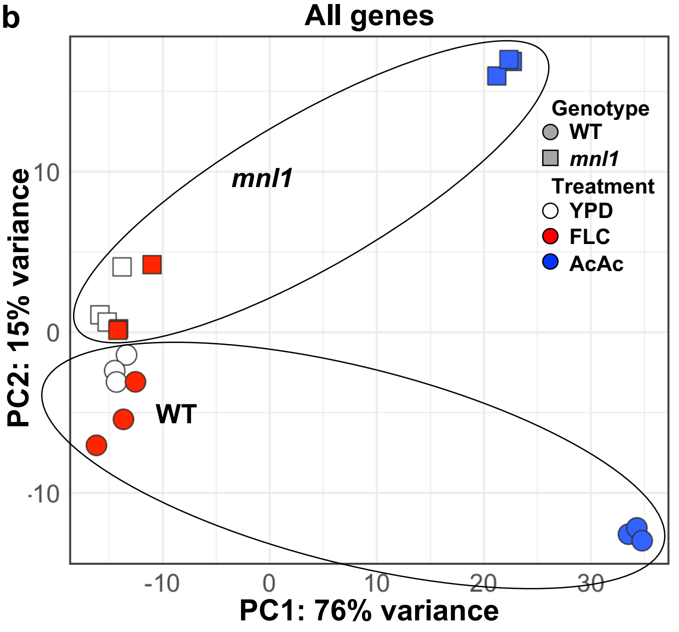


**c**

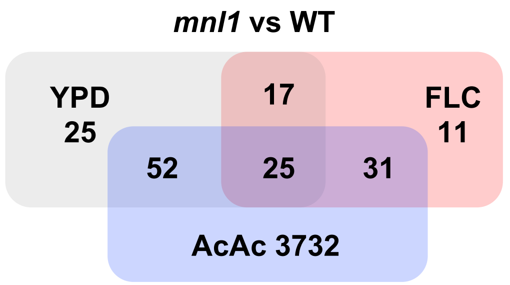


**d**

| | | *mnl1* vs WT | | FLC vs YPD | |
|---|---|---|---|---|---|
| | | YPD | FLC | WT | *mnl1* |
| ergosterol biosynthesis genes | *UPC2* | -0.24 | -0.13 | 0.45 * | 0.56 * |
| | *ERG1* | -0.10 | 0.00 | 0.74 * | 0.84 * |
| | *ERG2* | -0.07 | 0.00 | 0.65 * | 0.73 * |
| | *ERG3* | -0.38 | 0.06 | 1.89 * | 2.33 * |
| | *ERG4* | -0.10 | 0.06 | 0.58 * | 0.73 * |
| | *ERG5* | -0.11 | -0.04 | 1.02 * | 1.09 * |
| | *ERG6* | -0.05 | -0.47 | 2.15 * | 1.74 * |
| | *ERG7* | -0.07 | -0.06 | 0.44 * | 0.46 * |
| | *ERG8* | 0.03 | 0.09 | -0.06 | 0.00 |
| | *ERG9* | 0.12 | 0.18 | 0.42 * | 0.48 * |
| | *ERG10* | -0.07 | 0.02 | 0.68 * | 0.76 * |
| | *ERG11* | -0.16 | -0.02 | 1.09 * | 1.23 * |
| | *ERG12* | -0.02 | 0.09 | -0.15 | -0.03 |
| | *ERG13* | -0.12 | 0.03 | 0.60 * | 0.75 * |
| | *ERG20* | 0.05 | 0.03 | -0.09 | -0.10 |
| | *ERG24* | -0.06 | -0.19 | 0.95 * | 0.82 * |
| | *ERG25* | 0.12 | -0.03 | 0.84 * | 0.69 * |
| | *ERG26* | -0.13 | -0.09 | 0.43 * | 0.47 * |
| | *ERG27* | 0.14 | -0.11 | 0.43 * | 0.19 |
| | *ERG28* | -0.06 | 0.02 | 0.36 | 0.43 |
| | *ERG251* | 0.05 | -0.06 | 0.95 * | 0.84 * |
| ABC and MDR transporter genes | *MRR1* | -0.04 | -0.16 | -0.02 | -0.15 |
| | *MRR2* | 0.07 | 0.04 | 0.00 | -0.02 |
| | *ADP1* | 0.07 | 0.11 | -0.07 | -0.03 |
| | *ATM1* | -0.04 | -0.04 | -0.19 | -0.19 |
| | *CDR1* | 0.19 | 0.26 | -0.28 | -0.20 |
| | *CDR11* | 0.22 | 0.36 | -0.07 | 0.07 |
| | *CDR2* | 0.03 | 0.28 | -0.19 | 0.06 |
| | *CDR3* | -0.07 | 0.08 | 0.09 | 0.24 |
| | *CDR4* | -0.06 | 0.19 | -0.05 | 0.20 |
| | *FLU1* | 0.12 | 0.10 | -0.35 | -0.37 |
| | *MDL1* | -0.05 | 0.06 | -0.11 | 0.00 |
| | *MDL2* | 0.04 | 0.00 | -0.08 | -0.12 |
| | *MDR1* | 0.30 | 0.62 | 0.11 | 0.44 |
| | orf19.3120 | 1.14 | 0.06 | 1.11 | 0.02 |
| | orf19.6382 | -0.02 | -0.03 | 0.03 | 0.02 |
| | *ROA1* | -0.38 | -0.47 | 0.43 | 0.34 |
| | *SNQ2* | 0.00 | 0.16 | -0.24 | -0.07 |
| | *TAC1* | 0.15 | 0.08 | -0.17 | -0.23 |
| | *YCF1* | -0.06 | -0.03 | -0.10 | -0.07 |
| | *YOR1* | 0.17 | 0.30 | -0.33 | -0.20 |

$\log_2$ fold change: -2 0 2

**e**

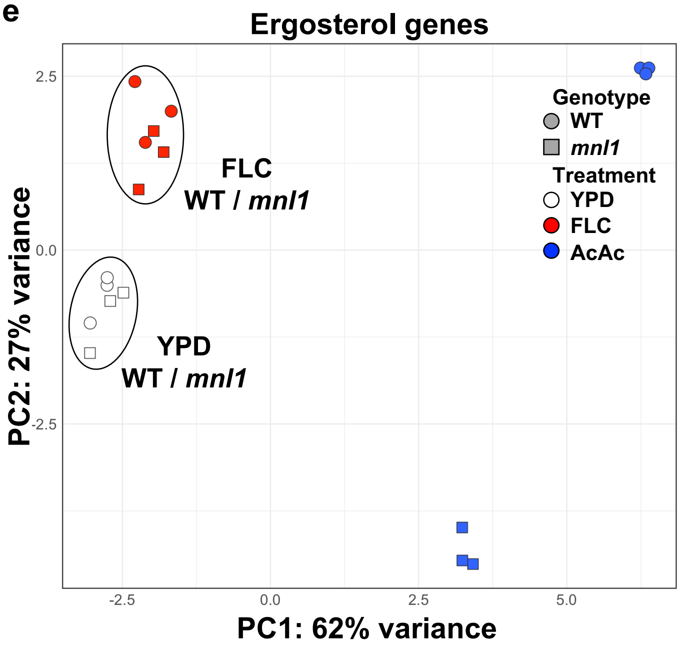


**f**

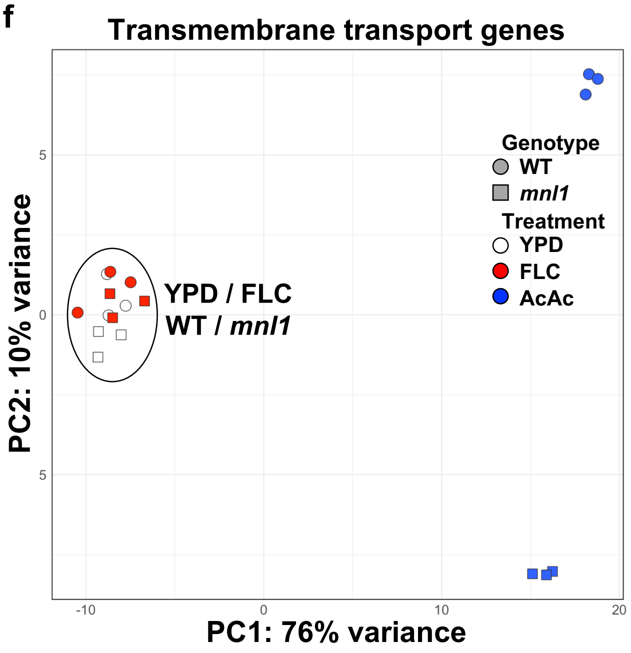

### Supplementary Figure 7.

**a**

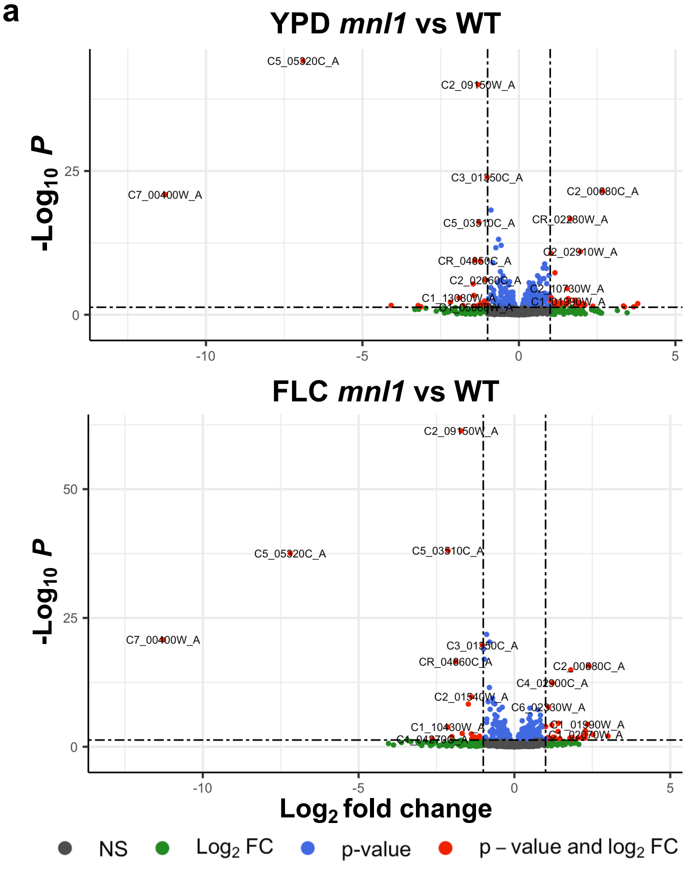


**b**

***mnl1* vs WT** — common in YPD, FLC and AcAc

| | YPD | FLC | AcAc | |
|---|---|---|---|---|
| ***MIT1*** | -1.30 | -1.71 | -1.00 | lipid homeostasis |
| ***orf19.6660*** | -1.27 | -2.13 | 0.94 | |
| ***URA3*** | -1.02 | -1.03 | -0.82 | metabolism |
| ***IRO1*** | -0.89 | -0.99 | -1.66 | iron homeostasis |
| ***RSD1*** | -0.83 | -0.78 | 1.93 | lipid homeostasis |
| ***LAG1*** | -0.74 | -0.56 | -0.57 | lipid homeostasis |
| ***SUR2*** | -0.65 | -0.89 | -1.34 | lipid homeostasis |
| ***AIF1*** | -0.59 | -0.41 | 1.25 | mitochondria |
| ***IML2*** | -0.57 | -0.80 | -1.91 | stress response |
| ***JSN1*** | -0.51 | -0.63 | -0.41 | mitochondria |
| ***orf19.5754*** | -0.43 | -0.50 | 0.72 | ion homeostasis |
| ***orf19.6601*** | -0.41 | -0.97 | -0.29 | |
| ***PRB1*** | -0.39 | -0.34 | -0.89 | metabolism |
| ***BZZ1*** | -0.30 | -0.31 | -0.46 | cytoskeleton organization |
| ***SIP5*** | -0.24 | -0.39 | -0.29 | stress response |
| ***BMT6*** | 0.29 | 0.32 | -1.30 | cell wall integrity |
| ***IQG1*** | 0.47 | 0.62 | -0.30 | cell cycle |
| ***RHD1*** | 0.55 | 0.60 | -1.86 | cell wall integrity |
| ***orf19.3475*** | 0.63 | 1.09 | 0.74 | stress response |
| ***orf19.3219*** | 0.68 | 0.69 | 1.09 | ATP |
| ***RNH1*** | 0.77 | 0.48 | 0.37 | |
| ***CTA26*** | 0.80 | 0.82 | -0.69 | stress response |
| ***CRH11*** | 0.88 | 1.22 | -0.57 | cell wall integrity |
| ***orf19.2125*** | 1.02 | 0.83 | 0.39 | |
| ***PGA23*** | 1.64 | 1.81 | 1.23 | PKA regulated |

common in YPD and FLC

| | YPD | FLC | AcAc | |
|---|---|---|---|---|
| ***orf19.6308*** | -1.41 | -1.48 | -0.45 | |
| ***orf19.6307*** | -1.23 | -1.89 | 0.15 | |
| ***orf19.1461*** | -0.89 | -1.38 | -0.17 | stress response |
| ***URO99*** | -0.64 | -0.51 | 0.23 | metabolism |
| ***PMU2*** | -0.58 | -0.68 | 0.00 | metabolism |
| ***ASM3*** | -0.56 | -0.63 | 0.06 | lipid homeostasis |
| ***orf19.4883*** | -0.56 | -0.84 | -0.24 | stress response |
| ***PRP40*** | -0.44 | -0.67 | 0.20 | splicing |
| ***AUR1*** | -0.42 | -0.80 | -0.03 | lipid homeostasis |
| ***orf19.5370*** | 0.37 | 0.51 | -0.10 | vacuole |
| ***RTA4*** | 0.53 | 0.50 | -0.15 | lipid homeostasis |
| ***FGR29*** | 0.63 | 0.77 | -0.29 | stress response |
| ***orf19.5468*** | 0.82 | 1.00 | 0.03 | |
| ***orf19.270*** | 0.90 | 1.20 | 0.14 | |
| ***RTA2*** | 1.15 | 0.86 | -0.46 | lipid homeostasis |
| ***orf19.5814.1*** | 1.96 | 1.41 | 0.52 | |
| ***SOD5*** | 2.68 | 2.39 | 0.03 | stress response |

log2 fold change: -3, 0, 3

***mnl1* vs WT** — common in YPD and AcAc

| | YPD | FLC | AcAc | |
|---|---|---|---|---|
| ***AGA1*** | -1.46 | -0.47 | 2.31 | |
| ***DAL1*** | -0.82 | -0.06 | -0.51 | metabolism |
| ***orf19.36.1*** | -0.80 | -0.71 | 0.89 | |
| ***GOR1*** | -0.68 | -0.18 | -0.82 | metabolism |
| ***orf19.2968*** | -0.60 | -0.24 | 1.82 | |
| ***orf19.2244*** | -0.58 | -0.55 | -0.47 | metabolism |
| ***MNN42*** | -0.56 | -0.41 | 0.30 | ATP |
| ***orf19.1449*** | -0.49 | -0.20 | 3.86 | stress response |
| ***SAD2*** | -0.42 | -0.14 | 1.14 | metabolism |
| ***MDH1-3*** | -0.37 | -0.27 | -0.47 | metabolism |
| ***orf19.3139*** | -0.34 | -0.21 | 0.18 | stress response |
| ***PTC6*** | -0.33 | -0.25 | -0.44 | mitochondria |
| ***TPS1*** | -0.27 | -0.12 | -0.41 | stress response |
| ***ATG15*** | -0.26 | -0.21 | -0.96 | lipid homeostasis |
| ***HEM15*** | -0.19 | -0.01 | -0.50 | stress response |
| ***CTA3*** | -0.19 | -0.16 | -0.52 | transport |
| ***SVL3*** | 0.26 | 0.13 | 1.03 | transport |
| ***orf19.3215*** | 0.26 | 0.23 | -0.57 | stress response |
| ***HAP43*** | 0.28 | 0.06 | 1.14 | ion homeostasis |
| ***PNG2*** | 0.30 | 0.26 | 0.20 | cell wall integrity |
| ***YJU3*** | 0.30 | 0.14 | -0.21 | lipid homeostasis |
| ***MUC1*** | 0.33 | 0.21 | -0.30 | filamentation |
| ***OPY2*** | 0.34 | 0.25 | 0.82 | cell wall integrity |
| ***ALS2*** | 0.35 | 0.25 | -2.95 | adhesion |
| ***orf19.90*** | 0.36 | 0.16 | -1.01 | lipid homeostasis |
| ***DFI1*** | 0.36 | 0.17 | 1.09 | cell wall integrity |
| ***orf19.1350*** | 0.36 | 0.27 | 1.18 | stress response |
| ***AXL2*** | 0.39 | 0.33 | 1.61 | cell division |
| ***orf19.3737*** | 0.39 | 0.18 | -0.90 | vacuole |
| ***ECM22*** | 0.40 | 0.05 | -0.66 | |
| ***orf19.7210*** | 0.42 | 0.13 | -0.64 | |
| ***DRS2*** | 0.44 | 0.23 | -0.87 | lipid homeostasis |
| ***PMA1*** | 0.45 | 0.26 | 0.33 | ATP |
| ***orf19.374*** | 0.46 | 0.28 | -0.51 | transport |
| ***CEK1*** | 0.46 | 0.22 | 0.95 | stress response |
| ***FRK1*** | 0.48 | 0.30 | 0.25 | biofilm formation |
| ***PIR1*** | 0.48 | 0.36 | -0.91 | cell wall integrity |
| ***AKR1*** | 0.48 | 0.14 | -0.70 | |
| ***BMT5*** | 0.49 | 0.35 | -0.86 | cell wall integrity |
| ***orf19.3264.1*** | 0.50 | 0.07 | -1.10 | |
| ***CRZ1*** | 0.52 | 0.21 | -0.46 | stress response |
| ***CDC20*** | 0.55 | 0.44 | -0.54 | cell cycle |
| ***PHR1*** | 0.59 | 0.25 | 0.58 | cell wall integrity |
| ***SKN1*** | 0.59 | 0.44 | 0.72 | cell wall integrity |
| ***BMT3*** | 0.60 | 0.34 | -0.78 | cell wall integrity |
| ***PMC1*** | 0.61 | 0.37 | -0.43 | ATP |
| ***CCN1*** | 0.65 | 0.10 | 1.07 | cell cycle |
| ***orf19.9*** | 0.68 | 0.66 | -0.44 | |
| ***EXG2*** | 0.77 | -0.17 | -1.07 | cell wall integrity |
| ***orf19.3694*** | 0.82 | 0.42 | 0.66 | |
| ***orf19.3615*** | 0.89 | 0.18 | -1.01 | stress response |
| ***BMT4*** | 0.96 | 0.65 | -0.59 | cell wall integrity |

common in FLC and AcAc

| | YPD | FLC | AcAc | |
|---|---|---|---|---|
| ***GDA100*** | -0.52 | -0.89 | 2.16 | metabolism |
| ***ASR1*** | -0.53 | -0.89 | 0.99 | stress response |
| ***orf19.6084*** | -0.24 | -0.70 | 2.72 | |
| ***CBF1*** | -0.26 | -0.61 | -0.98 | metabolism |
| ***AGP2*** | -0.12 | -0.57 | 0.74 | metabolism |
| ***HHF22*** | -0.15 | -0.53 | -2.57 | chromatin assembly |
| ***orf19.5070*** | -0.07 | -0.48 | 1.12 | stress response |
| ***HTB1*** | -0.16 | -0.45 | -2.24 | chromatin assembly |
| ***STP4*** | -0.28 | -0.45 | 2.35 | stress response |
| ***ASR2*** | -0.29 | -0.43 | 0.86 | stress response |
| ***BUD14*** | -0.18 | -0.43 | 0.45 | cytoskeleton organization |
| ***AUT7*** | -0.20 | -0.42 | -1.04 | mitochondria |
| ***orf19.3659*** | -0.30 | -0.40 | -0.58 | transcription regulator |
| ***orf19.225*** | -0.20 | -0.32 | -0.37 | |
| ***CDC34*** | -0.18 | -0.29 | -0.68 | cell cycle |
| ***BBC1*** | -0.16 | -0.24 | -0.53 | cytoskeleton organization |
| ***BOI2*** | 0.18 | 0.24 | 0.69 | cytoskeleton organization |
| ***GSF2*** | 0.13 | 0.25 | -0.65 | metabolism |
| ***CAS5*** | 0.19 | 0.31 | 0.65 | cell wall integrity |
| ***TPM1*** | 0.21 | 0.35 | 0.91 | |
| ***orf19.3080*** | 0.12 | 0.38 | 0.49 | ATP |
| ***DCK1*** | -0.06 | 0.38 | 1.23 | citokinesis |
| ***ZRT2*** | 0.30 | 0.45 | -1.24 | ion homeostasis |
| ***GIT2*** | 0.18 | 0.47 | 0.75 | transport |
| ***CHS8*** | 0.28 | 0.47 | -0.42 | cell wall integrity |
| ***SUN41*** | 0.20 | 0.48 | 0.34 | cell wall integrity |
| ***TOS1*** | 0.25 | 0.50 | -0.68 | |
| ***MNN15*** | 0.34 | 0.53 | 0.39 | cell wall integrity |
| ***orf19.6079*** | 0.39 | 0.55 | 0.30 | |
| ***FAD3*** | 0.37 | 0.57 | 0.43 | lipid homeostasis |
| ***ZSP12*** | 0.18 | 0.57 | 0.90 | transport |

**Supplementary Figure 8.**

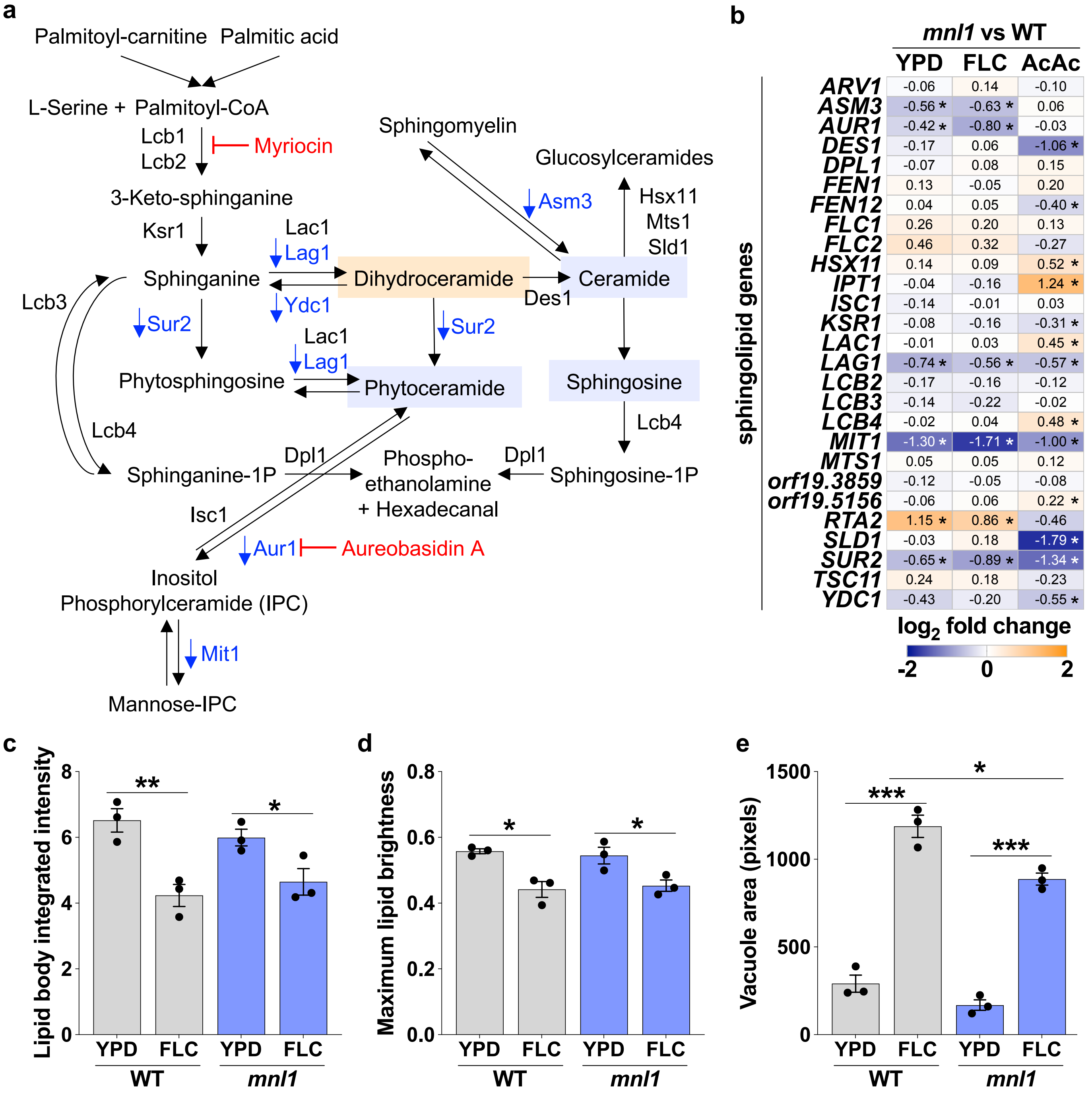

**Supplementary Figure 9.**

**a**

| | WT | *mnl1* |
|---|---|---|
| Chr1 | 1.98 | 1.98 |
| Chr2 | 2.00 | 1.98 |
| Chr3 | 2.04 | 2.04 |
| Chr4 | 2.00 | 1.98 |
| Chr5 | 1.98 | 1.98 |
| Chr6 | 2.04 | 2.00 |
| Chr7 | 2.08 | 2.10 |
| ChrR | 2.00 | 2.00 |
| ChrM | 27.14 | 34.12 |

Chr copy number

10 20 30

**b**

| | | *mnl1* vs WT | | |
|---|---|---|---|---|
| | | YPD | FLC | AcAc |
| mitochondria | *AIF1* | -0.59 * | -0.41 * | 1.25 * |
| | *AUT7* | -0.20 | -0.42 * | -1.04 * |
| | *CBF1* | -0.26 | -0.61 * | -0.98 * |
| | *JSN1* | -0.51 * | -0.63 * | -0.41 * |
| | *MRF1* | -0.41 * | -0.25 | 0.14 |
| | *PTC6* | -0.33 * | -0.25 | -0.44 * |
| PKA pathway | *BCY1* | -0.08 | -0.05 | -0.28 * |
| | *CYR1* | -0.06 | 0.07 | 0.10 |
| | *GPA2* | -0.01 | -0.06 | 0.38 * |
| | *GPR1* | -0.07 | -0.10 | 0.25 * |
| | *PDE2* | 0.07 | 0.21 | 0.84 * |
| | *RAS1* | 0.19 | 0.19 | 0.70 * |
| | *TPK1* | 0.11 | -0.10 | 0.11 |
| | *TPK2* | -0.18 | 0.05 | -0.57 * |

$\log_2$ fold change

-3 0 3

**c**

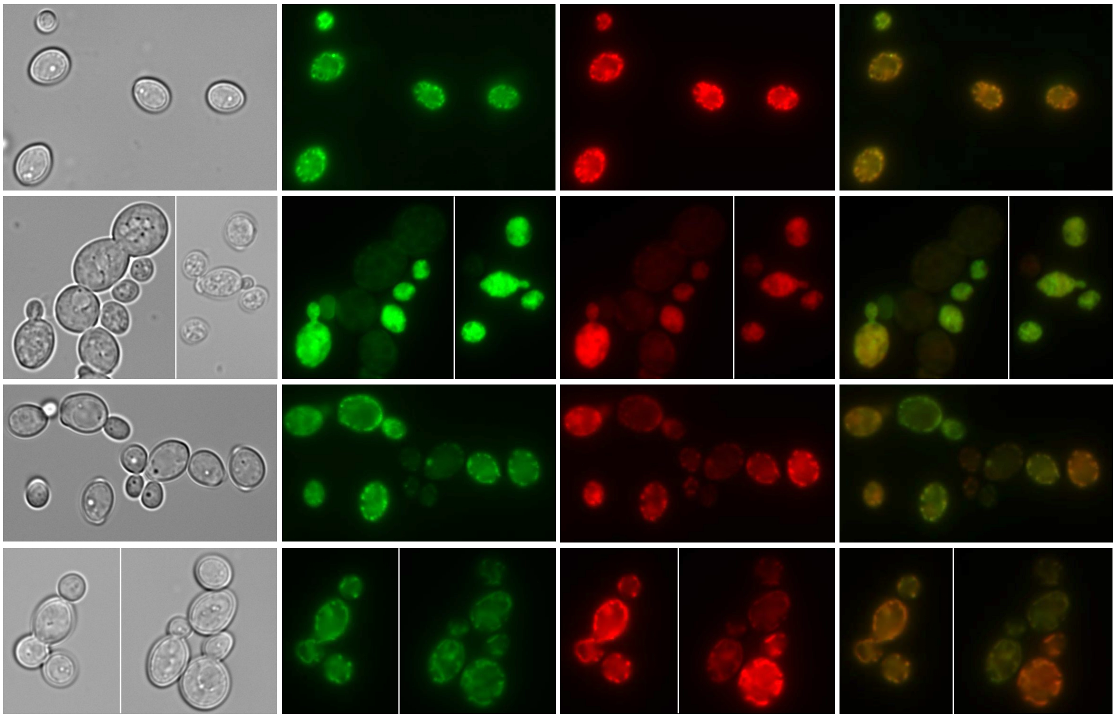


**d**

| | | *mnl1* vs WT | | |
|---|---|---|---|---|
| | | YPD | FLC | AcAc |
| PKA-regulated | *ASR1* | -0.53 | -0.89 * | 0.99 * |
| | *ASR2* | -0.29 | -0.43 * | 0.86 * |
| | *CCW14* | 0.18 | -2.20 | -1.64 |
| | *FAV2* | -0.54 | -0.48 | 0.95 * |
| | *MEP2* | -1.09 | -0.25 | -1.14 |
| | *orf19.4706* | 0.83 | 0.13 | -1.12 * |
| | *orf19.5070* | -0.07 | -0.48 * | 1.12 * |
| ATP | *ATP6* | 2.04 | -0.16 | 0.44 |
| | *ATP8* | 1.97 | 0.31 | 0.95 |
| | *ATP9* | 1.82 | 0.31 | 1.32 |
| | *CDC6* | 0.58 | -0.70 | 0.29 |
| | *MNN42* | -0.56 * | -0.41 | 0.30 * |
| | *orf19.3219* | 0.68 * | 0.69 * | 1.09 * |
| | *PMA1* | 0.45 * | 0.26 | 0.33 * |
| | *PMC1* | 0.61 * | 0.37 | -0.43 * |
| | *PMS1* | 0.52 | 0.01 | -0.62 * |

$\log_2$ fold change

-3 0 3

Supplementary Figure 10.

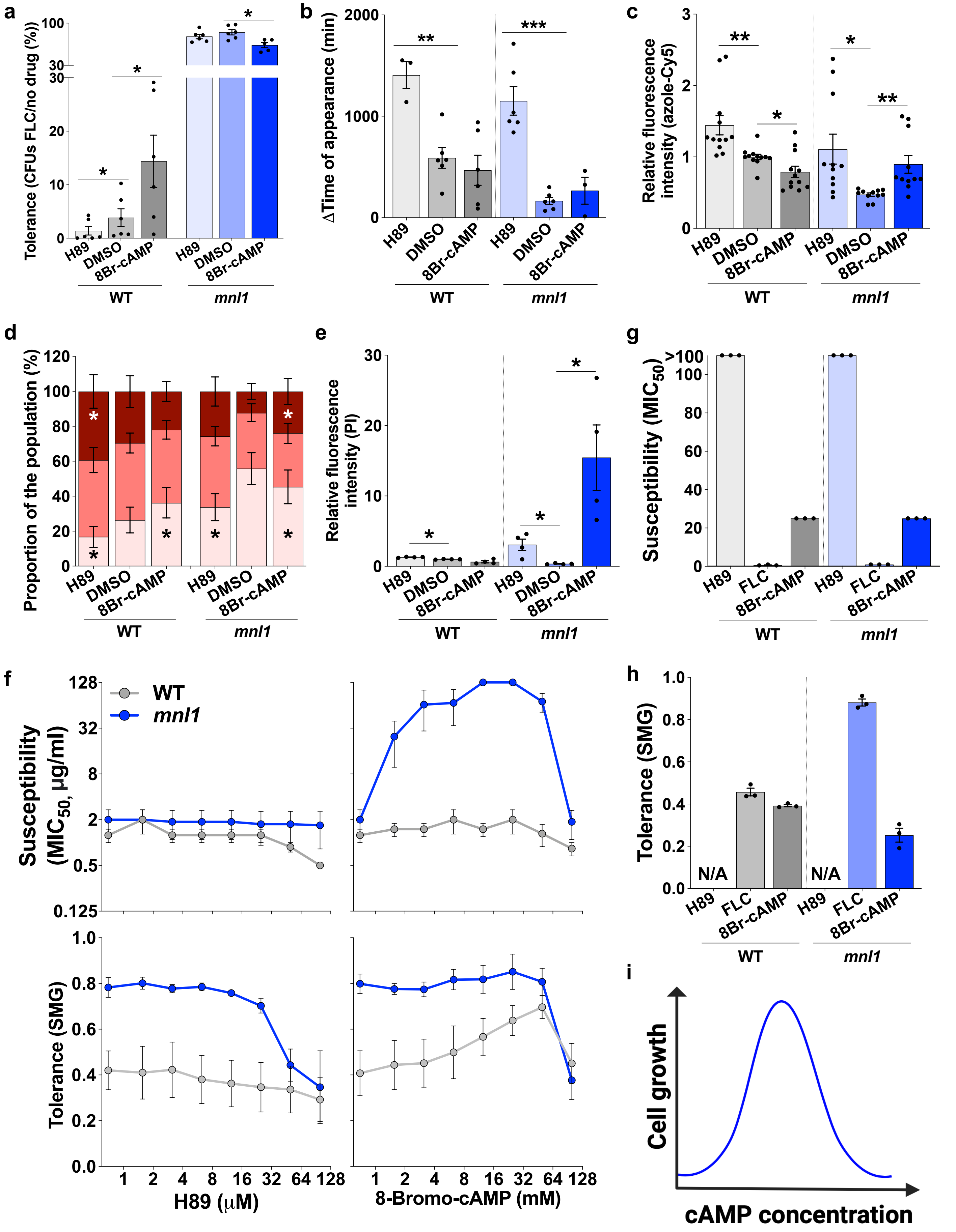

### Supplementary Figure 11.

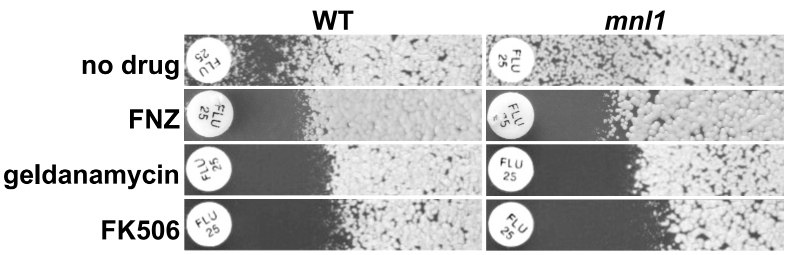


| | RAD$_{20}$ WT | RAD$_{20}$ *mnl1* | FoG$_{20}$ WT | FoG$_{20}$ *mnl1* |
|---|---|---|---|---|
| no drug | 1.00 | 1.02 | 1.00 | 2.05 * |
| FNZ | 1.02 | 1.04 | 0.35 * | 0.30 * |
| geldanamycin | 1.28 | 1.34 | 0.35 * | 0.47 * |
| FK506 | 1.41 * | 1.47 * | 0.44 * | 0.49 * |

**Fold change relative to WT**

0.5 1.0 1.5 2.0

**Supplementary Figure 12.**

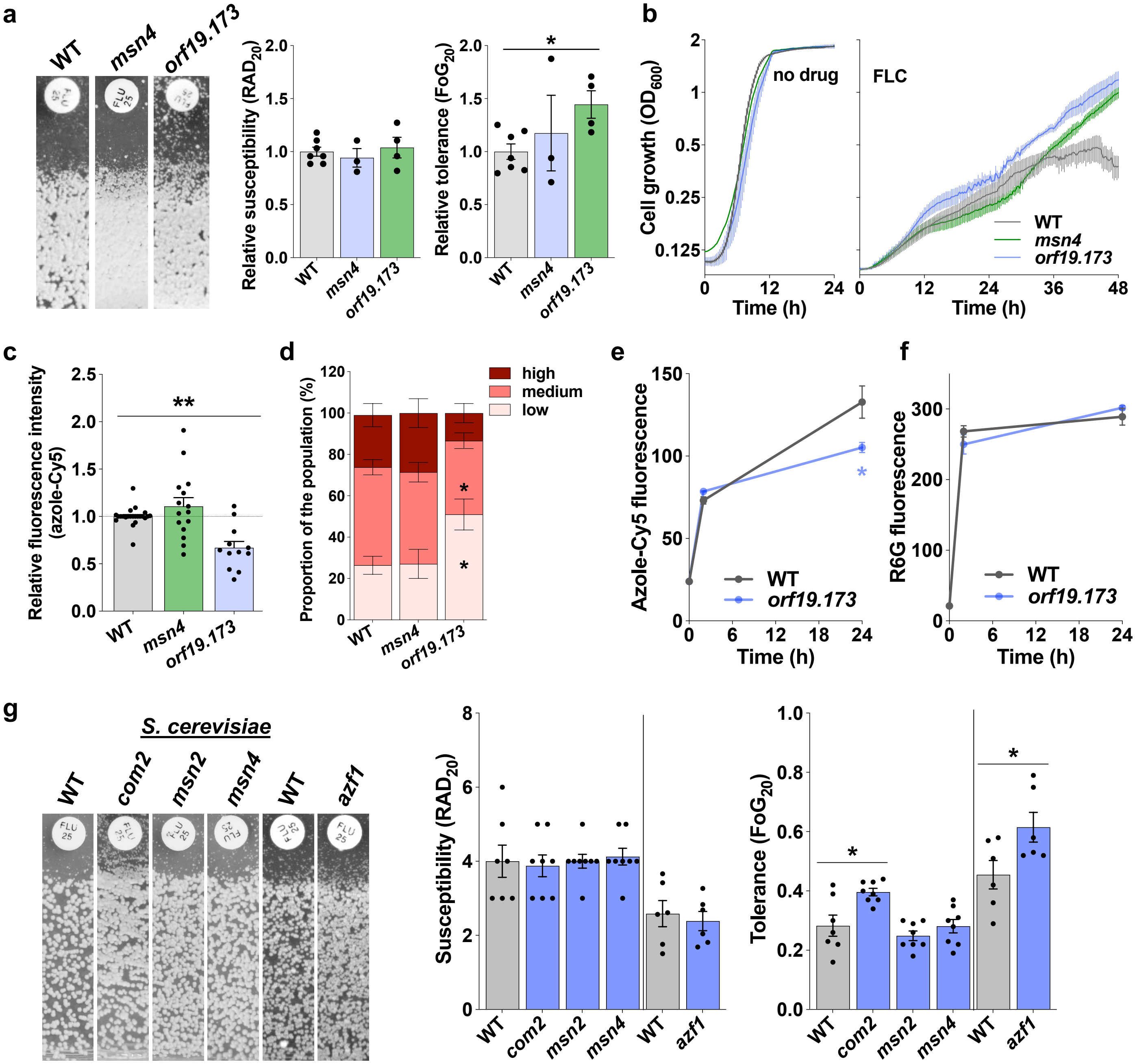

### Supplementary Figure 13.

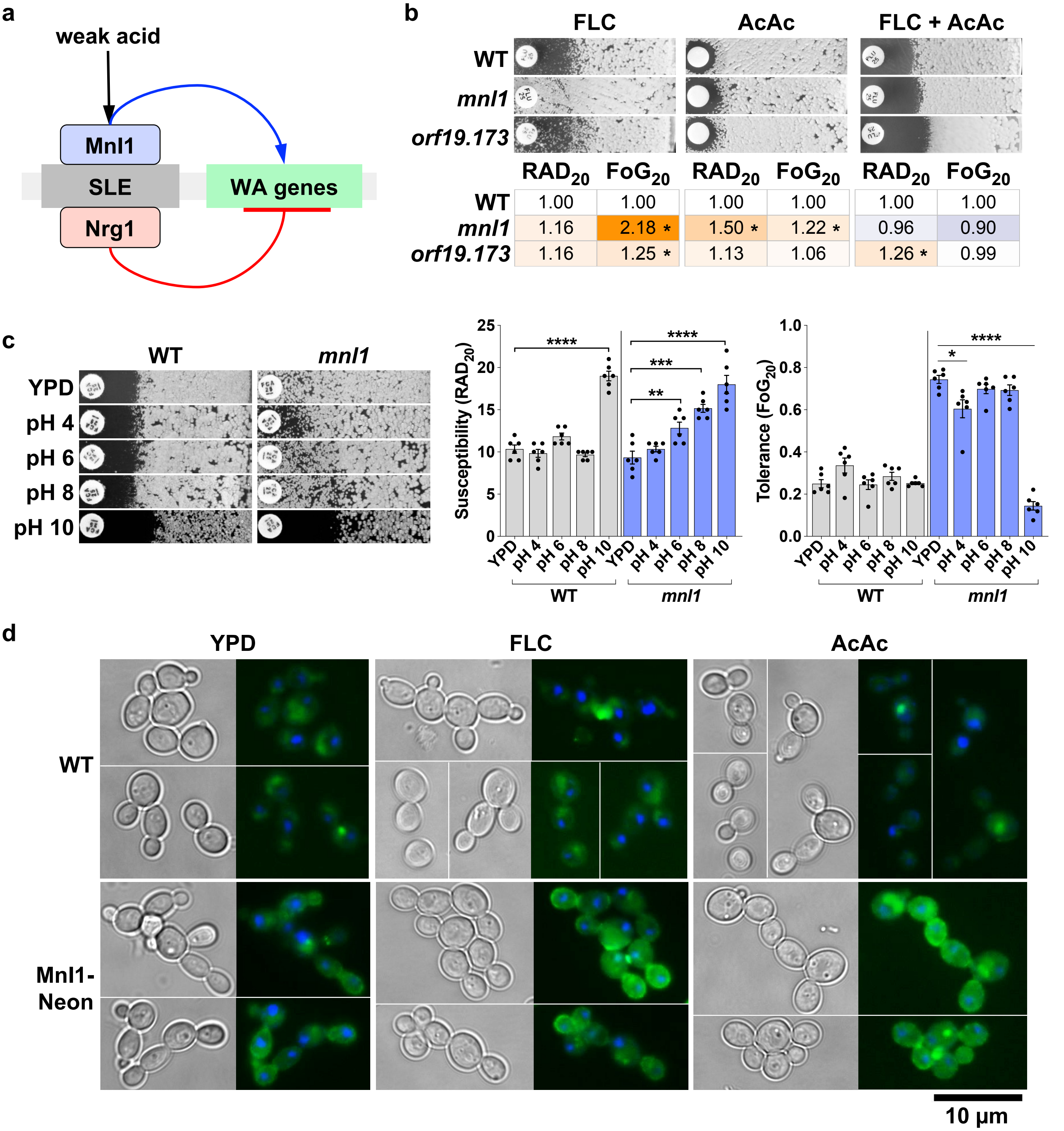


| | FLC RAD20 | FLC FoG20 | AcAc RAD20 | AcAc FoG20 | FLC + AcAc RAD20 | FLC + AcAc FoG20 |
|---|---|---|---|---|---|---|
| WT | 1.00 | 1.00 | 1.00 | 1.00 | 1.00 | 1.00 |
| *mnl1* | 1.16 | 2.18 * | 1.50 * | 1.22 * | 0.96 | 0.90 |
| *orf19.173* | 1.16 | 1.25 * | 1.13 | 1.06 | 1.26 * | 0.99 |

**Supplementary Figure 14.**

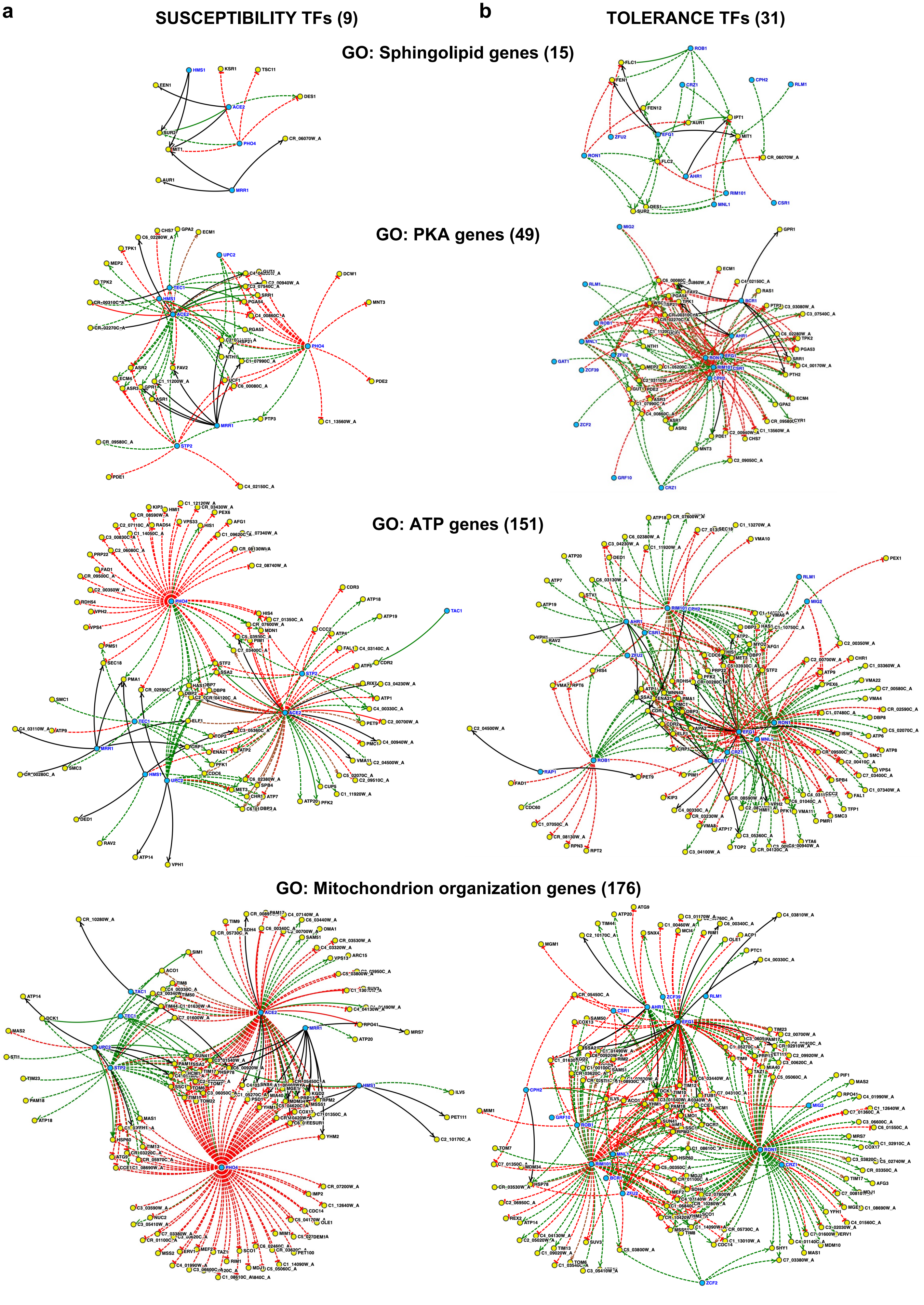

**Supplementary Figure 15.**

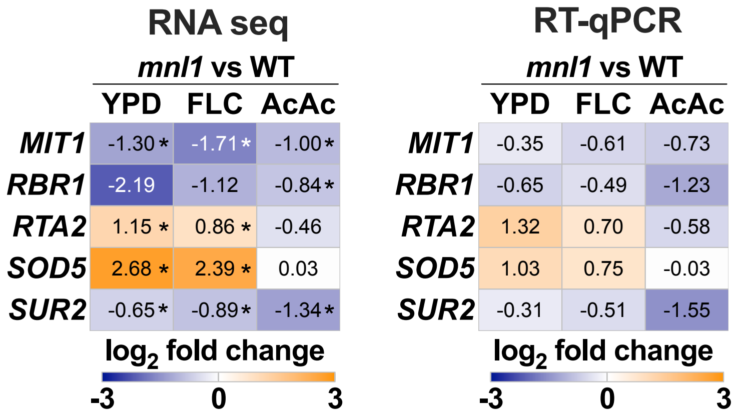

**Supplementary Figure 16.**

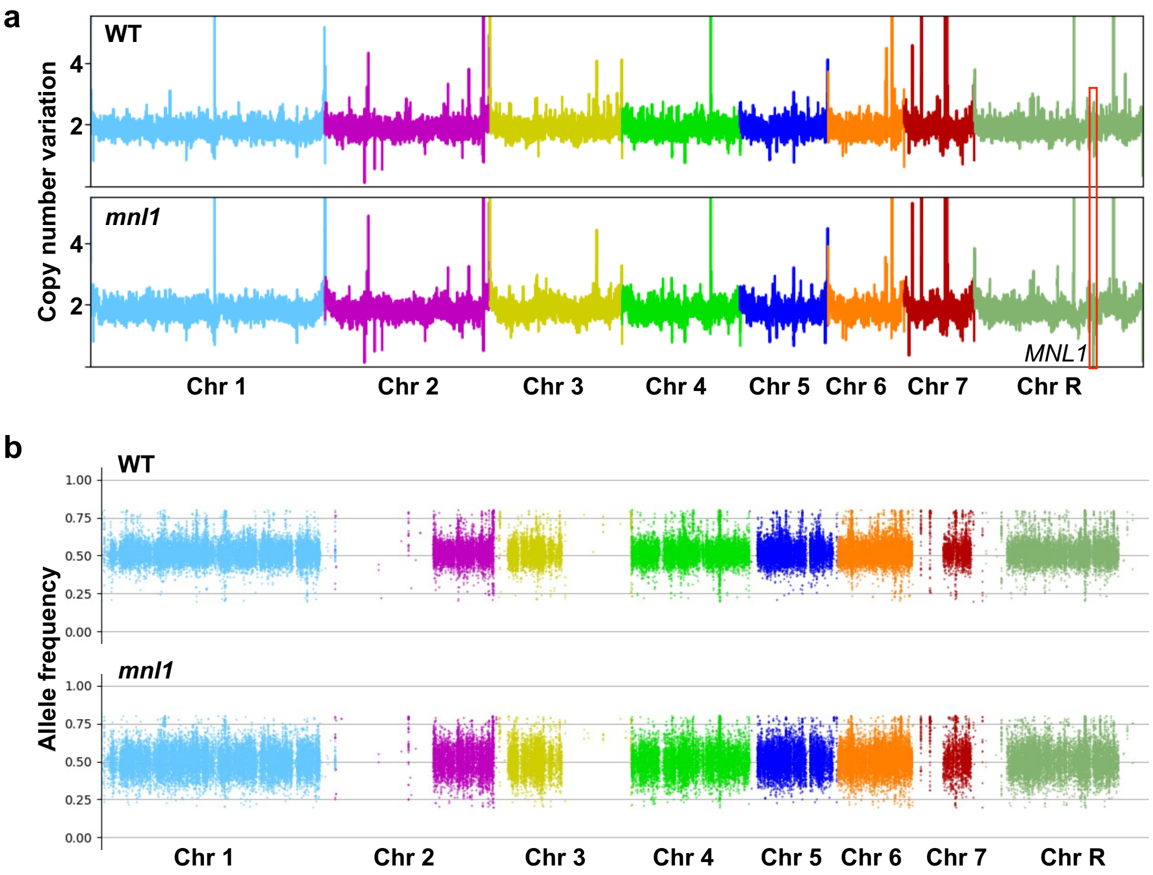
